# Mouse Single Islet β Cell Transcriptomics Reveal Sexually Dimorphic Transcriptomes and Type 2 Diabetes Genes

**DOI:** 10.1101/2020.09.22.307421

**Authors:** Gang Liu, Yana Li, Tengjiao Zhang, Mushan Li, Sheng Li, Qing He, Shuxin Liu, Minglu Xu, Tinghui Xiao, Zhen Shao, Weiyang Shi, Weida Li

**Affiliations:** Translational Medical Center for Stem Cell Therapy and Institute for Regenerative Medicine, Shanghai East Hospital, Frontier Science Center for Stem Cell Research, School of Life Sciences and Technology, Tongji University, Shanghai 200092, China; Tsingtao Advanced Research Institute, Tongji University, Tsingtao Shandong 266003, China; CAS Key Laboratory of Computational Biology, Shanghai Institute of Nutrition and Health, University of Chinese Academy of Sciences, Chinese Academy of Sciences, Shanghai 200031, China; University of Chinese Academy of Sciences, Beijing 100049, China; Department of Statistics, The Pennsylvania State University, University Park, Pennsylvania 16802, USA; Ministry of Education Key Laboratory of Marine Genetics and Breeding, College of Marine Life Sciences, Ocean University of China, Tsingtao Shandong 266003, China

**Keywords:** Type 2 diabetes mellitus, Pancreatic β cell, Sex-biased gene expression, Sex-dependent T2D altered genes, Precision medicine

## Abstract

Type 2 diabetes (T2D), characterized by malfunction of pancreatic β cells, is affected by multiple cues including sex differences. Nevertheless, mechanisms of sex differences in type 2 diabetes susceptibility and pathogenesis remain unclear. Using single-cell RNA sequencing (scRNA-seq) technology, we showed that sexual dimorphism of transcriptome exists in mouse β cells. Our analysis further revealed the existence of sex-dependent type 2 diabetes altered genes in high fat diet induced T2D model, suggesting divergences in pathological mechanisms of type 2 diabetes between sexes. Our results indicated that sex should be taken into consideration when treating diabetes, which was further validated by the sex-matched and sex-mismatched islet transplantation in mice. Compared to sex-matched transplants, sex-mismatched transplants showed downregulation of genes involved in the longevity regulating pathway in β cells and led to impaired glucose tolerance in diabetic mice. Taken together, our findings could advance current understanding of type 2 diabetes pathogenesis with sexually dimorphic perspectives and provide new insights to the development of precision medicine.

## Introduction

The efficacy of current anti-diabetic medication varies significantly among individuals with diabetes, highlighting the importance of personalized treatment for type 2 diabetes. However, the bottleneck for precision medicine lies in the heterogeneous nature of the disease, which not only hinges on genetic predispositions that have been identified by genome wide association study (GWAS), but also other cues, including sex, diet, and aging. Among these factors, sex differences should be first considered for personalized therapy since sex is one of the most recognizable traits.

However, in most current studies on metabolism using rodents, female animals are usually neglected, because male animals have the tendency to show better disease phenotypes and are dominantly used [1]. And this experimental bias on sex hampered novel and comprehensive acknowledgement of metabolic pathological mechanisms. Thus, the National Institutes of Health (NIH) demands sex differences should be emphasized in preclinical studies [2, 3], which should be especially stressed in T2D as a global pandemic.

Glucose homeostasis is controlled by pancreatic islet, which is mainly composed of α cells, β cells, δ cells and PP cells. α cells elevate glucose level by secreting glucagon to promote hepatic glucose synthesis, and β cells release insulin to decrease glucose level by stimulating blood glucose uptake by fat, muscle, liver, and intestine cells, etc. δ cells secret somatostatin to downregulate hormones releasing from both α cells and β cells via paracrine signaling [4]. Polypeptide from PP cells, the most infrequent islet cell type, has effects on both gastric and pancreatic secretions [5].

Despite the similar cell type components of islet architecture shared by both male and female, there exist profound sex differences in islet physiological function and metabolism, signaling pathways involved in hormone releasing, and diabetes occurrence [6]. For example, female rats are more susceptible than males to streptozotocin (STZ) induced diabetes, suggesting that female rat β cells are more sensitive to STZ toxicity than male β cells [6]. Similarly, maternal high fat diet results in insulin resistance and oxidative stress-induced β cell loss specifically in male offsprings, rather than in female offsprings, indicating that female islets might have self-protective machinery against oxidative stress [7]. In human, type 2 diabetes occurs more frequently in men with younger age and less BMI than women [8–10]. Insulin sexual dimorphism of DNA methylation was also observed in human islets by whole islet genome-wide DNA methylation sequencing. It is suggested that sex differences of methylome are associated with differences of islet genes expression and insulin secretion level [11]. However, sexual dimorphism of pancreatic islet β cell has not been investigated at the single cell level, and these studies could reveal sex differences of gene expression in islet β cells between male and female and provide better treatment plans for diabetic patients from both sex groups.

Sex differences in diabetes susceptibility, development and progression have been previously reported, suggesting the existence of sex-dependent diabetes associated genes. Previous studies showed that androgen receptor specifically expressed in male islet β cells and plays an important role in regulating glucose-stimulated insulin secretion in both mice and humans [12]. It has also been reported that *KLF14* allele variants show increased female-specific T2D risk, probably via female-specific fat storage and distribution [13]. Recent banding studies reported that single-cell RNA sequencing technology has been applied to identify novel diabetes altered genes [14–16]. Nevertheless, the diverse sex-dependent diabetes associated genes and molecular pathways have not been comprehensively investigated from these studies.

Here, we systematically analyzed the single cell gene expression profiles of healthy and diabetic β cells from mice. We found a considerable number of genes had sex-biased expression in β cells. Furthermore, we identified 122 sex-dependent diabetes altered genes, suggesting that the molecular mechanisms mediating diabetes pathogenesis in males and females have important differences. Based on the recognition of the sex differences in T2D altered genes and pathways in β cells, we concluded that sex as a biological variance should be emphasized in diabetes treatment. And this conclusion was further supported by the sex-matched and sex-mismatched islet transplantation in mice. Compared to sex-matched transplants, we found that genes involved in the longevity regulating pathway tended to be down-regulated in β cells of sex-mismatched transplants, and glucose tolerance notably decreased in diabetic mice transplanted with sex-mismatched islets. Together, our results not only advanced current understanding of T2D pathogenesis, but also provided new insights and targets for developing sex-dependent precision medicine to treat diabetes.

## Results

### Identification of male and female pancreatic β cell transcriptomes in mouse

To obtain expression profiles of mouse pancreatic β cells, we employed flow cytometry to isolate single islet cells from dissociated mouse islets with live dye staining (**Figure 1A**). Totally, we collected and sequenced 5472 islet cells (3264 male; 2208 female) from both male and female mice, including 1056 islet cells (1056 male; 768 female) of 8-week-old healthy mice, 2208 islet cells (1152 male; 1056 female) of 9-month-old healthy and diabetic mice, 672 male transplanted islet cells (9 months post-transplant) in kidney capsules of both male and female recipient mice, and 768 endogenous pancreatic islet cells (384 male; 384 female) of recipient mice (detailed sample information in **Table 1**). Single-cell RNA sequencing library was constructed by a modified Smart-seq2 protocol. The average library size is 370K reads per cell, and the average number of genes detected in these cells is about 1500 per cell. We retained 4662 cells that have at least 500 genes detected (Figure S1A). The retained cells had high total counts of unique molecular identifiers (UMIs) mapped to gene exons (Figure S1B), and low fractions of UMI counts of mitochondrial genes (Figure S1C), suggesting we obtained a set of qualified scRNA-seq profiles of mouse islet for the main purpose of this study.

**Figure 1.**
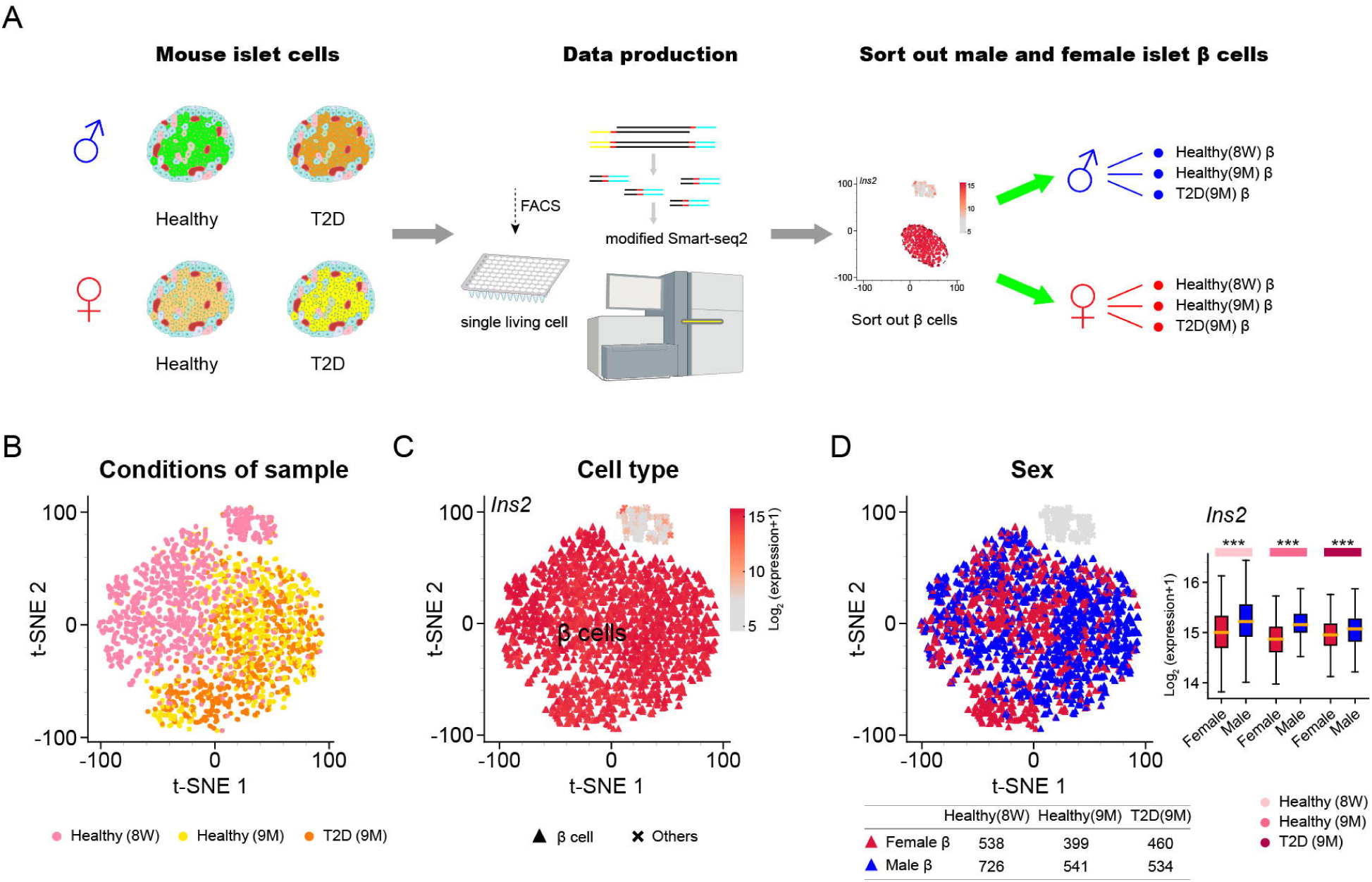
Single β cell transcriptomes reveal sexually dimorphic genes in healthy and T2D mice. **A.** Schematic diagram for mouse single β cell RNA-seq data preparation and analysis. Mouse single living islet cells are labelled by Calcein Blue, AM, and collected through Fluorescence activated cell sorting (FACS). β cells are sorted out and divided into different groups according to donor conditions. 8W, 8-week-old; 9M, 9-month-old. **B.** t-SNE map of cells from mouse in different conditions. The cells from healthy and T2D mice are colored with different red-colored items. **C.** Mouse β cells are identified by highly expressed marker gene *Ins2*. **D.** Mouse β cells are colored by sex information. And the expression of β cell marker *Ins2* is significantly different between cells of males and females (K-S test) as shown in the box plot.

**Table 1.** The number of collected cells from mice in different conditions.

From these 4662 cells, we went on to identify β cells for downstream analysis. Firstly, we applied an adjusted CPM method (adjCPM) to normalize our scRNA-seq data by excluding the union of the top 2 genes with the highest expression from all cells while calculating the normalization factors (Figure S1D and S1E; see Methods). Secondly, hierarchal clustering of scRNA-seq profiles based on the union of the top 10 highly expressed genes from all cells was used to discriminate the identity of the cells (Figure S1F). After performing principal component analysis with the identified hyper variable genes (HVGs) and subsequent visualization by t-distributed stochastic neighbor embedding (t-SNE) (Figure S1G; see Methods), the retained islet cells of healthy (8-week-old; 9-month-old) and diabetic mice were found to be aggregated into two clusters exhibiting differential expression of β cell marker gene *Ins2* (**Figure 1B and 1C**). Then, 3912 β cells (2197 male; 1715 female) from mice in different conditions were identified through high expression of *Ins2* (Figure 1C and S4A). The sex of mouse β cells was labeled on the t-SNE map (male in blue; female in red; **Figure 1D**), and further confirmed by X and Y chromosome genes (Figure S2B). We found male and female β cells were not completely overlapped on the t-SNE plot (Figure 1D). To verify the existence of sexually dimorphic gene expression, we applied Kolmogorov-Smirnov test (K-S test) to analyze the differential expression of *Ins2* between male and female β cells, and found its expression was significantly different between two sexes (Figure 1D), indicating sexual dimorphism of β cell transcriptomes exist in mouse and could be important to β cell functions.

### Sex-biased gene expressions in mouse β cells under healthy and T2D conditions

To identify genes that display sex-biased expression pattern in mouse pancreatic β cells, we first compared the transcriptional profiles of male β cells with female β cells from 8-week-old C57BL/6J mice (**Figure 2A**). Differentially expressed genes were sorted out by MAST [17], with a FDR cutoff of 0.05 (the same cutoffs were used for all the differential analysis using MAST in our study). In total, we obtained 162 differentially expressed genes (DEGs), comprising 37 genes expressed higher in male and 125 genes expressed higher in female (Figure 2A). Among them, only four genes were on the sex chromosomes, including X chromosome (*Xist*) and Y chromosome (*Eif2s3y*, *Ddx3y*, *Uty*) (**Figure 2B**), suggesting that considerable level of sexual dimorphism exist in healthy β cell transcriptomes.

**Figure 2.**
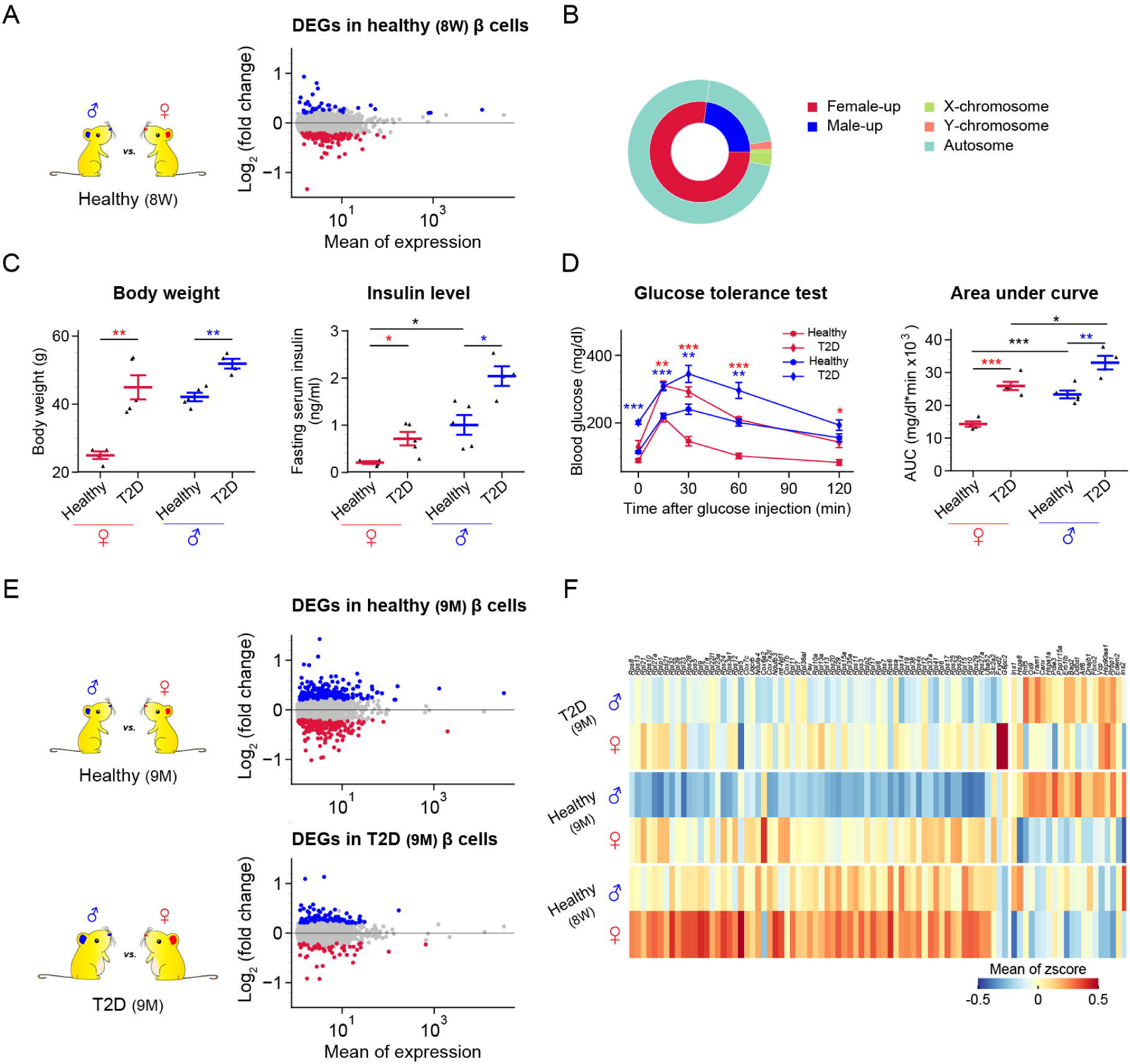
Single β cell transcriptomes reveal sex-biased genes in healthy and T2D mice. **A.** Comparison between β cells of male and female 8W (8-week-old) C57 mice. In the MA plot, the sex-biased genes are highlighted by blue (male-biased) and red (female-biased) dots. See also Table S1. **B.** Nested pie chart depicting genomic location of sex-biased genes. **C.** Diabetes associated physiological phenotypes. Body weight, fasting insulin serum level are detected. The blue line represents the sample of male healthy (n=5 mice) and male T2D (n=4 mice); red line represents the sample of female healthy (n=4 mice) and female T2D (n=5 mice). Results are presented as mean ±SEM; * p ≤ 0.05, ** p ≤ 0.01, two sample t-test. **D.** Intraperitoneal injection glucose tolerance test with area under curve (AUC). Results are presented as mean±SEM;* p ≤ 0.05, ** p ≤ 0.01; *** p ≤ 0.001. Red *, **, *** represent female T2D versus female healthy mice; blue *, **, *** represent male T2D versus male healthy mice, two sample t-test. **E.** Comparisons between male and female β cells of 9M (9-month-old) healthy and T2D mice. In the MA plots, sex-biased genes are highlighted by blue (male-biased) and red (female-biased) dots. See also Table S2 and S3. **F.** Heatmap of genes selected from sex-biased genes that overlapped with the leading-edge genes of GSEA. See also Table S4.

To investigate whether such sexually dimorphic β cell transcriptomes also exist in diabetic conditions, β cells from high-fat-diet (HFD) induced diabetic mice were collected for scRNA-seq. Firstly, we used HFD to induce T2D model with β cell failure in C57BL/6J mice (fed with HFD from 8-week-old) as previously described [18]. Consequently, characteristic T2D phenotypes were observed in those diabetic mice of both male and female, including high body weight, increased fasting insulin level, and impaired glucose tolerance (**Figure 2C and 2D**). Notably, sex differences of fasting serum insulin level in healthy mice were also observed: the fasting serum insulin level of healthy male mice is significantly higher than healthy female mice (Figure 2C). In addition, the area under curve (AUC) of glucose tolerance test showed that the sex differences of glucose tolerance exist between male and female in healthy and T2D mice (Figure 2D). Then, to explore the genetic basis for such differences, we obtained single β cell transcriptomes from 9-month-old diabetic mice (HFD feeding up to 7 months) and age-matched healthy mice (normal diet, ND) of both sexes. The comparison of single β cell transcriptional profiles of male and female from either healthy or T2D mice was carried out by MAST with the same cutoffs as described above. As shown in the results, 394 DEGs between male and female in 9-month-old healthy mice were identified, including 200 genes expressed higher in male and 194 genes expressed higher in female. In parallel, 81 male highly expressed genes, and 52 female highly expressed genes were identified in β cells from T2D mice (**Figure 2E**).

To further elucidate sex-biased pathways based on these comparisons, gene set enrichment analysis (GSEA) was performed with the cutoff set as FDR <= 0.25 (the same cutoff was used for all GSEA in our study) [19]. In β cells from both 8-week-old and 9-month-old healthy animals, the longevity regulating pathway was enriched in males, and related genes (*Ins1*, *Ins2*, *Hspa1a*, *Hspa8*) were expressed significantly higher in males (**Figure 2F**; S2C and S2D). Intriguingly, the ribosome pathway was consistently enriched in females in β cells from both healthy (8-week-old; 9-month-old) and diabetic mice (Figure S2C; S2D and S2E). Importantly, in β cells from diabetic mice, both the N-Glycan biosynthesis pathway and Notch signaling pathway were enriched in males. Conversely, the JAK-STAT signaling pathway, ferroptosis pathway, spliceosome pathway, carbohydrate digestion and absorption pathway were enriched in female β cells from diabetic mice (Figure S2E). Expression patterns of representative sex-biased DEGs involved in the GSEA results were shown in the heatmap (Figure 2F). Above all, our results validated the existence of sex-biased gene expression in β cells of both healthy and diabetic mice, indicating that the pathological mechanism of T2D might differ between males and females.

### Abundant sex-dependent T2D altered genes were found in mouse β cells

Given the sex-biased gene expressions detected in T2D β cells, we hypothesized T2D development may differ in males and females. Previous studies compared the single-cell transcriptional profiles of T2D islets with that of healthy islets to identify T2D altered genes [14–16] without considering the factor of sex. To dissect that in a sex-dependent manner, we compared the single-cell transcriptome of β cells from T2D mice with that from age-matched healthy mice. Firstly, we performed differential analysis between T2D and healthy group with scRNA-seq profiles of all beta cells from both sexes using MAST (here both cellular detection rate (CDR) and sex were provided as covariates) and defined the DEGs identified as sex-independent T2D altered genes

Finally, we identified 98 sex-independent (40 downregulated, 58 upregulated) T2D altered genes (Table S5). Then, we compared the diabetic β cells with healthy β cells of the same sex to obtain DEGs in either male or female specific manner, and consequently compared the DEGs of each sex with the 98 sex-independent genes. The non-overlapping parts were defined as candidate genes for sex-dependent T2D altered genes, including 56 candidate genes in female and 65 candidate genes in male (**Figure 3A**). Based on the defined candidate genes, we introduced two additional filtering criteria to define sex-dependent T2D altered genes in a more reliable way: 1. The estimated 95% confidence interval of log2 fold change (log2FC) by MAST from the differential expression analysis in one sex does not overlap with that from the other sex. 2. The p-value in the non-significant sex group is higher than 0.2. We finally used the genes passing both of these two additional criteria as the final sex-dependent T2D altered genes. As the result, we obtained 31 female-specific (22 up and 9 down regulated in T2D compared to Healthy) and 31 male-specific (18 up and 13 down regulated in T2D compared to Healthy) T2D altered genes (**Figure 3B** and Table S6). Moreover, 14 selected sex-dependent T2D altered genes with basemean > 10 were validated by q-PCR experiments (Figure S3A).

**Figure 3.**
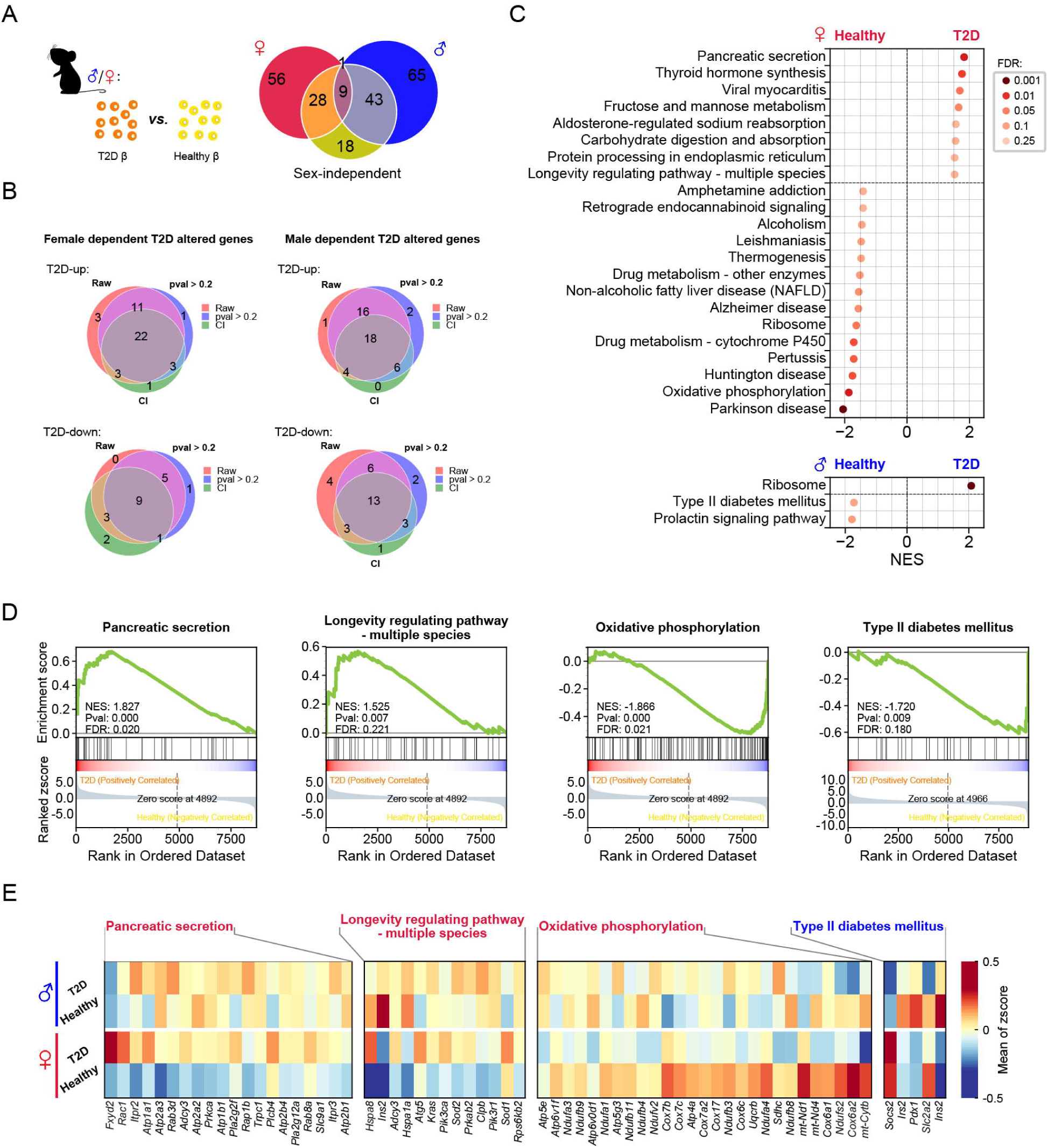
Single β cell transcriptomes reveal sex-dependent T2D altered genes and pathways in β cells. **A.** Venn diagram depicting the definition of candidate genes for sex-dependent T2D altered genes. **B.** Venn diagram show the definition of sex-dependent T2D altered genes. The genes in the overlapped part of venn diagram are sex-dependent T2D altered genes. “Raw” represents the candidate genes defined in Figure 3A. “CI” represents the genes that confidence interval of log2FC from analysis of one sex does not overlap with that from the other sex. “p > 0.2” represents the p-values in the non-significant sex group are high. See also Table S6. **C.** Results of GSEA showing pathways significantly enriched in the healthy group and the T2D group in male (labeled in blue) and female (labeled in red). Pathways with NES > 0 are enriched in T2D β cells and pathways with NES < 0 are enriched in healthy β cells. **D.** GSEA plots of pathways involved in the onset of diabetes or the function of β cell. **E.** Heatmap of leading-edge genes of the selected pathways.

To note, among these defined sex-dependent T2D altered genes of female, we found upregulated expressions of *Serp1* and *Ero1lb*, which encode endoplasmic reticulum (ER) stress associated proteins. Overexpression of *Ero1lb* in β cells results in the upregulation of unfolding protein associated genes, and leads to ER stress [20]. Interestingly, intervention of conjugated estrogens could protect postmenopausal women from diabetes by promote degradation of misfolded proteins during the process of insulin synthesis in β cells [21]. Taken together, these findings indicate sex differences of ER stress might exist in T2D pathogenesis.

Among male-dependent T2D altered genes, several genes have been reported to confer sex specific effect in physiological function of β cells or onset of T2D. Notably, our analysis identified *Iapp* as male dependent T2D upregulated gene. Accumulation of IAPP is toxic to pancreatic β cells. It was reported that ectopic expression of human *IAPP* in pancreatic β cells of transgenic mice resulted in more severe diabetic phenotype in males than in females, suggesting sex dependent pathogenesis [22], which is in line with our result. Moreover, insulin secretion capacity associated with mitochondrial function declines with aging which is a contributing factor to T2D. Evidence showed that elderly male Wistar rats show more significantly decreased mitochondrial function and insulin secretion than elderly females [23], consistent with our results that two mitochondrial function related genes, *Cox4i1* and *Ndufb7,* were identified as downregulated male dependent T2D altered genes.

For other sex-dependent T2D altered genes, there was no previous report on sex differences of them in diabetes pathogenesis, such as *Cpe*, *Scg3*, *Ttr*, *Rnase4*, *Malat1*, *Ucn3*, *Pura*, *Tmed3*. Because in most current studies on metabolism with rodent models, there is a experimental bias on dominant using of males which have the tendency to show better disease phenotypes [24], limiting understanding of T2D pathogenesis with sex specific perspective. Thus, it will be valuable to examine sex specific role of these genes in T2D pathogenesis in future study.

To further systematically investigate the sex differences in T2D altered pathways, GSEA was performed based on comparison of β cells from T2D and healthy mice of each sex. In females, several pathways enriched in T2D-β cells such as pancreatic secretion pathway and longevity regulating pathway (multiple species) have direct links to β cell function and onset of T2D. Conversely, oxidative phosphorylation pathway and ribosome pathway were enriched in healthy β cells. At the same time, in males, T2D-β cells had ribosome pathway enriched and healthy β cells had type 2 diabetes mellitus pathway enriched (**Figure 3C and 3D**). The expression patterns of genes involved in the pathways mentioned above were shown in the heatmap (**Figure 3E**) and violin plots (Figure S3B). Altogether, both the distinctive patterns of sex-dependent T2D altered genes and different T2D associated pathways suggested divergence of T2D pathogenesis between sexes.

Although we detected that no gene set showed significant enrichment in both sexes, we found many of the sex-independent DE genes (98 in total, 40 downregulated, 58 upregulated) (Table S5) are involved in fundamental β cell function, or T2D pathogenesis. For example, in regards to regulating insulin secretion process, we detected that *Slc2a2*, *Calm2*, and *c-Fos* were downregulated in sex-independent manner. *Slc2a2* encodes the glucose transporter 2 (Glut-2). Impaired insulin secretion is a key manifestation of T2D. Glut-2 functions as a membrane protein to transport glucose into β cells to mediate glucose-stimulated insulin secretion (GSIS) which is crucial for glucose homeostasis [25]. Evidence showed that mutations of *Slc2a2* result in neonatal diabetes in both male and female patients [26]. It was also reported that *Slc2a2* pathological variants are associated with increased risk of transition from impaired glucose tolerance to T2D [27]. Moreover, *Calm2*, encodes Camodule-2 to mediate calcium signaling in GSIS [28]. At the same time, overexpression of *c-Fos* in β cells was reported to contribute to increased insulin secretion, whereas its knock-down results in decreased insulin secretion [29].

Meanwhile, we also observed sex-independent upregulated genes involved in critical cellular events in T2D pathogenesis, such as transdifferentiation and ER stress. As reported previously, β cell to alpha cell (glucagon producing) transdifferentiation occurs in T2D pathogenesis [30, 31]. Consistently, *Gcg* (encoding glucagon) was identified as upregulated sex-independent T2D gene in our analysis. Moreover, *Hsp90aa1* was also upregulated in both male and female T2D β cells. Previous study identified Hsp90 (encoded by *Hsp90aa1*) as a therapeutic target for ER stress in T2D, and inhibition of Hsp90 alleviates insulin resistance and improves glucose tolerance [32, 33]. Collectively, these results suggest not only divergence of pathological mechanism but also common pathogenesis in T2D between sexes.

### Mice with sex-matched **β** cell transplant exhibited better control of glucose homeostasis

Altogether, the existence of sexual dimorphism in mouse β cell transcriptome informed us that sex as an important factor in β cell function and pathogenesis of T2D should be emphasized when treating diabetes. In order to validate this critical concept, long-term islet transplantations (an experimental treatment for insulin insufficient diabetes mellitus) with sex-matched and sex-mismatched islets were performed in ICR mice. The ICR mice selected as donor and recipient in this assay for the reason that ICR mouse do not develop spontaneous insulitis and diabetes, therefore they are more suitable for long-term assessment of the effects of islet transplantation than another inbred strain C57BL/6. Male islets were isolated from 6-8 weeks old mice and transplanted into kidney capsules of age-matched male and female mice, respectively. Transplanted islets (TX-islet) and endogenous islets (Endo-islet) from recipient mice were all collected and dissociated for scRNA-seq profiling, 9 months post-transplant (**Figure 4A**). The β cells identified by high expression of *Ins2* as previously described (**Figure 4B**). Differential expression gene analysis in β cells was performed as the following two ways. Firstly, we plotted the gene expression changes observed in comparing the transcriptomes of sex-matched and sex mismatched transplanted β cells to those of male (**Figure 4C** left panel) or female (Figure 4C right panel) endogenous beta cells. In both analysis, we observed a obvious correlation between the gene expression changes detected the two comparisons, which indicates that most of the differentially expressed genes showed consistent expression changes between transplanted and endogenous beta cells in male and female recipients. However, it can also be seen that the distribution of the log2 expression fold changes shown on x-axis was clearly wider than the distribution of those shown on y-axis in both plots, suggesting that the overall sex-matched transplant-β cell transcriptomes were closer to β endogenous islet β cells from both male and female recipients, than the sex-mismatched transplant- β cell transcriptomes. Secondly, transplant- β cell transcriptomes were directly compared between sex-matched and sex-mismatched groups for DEGs. Compared to sex-mismatched group, sex-matched transplant- β cells ad 3 genes (*Kap*, *Sfrp5* and *Akr1c21*) significantly up-regulated and 2 genes (*Ovol2*, *Matn2*) significantly down-regulated (Figure S3C and S3D). Notably, *Sfrp5* was a conservative male-biased expression gene in 8-week-old and 9-month-old healthy β cell (Table S1 and S2), previous study reported that overexpression of *Sfrp5* (down-regulated in obesity and T2D) can ameliorate impaired glucose tolerance in mice [34]. In humans, high level of serum SFRP5 was correlated with lower risk of T2D onset [35]. Moreover, the results of GSEA showed that longevity regulating pathway-multiple species was significantly enriched in β cells of sex-matched transplant-islets and among the 16 leading-edge genes, *Ins2, Sod1, Sod2* and *Foxa2* are directly linked to β cell secretion (**Figure 4D**). Collectively, these results suggested that sex-matched islet transplantation could be more beneficial to the function of β cells of the transplants, thus better controlling glucose homeostasis.

**Figure 4.**
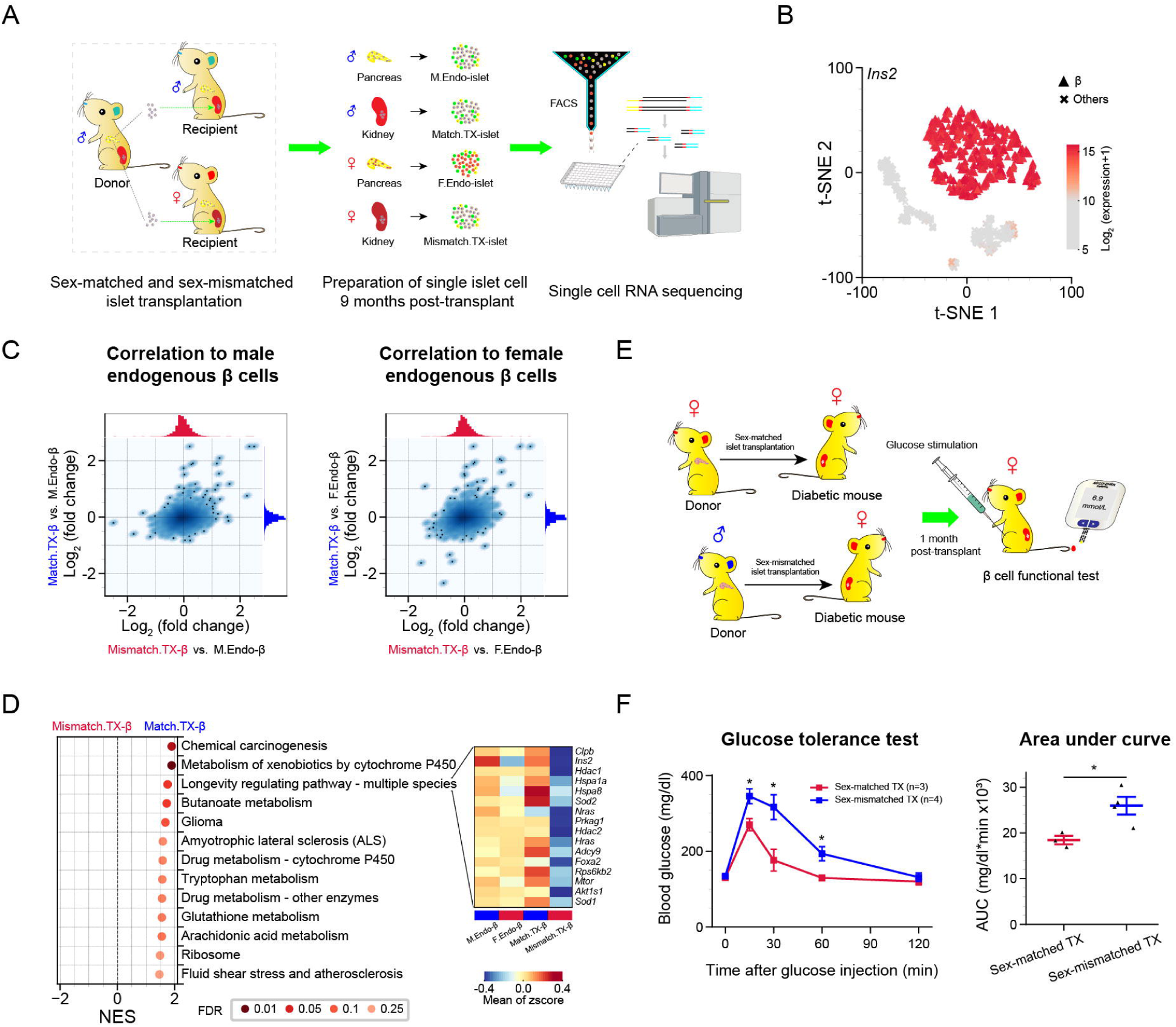
Sex-matched islet transplantation confers better glucose tolerance than sex-mismatched transplantation. **A.** Schematic diagram of sex-matched and sex-mismatched islet transplantation for scRNA-seq. Single islet cells are collected for scRNA-seq 9 months post-transplant. M.Endo-islet, endogenous islets of male recipients; F.Endo-islet, endogenous islets of female recipients; Match.TX-islet, transplanted islets in male recipients; Mismatch.TX-islet, transplanted islets in female recipients. **B.** t-SNE map of endogenous and transplanted islet cells in both sex-matched and sex-mismatched transplantation, and β cells are identified by highly expressed *Ins2*. Details see also Figure S2A. **C.** Scatterplot depicting correlation of Log2(fold change) generated by comparing β cells of TX-β (sex-matched or sex-mismatched) and endo-β. M.Endo-β, endogenous β cells of male recipients; F.Endo-β, endogenous β cells of female recipients; Match.TX-β, sex-matched transplant-β cells in male recipients; Mismatch.TX-β, sex-mismatched transplant-β cells in female recipients. **D.** Pathways enriched in sex-matched transplant-β cells. Leading-edge genes of longevity regulating pathway are zoomed in with heat-map. M.Endo-β, endogenous β cells of male recipients; F.Endo-β, endogenous β cells of female recipients; Match.TX-β, sex-matched transplant-β cells in male recipients; Mismatch.TX-β, sex-mismatched transplant-β cells in female recipients. **E.** Schematic diagram of sex-matched and sex-mismatched islet transplantation for β cell functional test in STZ induced diabetic mouse. **F.** Oral glucose tolerance test (OGTT) for mice after sex-matched (n=3 mice) and sex-mismatched (n=4 mice) islet transplantation and results are summarized by the area under curve (AUC). 2 g/Kg glucose is gavaged after 6 hours fasting, and blood glucose level is detected at 0, 15, 30, 60, 120 minutes after glucose gavage. Results are presented as mean±SEM; * p ≤ 0.05, two sample t-test. TX, transplantation.

To further validate the advantage of sex-matched transplantation, we transplanted islets from male and female mice into streptozotocin induced female diabetic mice, respectively (**Figure 4E**). Although the hyperglycemia (> 350 mg/dl) of the diabetic recipient with sex-matched or sex-mismatched transplantation restored to normal levels 1 month post-transplant (< 144 mg/dl; Figure S3E), the glucose tolerance of the diabetic mice with sex-matched transplantation was significantly better as evidenced by the oral glucose tolerance test (**Figure 4F**). Above all, our scRNA-seq analysis of β cells and experimental validation concluded that sex should be taken into consideration in diabetes treatment.

## Discussion

According to previously published work, it has been well recognized that sexual dimorphism exists in many organs or systems, such as heart, kidney and immune responses [36–38]. What’s more, a recent study in humans has shown that sexual dimorphism not only exhibits in neuron cells, but also is associated with susceptibility of mental diseases [39]. Herein, T2D is a complex metabolic disorder characterized with islet β cell failure and impacted by sex. Sex differences in islet β cell physiological function and diabetes prevalence have been recognized, but still need to be better understood. To decipher the sexually dimorphic T2D pathogenesis, we used scRNA-seq to comprehensively measure the transcriptomes of healthy and diabetic β cells from mice of both sexes. We identified abundant genes have significantly sex-biased expression in β cells of both healthy and T2D mouse.

Besides, we found longevity regulating pathway was male specific enriched in healthy β cells. Evidence has shown that insulin and insulin like growth factor signaling are involved in longevity pathway, suggesting the crucial role of metabolic homeostasis of glucose for aging and life span [40–42]. Expressions of two insulin encoding genes, *Ins1* and *Ins2*, were specifically upregulated in β cells of healthy male mice, suggesting enhanced insulin synthesis. At the same time, we observed another two longevity pathway related genes *Hsp1a* (in β cells of healthy 8-week-old mice) *and Hsp1a* (in β cells of healthy 8-week-old and 9-week-old mice) exhibiting healthy male specific upregulation. Both *Hsp1a* and *Hspa8* were reported to protect β cells from oxidative stress, and upregulation of *Hspa1a* is associated to extended longevity [43, 44], which may suggest a protective mechanism against the stress caused by increased insulin synthesis in healthy male mice. Furthermore, we identified 62 sex-dependent T2D altered genes in mouse, suggesting important differences in the molecular mechanisms of diabetes pathogenesis between males and females in mouse T2D model. Collectively, these results provided innovative sex-specific targets for future studies on the precision treatment of T2D.

In current clinical trials of islet transplantation, sex matching between donor and recipient has not been stressed. Based on the recognition of sex differences in the transcriptome of β cells, we concluded that sex as a crucial biological variance should be emphasized in the treatment of diabetes. And this conclusion was further supported by the sex-matched and sex-mismatched islet transplantation in mice. Compared with the sex-mismatched group, the β cells of transplants in the sex-matched group showed significant enrichment of the longevity regulating pathway (consistent with the results of Figure S2C and S2D) and glucose tolerance notably improved. The long-term curative effect of islet transplantation is impacted by the survival of the transplants [45]. In comparison of sex-matched transplantation and sex-mismatched transplantation, we found that the longevity regulating pathway was also enriched in β cells of sex-matched transplantation with upregulated expressions of genes related to longevity pathways. Among them, *Ins2*, *Hsp1a* and *Hspa8* were also upregulated in the sex-matched group. At same time, upregulated expressions of gene involved in extending longevity of organism were also detected, such as *Sod1*, *Sod2*, *Hdac1* [46–48]. Taken together, our results suggested that sex should be taken into consideration as an important factor for islet transplantation to achieve long-term stability and functionality of the transplants. For better curative effect, our result raised the necessity for sex-matched islet transplantation, and even for sex-matched stem cell-based cell replacement therapy for diabetes treatment.

Beyond the islet β cells focused in this study, sex differences of T2D susceptibility were also associated with sex steroid hormones. It has been found that endogenous estrogens are protective against T2D in females and the risk of T2D increases following menopause [49, 50]. Estrogen not only improves islet β cell function and survival [51], but also stimulates the secretion of GLP-1 by both islet α cells and intestine L cells to maintain glucose homeostasis [52]. Along with that, deficiency of the male sex hormone, testosterone, increases the T2D risk in male [53, 54]. Furthermore, a recent report revealed that testosterone improves insulin secretion through androgen receptor on male islet β cells in both mouse and human [12]. Taken together, sex differences in both islet β cells and sex steroid hormones need to be highlighted in the development of sex-specific precision medicine.

## Materials and methods

### Animals and high fat diet induced diabetic mice

We housed all the mice under the specific pathogen free grade environment of animal facility at Tongji University, Shanghai, China. Adult male and female mice (6-8 weeks old) were purchased from Shanghai Slac Laboratory Animal. To induce mouse T2D model, male and female C57BL/6J mice (n=5 per sex) were fed with high-fat diet from 8-week-old to 9-month-old as a high-fat-diet (HFD) group, and equal number of male and female mice were fed up with normal diet (ND) in an age-matched way to a control potential confounding factor, age.

### Islet transplantation

Islets were isolated from 6-8 weeks old male ICR mice (n=20), and the detailed process of islet isolation was as previous description [55]. Similar size islets were handpicked after purification under a stereomicroscope, and 3 age-matched healthy male and 3 female ICR mice were selected as a recipient with about 300-400 islets were transplanted under the kidney capsule. About 9 months later, transplanted islets were dissected and scraped from the kidney capsule, and the pellet was collected and dissociated into a single cell for scRNA-seq in two batches. At the same time, the endogenous islets of recipient mice were also isolated and dissociated into a single cell for scRNA-seq. Diabetic mice and islet transplantation were both performed as previous description [55]. Adult male and female C57BL/6J mice were selected as islet donor, and age-matched female C57BL/6J mice were chosen as recipient. C57BL/6J mice were injected with STZ at the dose of 170 mg/kg after 6 hours of fast. Mice that exhibited non-fasting hyperglycemia (> 350 mg/kg) with 3 consecutive detection were regarded as diabetic mice for islet transplantation. Each diabetic mouse was transplanted with about 350 islets, and mice that non-fasting blood glucose (< 144 mg/kg) recover normal were selected for physiological experiments 1 month post-transplant.

### Preparation of single islet cells

Mouse islets were isolated from either 6-8 week old male or female C57BL/6J and ICR mice. For islets isolation, 0.5 mg/mL collagenase P (11213873001, Roche, Basel, CH) was poured into pancreas by perfusion of the common bile duct. After digestion, islets were purified through Histopaque (11191 and 10771, Sigma Aldrich, St. Louis, MO, USA) gradient centrifugation. The Histopaque buffer was made with mixing Histopaque-10771 and Histopaque-11191 together as the ratio of 5:6. Purified islets were dissociated into single cells as follows: washed the islets with cold PBS at least two times, spun at 1000 rpm for 2 minutes, then replace PBS with 1mL TrypLE Express (12604021, Gibco, NY, USA) and incubated at 37^◦^C for 10-15 minutes with pipetting cell pellet by using P1000 pipette gently. Stop the reaction with low glucose DMEM (containing 10% FBS, 1% HEPES, 1% PenStrep), and centrifuged cells at 4^◦^C, 1000 rpm for 2 min. Then washed the cell pellet with cold PBS for 1 time, and the resuspended cells in PBS (containing 0.5%BSA) was filtered with a 40-micrometer strainer to get single cell suspension. To obtain live single cell, Calcein Blue, AM (C1429, Invitrogen, Carlsbad, CA, USA) was used to measure the viability of cell and high viability single cell was sorted by using BD FACS Aria II flow cytometry (Pleasanton CA, USA). Single islet cell was sorted into 96-well plates containing lysis buffer.

### Library construction and NGS sequencing

Single-cell RNA-seq libraries were constructed according to Smart-seq2 protocol except that oligod (T) primers comprising 16bp cell barcode sequence and 9bp molecular barcode sequence were used to allow sample pooling and molecular counting. In addition, part of Truseq read2 sequence was used to replace ISPCR sequence in the oligod (T) primer thus to be compatible with Illumina sequence platform. Cells in lysis buffer were denatured, reverse transcribed in the presence of template switching oligos and pre-amplified by adding both ISPCR and ISPCR-read2 primers with 24 PCR cycles. cDNA of cells with different barcodes were then pooled together and were purified using 0.8x AMPure XP beads. Homemade Tn5 enzyme was used to tagment cDNA. Final amplification was processed using P7-index primers (Truseq) and P5-index primers (Nextera). Sequencing libraries were purified twice using 0.6x AMPure XP beads and once with 1x AMPure XP beads, and were sequenced on Illumina HiSeq X10 platform with default parameters. Primer and adaptor sequences were all listed in supplementary Table S8 (information of barcode).

### Quantitative real-time PCR

The cDNA for quantitative real-time PCR synthesized following the protocol that we described in previous part (library construction). The β cell expressing *Ins2* from healthy and T2D mice of both sexes were identified by single cell qPCR. Quantitative real-time PCR was performed with SuperReal PreMix Plus (SYBR Green) (EP205, TIANGEN, Beijing, China) by using *Sdha* gene as an internal control. The relative expression levels of selected genes was analyzed based on the formula of 2^−ΔΔct^. The primers used for qPCR are listed in Table S7.

### Physiology experiment

For intraperitoneal injection glucose tolerance test, normal diet and high-fat diet feeding mice fasted for 16 hours. 1 g/Kg body weight of glucose was intraperitoneally injected, and blood glucose was measured at time point of 0 min, 15 min, 30 min, 60 min, 120 min after glucose injection using glucometer (ACCU-CHEK, Roche). Blood samples were collected from the mice that fasted for 6 hours, to measure the insulin level by using Mouse Ultrasensitive Insulin ELISA Kit (80-INSMSU-E01, ALPOC, Salem, NH, USA). For oral glucose tolerance test, mice were fasted for 6 hours, and glucose was gavaged at a dose of 2 g/Kg. Tail blood was collected for blood glucose detection, at time point of 0 min, 15 min, 30 min, 60 min, 120 min, using glucometer (ACCU-CHEK).

### Quantification and statistical analysis

#### Preprocessing before normalization

Reads were stored in paired-end fastq format. Reads of one end of fragment contain cell barcode and UMI information which was subsequently extracted and added to the name of corresponding reads of the other end in fastq file containing molecular sequence. That barcode/UMI information was in fastq-R2 files. Next, the generated single-end fastq files were cleaned by Trim Galore! http://www.bioinformatics.babraham.ac.uk/projects/trim_galore/ with the parameter --length 30.

Then, the FastQC https://www.bioinformatics.babraham.ac.uk/projects/fastqc/ was applied to check reads quality before subsequent alignment on mm9 genome using STAR [56]. The parameters of STAR software were --AlignEndsType EndToEnd --out FileterMismatchNoverReadLmax 0.04 --outSAMattrIHstart 0 --outSAMmultNmax 1 --outFilterMultimapNmax 1. GTF file of mm9 reference genome was derived from the RefSeq gene annotation [57] file downloaded from UCSC genome browser database, http://genome.ucsc.edu/cgi-bin/hgGateway?db=mm9. After alignment, SAM files underwent demultiplexing by Catadapt https://github.com/marcelm/cutadapt/releases with --overlap 16 –no-indels --match-read-wildcards parameters; reads were removed if they were assigned to more than one cells. Then, we removed PCR duplicates by UMI-tools [58]. Next, featureCounts [59] was utilized to quantify gene expression levels according to the above mentioned GTF file. Once the expression profiles were generated, cells with fewer than 500 expressed genes (UMI count > 1) were considered low-quality and were removed (Figure S1). The fraction of transcripts from mitochondrial genes in each single cell was investigated as it was used as an indicator for general quality control. Part of our ICR cells had high transcripts fraction of mitochondrial genes, which can be accounted for that kidney cells intrinsically express high-level mitochondrial genes [60]. So, we kept those cells for analysis because our transplantation cells were under renal capsule and either kidney cells or affected transplant cells may well be introduced in our data. Next, only genes expressed (UMI count > 1) in 5 or more cells were used for further analysis. Eventually, there are 14,152 genes and 4662 cells retained in mouse scRNA-seq data. The sample information of the Illumina high-throughput sequencing data is listed in Table 1.

#### Normalization

Here we used an adjusted count per million (CPM) normalization method, named adjCPM. The CPM method assumes that the total molecule reads are equal among cells. Similarly, our adjCPM method has an assumption except excluding a few genes in calculating the total number of molecular reads of each cell. It is based on the observation that sometimes a few top expressed genes can possess more than 50% total UMI count (Figure S1). These genes were selected as union set of the top 2 expressed genes of single cells in which the sum of the corresponding top 2 genes’ UMI counts is beyond 50% of the total count of the cell. Specifically, we obtained 9 genes (*Gcg*, *Ins1*, *Ins2*, *Malat1*, *Ppy*, *Pyy*, *Rn45s*, *Ttr*, *mt-Rnr2*) and they were excluded when calculating the total count of each cell. It is well known that the scRNA-seq experiments are affected by drop-out events especially for UMI method. Here, we also observed that many genes have zero expression value because of insufficient detection power or intrinsically under-expressed. To address this issue, SAVER [61] was applied to recover gene expression in the entire matrix.

### Cell type identification

Since cell type identification is critical for the rest analysis, two steps were carried out to robustly define the cell type. Firstly, we selected the top 10 expressed genes of each cell, resulting in a total of 118 genes to perform the subsequent hieratical clustering analysis. Accordingly, we primarily assigned 4 clusters of single cells after removing 111 cells which had ulterior distance in clustering dendrogram and annotated each cluster based on marker gene expression (Figure S1).

Secondly, by a lowess regression between the Log2 mean and Log2 coefficient of variance of each gene’s normalized UMI count across all single cells, we selected 3000 genes according to the residue of lowess regression as hypervariable genes (HVGs). Next, principal component analysis (PCA) was applied to the Log2 transformed normalized expression of these HVGs using *sklearn* [62] after centering and scaling, and top 25 PCs were selected based on the statistical significance of the fraction of total variance explained by each of them, which was estimated using jackstraw [63]. We used the top 25 PCs as input for subsequent t-Distributed Stochastic Neighbor Embedding (t-SNE) analysis. To have a stable t-SNE visualization, we tried a lot of combinations of parameters (perplexing [10, 15, 20, 25, 30], early exaggeration [12, 15, 20], learning rate [200, 300, 400, 500, 600, 800] and 50 random seeds) using *sklearn* [62], and found they generally gave very similar results. We finally chose perplexing:15, early exaggeration:12, learning rate:500 to generate the t-SNE plots shown in the figures. Next, Density-based spatial clustering algorithm, DBSCAN [64]with parameters eps=5 and MinPts=5 was applied on the resulting two-dimensional t-SNE map to group the single cells into cliques (Figure S1G). After removing singletons, we examined each single cell clique in order of clique size with following criteria: 1. only cells with consistent cluster/clique labels generated by hierarchical clustering using the top expressed genes and by DBSCAN were retained; 2. differential expression analysis of single cells in clique 3 (C3) compared the single cells in other cliques demonstrated that the single cells in C3 should be excluded because they highly expressed cell cycle related genes (Figure S1I). Finally, we kept 4467 high-quality mouse single cells for downstream analysis, and the cell type labels were inferred from the marker genes highly expressed in each single cell clique (Figure S1H).

#### Differential expression analysis and gene set enrichment analysis

MAST [17] was used to perform differential expression analysis between male and female or between T2D and healthy scRNA-seq profiles of β cells. Cellular detection rate (CDR, which means the number of genes detected in each single cell) was controlled as a covariate while estimating treatment effects. Significantly regulated genes were identified by using false discovery rate <= 0.05 as cutoff. For identification of functional pathways enriched in the detected differentially expressed genes between different sexes or between T2D and healthy conditions in our scRNA-seq data, we performed gene set enrichment analysis (GSEA) [19] on genes ranked based on their z-statistics, which were derived by mapping their MAST p-values to the standard normal distribution and using the sign of the Log2(fold change) of their expression values to represent the direction of regulation. Technically, the z-statistic of each gene was calculated as:

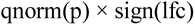

where p and lfc are p-value and Log2(fold change) of this gene obtained from MAST output, qnorm is the standard normal quantile function and sign is the signum function. At the same time, we also tried other methods to perform differential expression analysis, such as edgeR. performing differential analysis using edgeR combined with zingeR to incorporate an estimate of the dropout rate per cell (edgeR-zingeR), which is a DE-tool for UMI-based scRNA-seq data in the paper [65]. Using the same cutoff of MAST, abundant DEGs were also identified by edgeR-zingeR. In general, most significant DE genes from two separate methods are similar, most of which detected by MAST could also be covered by edgeR-zingeR (Figure S4), suggesting that the results of MAST are stable and could be repeated by another differential tool. The DEGs detected by edgeR are listed in Table S9 for reference. KEGG pathways [66] were used as input gene sets for GSEA, and fdr cutoff of 0.25 was used to select statistically significant pathways.

### Ethical statements

All experiments were performed in accordance with the University of Health Guide for the Care and Use of Laboratory Animals and were approved by the Biological Research Ethics Committee of Tongji University.

### Data availability

Raw data (Fastq files) for single pancreatic β cell RNA-seq in this study have been submitted to GSA at the National Genomics Data Center with accession number CRA003921 and can be accessed at https://bigd.big.ac.cn/gsa/s/FP8dq927.

## CRediT author statement

**Gang Liu:** Conceptualization, Methodology, Validation, Investigation, Visualization, Writing - original draft, Writing - review & editing. **Yana Li:** Software, Formal analysis, Data curation, Visualization, Data curation, Writing - original draft, Writing review & editing. **Tengjiao Zhang:** Methodology, Resources, Investigation, Writing original draft. **Mushan Li:** Software, Formal analysis, Data curation, Writing - original draft. **Sheng Li:** Validation, Resources. **Qing He:** Validation, Resources. **Shuxin Liu:** Resources. **Minglu Xu:** Resources. **Tinghui Xiao:** Resources. **Zhen Shao:** Visualization, Supervision, Writing - original draft, Writing - review & editing, Project administration, Funding acquisition. **Weiyang Shi:** Visualization, Supervision, Writing - original draft, Project administration, Funding acquisition. **Weida Li:** Conceptualization, Visualization, Supervision, Project administration, Funding acquisition. All authors read and approved the final manuscript.

## Acknowledgments

This work was supported by the National Key Research and Development Program of China (2016YFA0102200, 2017YFA0106500, 2018YFA0107102, 2020YFA0112500) awarded to WL; and the National Key Research and Development Program of China (2018YFA0107602) awarded to ZS; WS is supported by the Funding Project of National Key Research and Development Program of China (2018YFD0900604), Natural Science Foundation of China (41676119, 41476120) and start-up fund from Ocean University of China.

---

**Figure S1.**
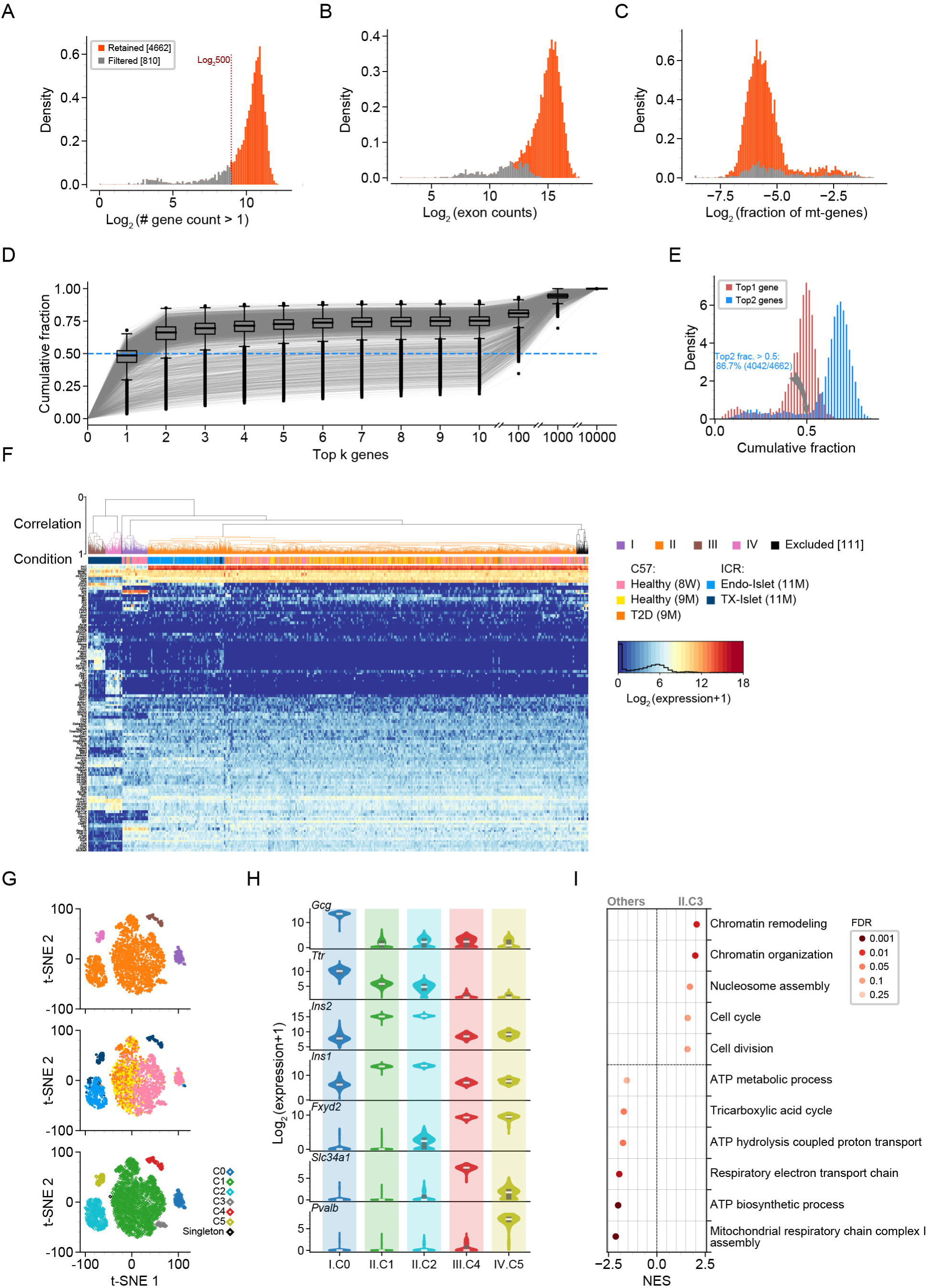
Data processing identifies a set of qualified single-cell RNA sequencing profiles of mouse islet. **A.** Distribution of detected gene numbers (Log2 transformed). 810 out of 5472 cells are removed from further analysis due to number of genes with more than 1 UMI count less than 500. **B.** Distribution of total exon counts (Log2 transformed). The retained cells (shown in orange) have relatively higher exon counts. **C.** Distribution of UMI fractions of mitochondrial genes (Log2 transformed). The retained cells (shown in orange) have relatively lower UMI fractions of mitochondrial genes. **D.** Boxplot of fraction of UMI count of top k most expressed genes. The fraction is beyond 0.5 for most cells when k = 2 Each gray line represents a cell. **E.** Distribution of UMI count fraction of top 1 (shown in red) and top 2 (shown in blue) most expressed genes. In 86.7% of the mouse islet cells, the UMI count fraction of the top 2 most expressed genes are more than 0.5. The union of the top 2 most expressed genes from all these cells includes 9 genes (*Gcg*, *Ins1*, *Ins2*, *Malat1*, *Ppy*, *Pyy*, *Rn45s*, *Ttr*, *mt-Rnr2*), which are excluded while calculating total count for normalization using adjCPM method. **F.** Hierarchical clustering for 4,662 mouse islet single cells. Using union of the top 10 expressed genes in all cells (total 141 genes) with “euclidean” metric and “complete” linkage and 110 cells are excluded. 8W, 8-week-old; 9M, 9-month-old; 11M, 11-month-old; Endo-islet, endogenous islets of recipients; TX-islet, transplanted islets in recipients. **G.** t-SNE map of retained cells. Cells are colored by the clustering result of the previous hierarchical clustering (up panel), conditions (middle panel) and clique labels from DBSCAN clustering (down panel), respectively. **H.** Violin plots of marker genes for C0, C1, C2, C4, C5 after filtering cells with inconsistent cluster labels given by previous hierarchical clustering. C1: *Ins2* (a known marker gene of β cells) highly expressing cells from mice of C57 strain (labeled as β cells); C2: *Ins2* highly expressing cells from 11-month-old mice of ICR strain (labeled as β cells); C0: *Gcg* (a known marker gene of α cells) highly expressing cells (labeled as α cells); C4 and C5: two groups of cells with high expression of *Slc34a1* and *Pvalb* (two known marker genes of kidney proximal tubule and distal convoluted tubule cells), respectively, which were labeled as kidney-like cells and were discarded. **I.** GO enrichment analysis for up- and down-regulated genes in C3. Key metabolic processes such as ATP biosynthetic process associated genes are down-regulated and cell cycle related genes are up-regulated in C3, so this clique is removed as cells in cell cycle.

**Figure S2.**
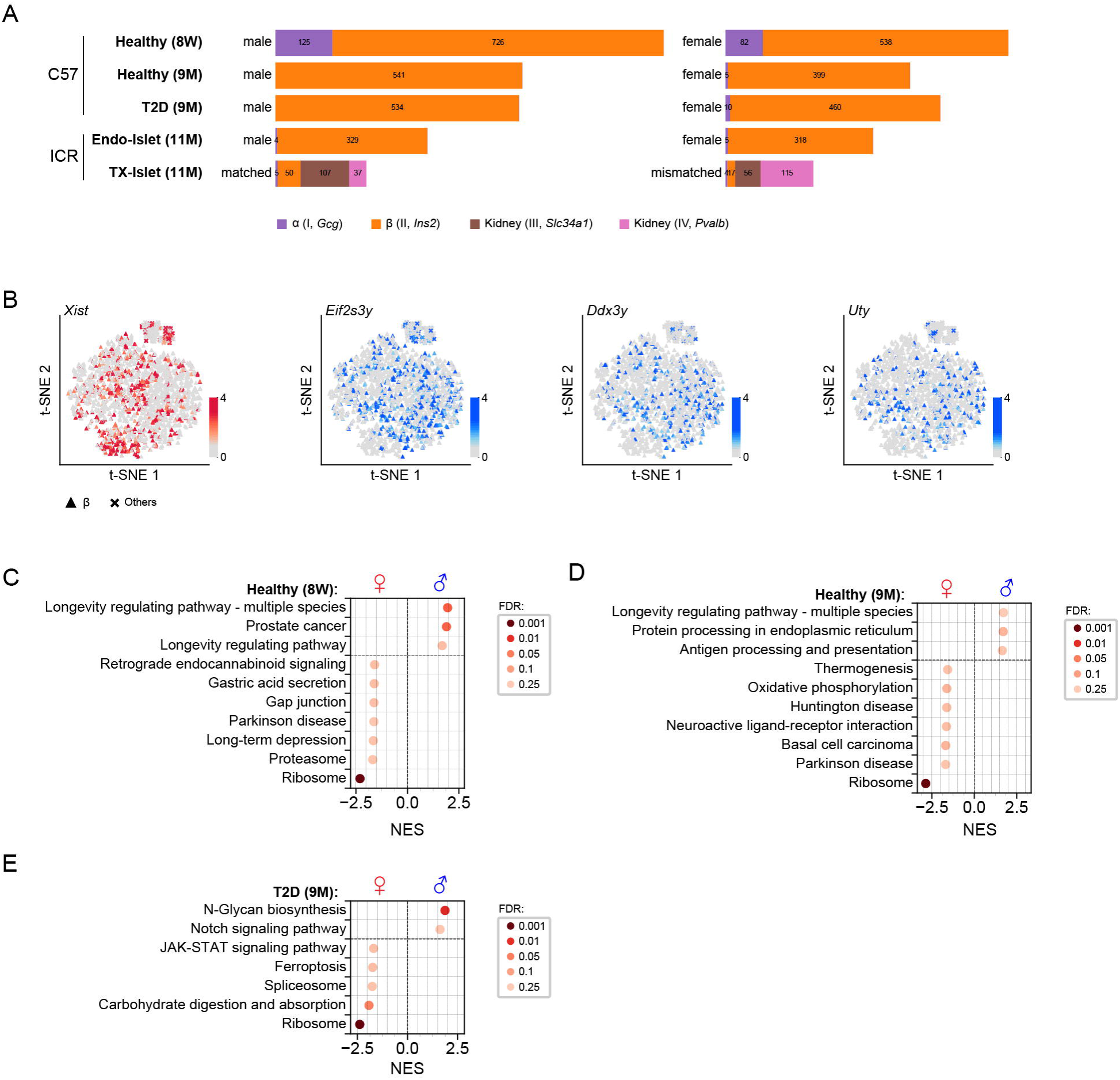
Single-cell RNA sequencing reveals sexually dimorphic gene expression and gene set enrichment in β cells. **A.** Barplot showing the cell type composition of male and female mice under different conditions. 8W, 8-week-old; 9M, 9-month-old; 11M, 11-month-old; Endo-islet, endogenous islets of recipients; TX-islet, transplanted islets in recipients. **B.** t-SNE map of retained mouse cells colored by expression of sex chromosome linked genes. The reds for X-chromosome linked gene *Xist*, and the blues for Y-chromosome linked genes *Eif2s3y*, *Ddx3y*, *Uty*. **C.** GSEA for sex comparisons of β cells from 8-week-old C57 mice. The gene sets with NES > 0 are enriched in males and NES < 0 in females. **D.** GSEA for sex comparisons of β cells from healthy 9-month-old C57 mice. The gene sets with NES > 0 are enriched in males and NES < 0 in females. **E.** GSEA for sex comparisons of β cells from T2D 9-month-old C57 mice. The gene sets with NES > 0 are enriched in males and NES < 0 in females.

**Figure S3.**
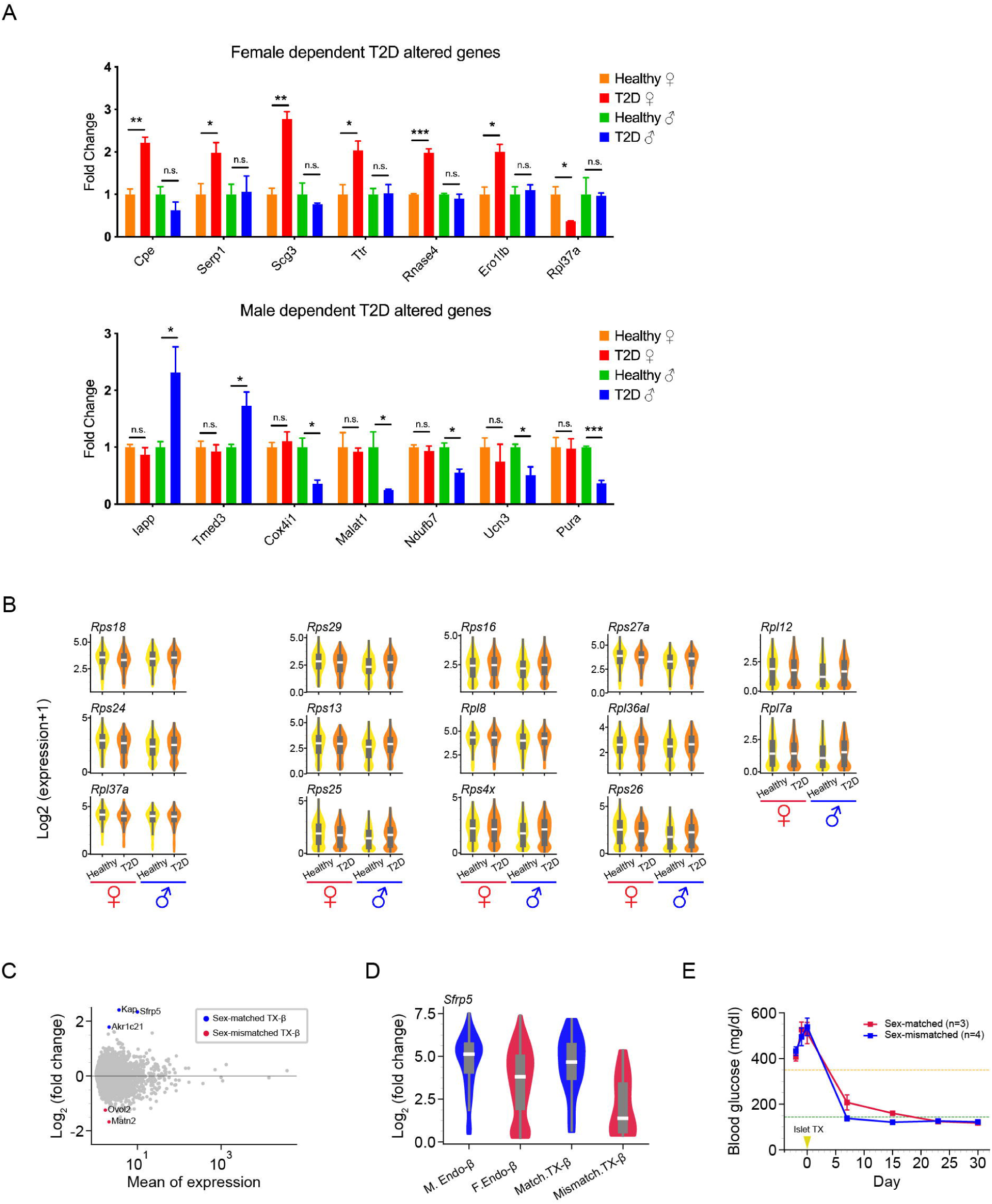
Single-cell RNA sequencing reveals sexually dimorphic genes in β cells from T2D model and transplanted islet. **A.** The qPCR analysis of sex-dependent T2D altered genes. Sex-dependent T2D altered genes with basemean > 10 are selected for validation. n=3 biological replicates. For each replicate, cDNA of 5 *Ins2*^+^ cells that randomly selected from scRNA-seq cDNA library was mixed with equal volume. The cDNA for qPCR is synthesized following the protocol of sequencing library construction. The relative expression (fold change) is calculated relative to the gene expression level of healthy condition within the same sex. Results are presented as mean±SEM; * p ≤ 0.05, ** p ≤ 0.01, *** p ≤ 0.001, n.s., not significant, two sample t-test. **B.** Violin plots of sex-dependent T2D altered genes involved in ribosome pathway. The left 3 down-regulated genes in female T2D β cells are female-dependent T2D altered genes; the right 11 up-regulated genes in T2D β cells are male-dependent T2D altered genes. **C.** MA plot showing the comparison between sex matched and mismatched transplant-β cells. Up-regulated and down-regulated genes in sex matched transplant-β cells are highlighted in blue and red, respectively. See also in Table S5. TX-β, transplant-β cells. **D.** Violin plot of expression of *Sfrp5* in endogenous and transplanted β cells. M.Endo-β, endogenous β cells from male recipient; F.Endo-β, endogenous β cells from female recipient; Match.TX-β, sex matched transplant-β cells; Mismatch.TX-β, sex mismatched transplant-β cells. **E.** Blood glucose level before and after islet transplantation in diabetic mice. The blood glucose level of diabetic mice with sex-matched (blue) and sex-mismatched (red) transplanted islet are all recovered normal (green baseline) from hyperglycemia (orange baseline) about 3 weeks after transplantation. TX, transplantation.

**Figure S4.**
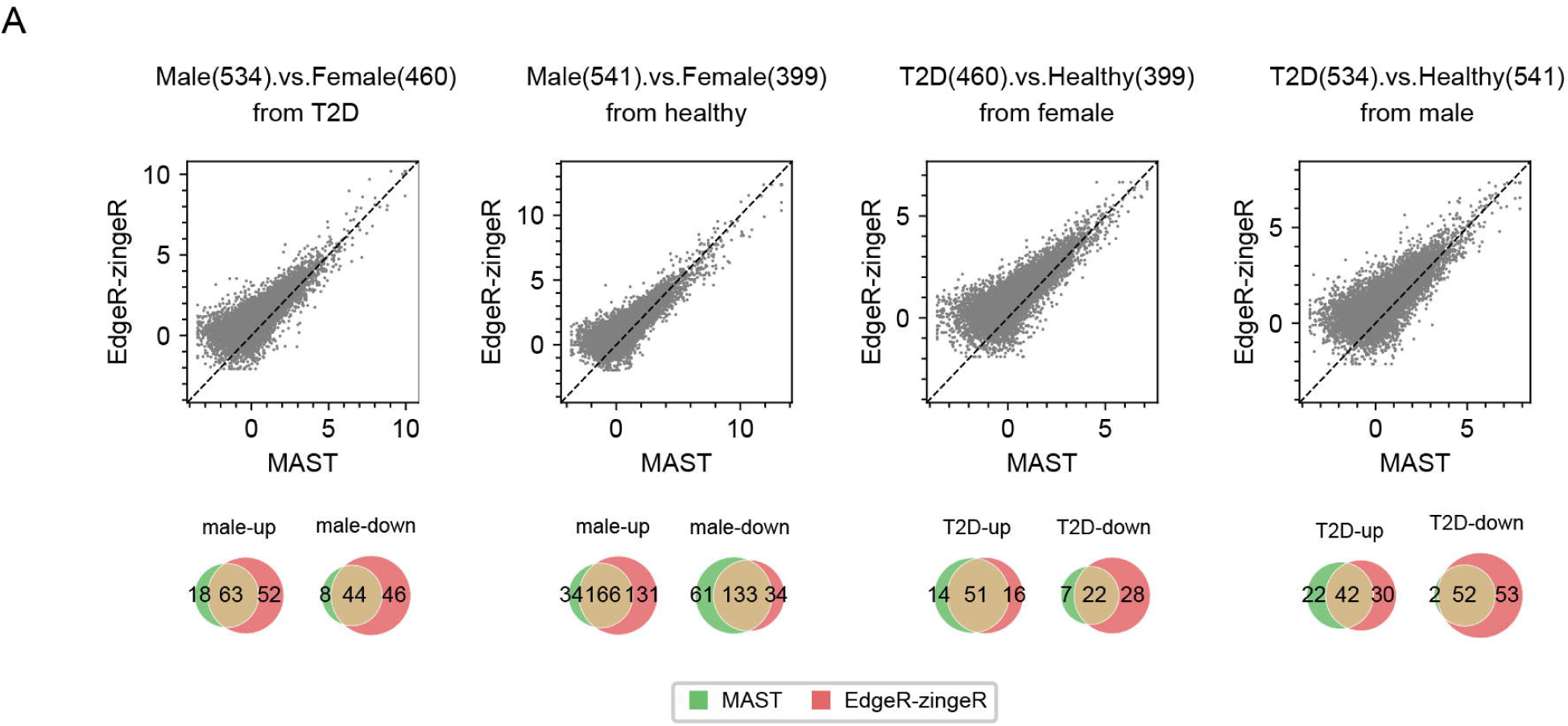
Differential expression genes called by MAST and edgeR have overlap. Scatter plots showing that p-value derived z-statistics of genes obtained by MAST and edgeR-zingeR are similar in different analysis. And venn diagrams showing the overlap of DEGs detected by MAST and edgeR-zingeR. The number in brackets represent the number of cells that were used for analysis in each condition.

**Table S1 Sex-biased expression genes in β cell of 8-week-old healthy mice**

**Table S2 Sex-biased expression genes in β cell of 9-mounth-old healthy mice**

**Table S3 Sex-biased expression genes in β cell of 9-mounth-old T2D mice**

**Table S4 Sex biased genes included in the leading edge gene of GSEA**

**Table S5 Sex-independent differential expression genes**

**Table S6 Sex-dependent T2D altered genes**

**Table S7 Primers for qPCR**

**Table S8 Primers for single-cell RNA sequencing library construction**

**Table S9 Differential expression genes detected by edgeR**

