## Supplemental table 1 to 9 for "Mouse Single Islet β Cell Transcriptomics Reveal Sexually Dimorphic Transcriptomes and Type 2 Diabetes Genes": Table 1.docx

| **Strain** | **Age** | **Condition** | **Theoretic number of cells** | **Total** |
| --- | --- | --- | --- | --- |
| C57BL/6J | 6-8 weeks | Healthy | Male:1056 Female:768 | 1824 |
| C57BL/6J | 9 months | Diabetic | Male:576 Female:576 | 1152 |
| C57BL/6J | 9 months | Healthy | Male:576 Female:480 | 1056 |
| ICR | 11 months | Endogenous | Male:384 Female:384 | 768 |
| ICR | 11 months | Transplanted | Male:288 Female:384 | 672 |
| Total 5472 | | | | |

**Table 1 The number of collected cells from mice in different conditions**
