## Supplemental table 1 to 9 for "Mouse Single Islet β Cell Transcriptomics Reveal Sexually Dimorphic Transcriptomes and Type 2 Diabetes Genes": Table S1.docx

**Table S1 Sex-biased expression genes in β cell of healthy 8-week-old mice**

| **Type** | **Gene symbol** | **Basemean** | **log2FoldChange** | **P value** | **P adjust** |
| --- | --- | --- | --- | --- | --- |
| Upregulated genes in male | *Eif2s3y* | 1.486932 | 0.933449 | 3.18E-72 | 1.4E-68 |
|  | *Ddx3y* | 1.2563 | 0.513994 | 1.36E-43 | 3.96E-40 |
|  | *Uty* | 1.19381 | 0.418525 | 5E-36 | 1.1E-32 |
|  | *Necab2* | 2.804386 | 0.803933 | 2.14E-23 | 2.69E-20 |
|  | *Ins1* | 11837.26 | 0.265875 | 6.68E-15 | 4.51E-12 |
|  | *Dapl1* | 2.984041 | 0.695924 | 1.84E-13 | 1.01E-10 |
|  | *Hspa8* | 22.87728 | 0.303789 | 1.26E-12 | 5.88E-10 |
|  | *Ero1lb* | 43.07791 | 0.35595 | 7.27E-12 | 3.19E-09 |
|  | *Gc* | 1.798429 | 0.42366 | 1.26E-10 | 4.42E-08 |
|  | *Sfrp5* | 2.295776 | 0.576154 | 2.59E-09 | 8.13E-07 |
|  | *Atp2a3* | 5.298574 | 0.357234 | 3.46E-08 | 9.78E-06 |
|  | *Malat1* | 878.4696 | 0.206466 | 4.6E-08 | 1.22E-05 |
|  | *Arl4a* | 1.960551 | 0.360029 | 3.23E-07 | 1.4E-68 |
|  | *Sez6l2* | 4.965141 | 0.357369 | 2.09E-06 | 3.96E-40 |
|  | *Ttr* | 53.7695 | 0.278966 | 3.7E-06 | 1.1E-32 |
|  | *Rn45s* | 795.7 | 0.202101 | 4.66E-06 | 2.69E-20 |
|  | *Pam* | 7.185896 | 0.328695 | 5.33E-06 | 4.51E-12 |
|  | *Gpr137b-ps* | 2.292307 | 0.292191 | 8.74E-06 | 1.01E-10 |
|  | *Pdyn* | 1.609304 | 0.387507 | 9.32E-06 | 5.88E-10 |
|  | *Cox6a2* | 3.588717 | 0.397078 | 1.27E-05 | 3.19E-09 |
|  | *Ntrk2* | 2.743986 | 0.287812 | 2.53E-05 | 4.42E-08 |
|  | *Depp1* | 1.458827 | 0.253881 | 4.03E-05 | 8.13E-07 |
|  | *Atp2a2* | 11.04407 | 0.287706 | 5.48E-05 | 9.78E-06 |
|  | *Dnajb9* | 5.345844 | 0.2684 | 0.000114 | 1.22E-05 |
|  | *Tmod2* | 3.835045 | 0.272921 | 0.000168 | 1.4E-68 |
|  | *Fkbp11* | 3.208839 | 0.327917 | 0.000184 | 3.96E-40 |
|  | *Cct8* | 3.952204 | 0.259018 | 0.000226 | 1.1E-32 |
|  | *Terf2* | 1.471991 | 0.215872 | 0.000234 | 2.69E-20 |
|  | *Tmed8* | 1.606234 | 0.214518 | 0.000292 | 4.51E-12 |
|  | *Armcx2* | 1.705799 | 0.217794 | 0.000391 | 1.01E-10 |
|  | *Os9* | 9.191558 | 0.251217 | 0.000567 | 5.88E-10 |
|  | *Qsox1* | 2.13188 | 0.224304 | 0.000592 | 3.19E-09 |
|  | *Tmem176a* | 2.512854 | 0.270973 | 0.000617 | 4.42E-08 |
|  | *Abcc8* | 10.62819 | 0.21968 | 0.00063 | 8.13E-07 |
|  | *4732471J01Rik* | 1.598672 | 0.217944 | 0.000853 | 9.78E-06 |
|  | *Hikeshi* | 1.620698 | 0.203613 | 0.000934 | 1.22E-05 |
|  | *Hsp90aa1* | 9.724375 | 0.251789 | 0.000958 | 1.4E-68 |
| Upregulated genes in female | *Xist* | *1.736828* | *-1.33529* | *7E-136* | 6.1E-132 |
|  | *Gnai2* | *28.30528* | *-0.53049* | *8.91E-26* | 1.56E-22 |
|  | *Enpp2* | *3.607559* | *-0.69724* | *1.49E-24* | 2.18E-21 |

**Table S1 Sex-biased expression genes in β cell of healthy 8-week-old mice (continued)**

| **Type** | **Gene symbol** | **Basemean** | **log2FoldChange** | **P value** | **P adjust** |
| --- | --- | --- | --- | --- | --- |
| Upregulated genes in female | *Sh3pxd2a* | 8.713551 | -0.6853 | 3.75E-19 | 4.11E-16 |
|  | *Hnrnpdl* | 4.666779 | -0.62076 | 5.83E-19 | 5.68E-16 |
|  | *Arpc1a* | 11.67276 | -0.62702 | 2.27E-17 | 2E-14 |
|  | *Cish* | 1.4946 | -0.42814 | 1.56E-16 | 1.24E-13 |
|  | *Zbtb20* | 4.178588 | -0.6063 | 1.49E-15 | 1.09E-12 |
|  | *mt-Nd1* | 83.47833 | -0.28322 | 2.75E-14 | 1.72E-11 |
|  | *Gck* | 3.369572 | -0.4442 | 5.42E-14 | 3.17E-11 |
|  | *Ddr1* | 3.745304 | -0.44963 | 1.5E-11 | 6.25E-09 |
|  | *Rpl5* | 59.77551 | -0.24079 | 2.66E-11 | 1.01E-08 |
|  | *Atp5a1* | 25.24974 | -0.33873 | 4.95E-11 | 1.81E-08 |
|  | *Glrx5* | 8.559683 | -0.43834 | 1.12E-09 | 3.77E-07 |
|  | *Appl2* | 5.730756 | -0.48993 | 1.3E-09 | 4.23E-07 |
|  | *Relt* | 2.623232 | -0.40005 | 4.23E-09 | 1.28E-06 |
|  | *Rps12* | 4.97714 | -0.48524 | 4.42E-09 | 1.29E-06 |
|  | *Mta1* | 1.8059 | -0.28735 | 4.61E-08 | 1.22E-05 |
|  | *Atp11b* | 1.555947 | -0.27904 | 8.72E-08 | 2.25E-05 |
|  | *Sfi1* | 1.814225 | -0.28849 | 9.5E-08 | 2.38E-05 |
|  | *Neurod1* | 5.002035 | -0.35509 | 1.02E-07 | 2.47E-05 |
|  | *Hnrnpa2b1* | 8.54366 | -0.33451 | 1.04E-07 | 2.47E-05 |
|  | *Atp6v1a* | 6.236968 | -0.39343 | 1.1E-07 | 2.54E-05 |
|  | *Wsb2* | 4.446329 | -0.36658 | 1.41E-07 | 3.1E-05 |
|  | *Gmpr* | 20.38672 | -0.29608 | 1.42E-07 | 3.1E-05 |
|  | *Fam174b* | 14.51884 | -0.30578 | 1.45E-07 | 3.1E-05 |
|  | *Ptma* | 15.74249 | -0.24473 | 2.32E-07 | 4.85E-05 |
|  | *Ubqln1* | 2.33663 | -0.33048 | 2.98E-07 | 6.07E-05 |
|  | *Mrgbp* | 1.848512 | -0.30803 | 3.24E-07 | 6.31E-05 |
|  | *Bex4* | 3.791685 | -0.33834 | 6.28E-07 | 0.00012 |
|  | *Cirbp* | 4.903129 | -0.36675 | 7.08E-07 | 0.000132 |
|  | *Rps3a1* | 9.979755 | -0.31142 | 7.37E-07 | 0.000135 |
|  | *Rfx6* | 3.765788 | -0.36914 | 8.19E-07 | 0.000147 |
|  | *Rammet* | 2.504908 | -0.28516 | 1.5E-06 | 0.000257 |
|  | *Casp3* | 1.348331 | -0.2032 | 2E-06 | 0.000337 |
|  | *Ncoa1* | 8.233512 | -0.40285 | 2.32E-06 | 0.000376 |
|  | *Tcf4* | 1.850338 | -0.25448 | 2.6E-06 | 0.000413 |
|  | *Abcb10* | 6.786951 | -0.28043 | 3.15E-06 | 0.000481 |
|  | *Cttn* | 5.286523 | -0.35176 | 3.18E-06 | 0.000481 |
|  | *Ppia* | 4.723158 | -0.23513 | 3.39E-06 | 0.000504 |
|  | *Scgn* | 9.887207 | -0.33334 | 3.78E-06 | 0.000539 |
|  | *Sf3b2* | 11.03542 | -0.30513 | 3.81E-06 | 0.000539 |
|  | *Eef1a1* | 30.96339 | -0.20437 | 4.15E-06 | 0.000577 |

**Table S1 Sex-biased expression genes in β cell of healthy 8-week-old mice (continued)**

| **Type** | **Gene symbol** | **Basemean** | **log2FoldChange** | **P value** | **P adjust** |
| --- | --- | --- | --- | --- | --- |
| Upregulated genes in female | *Peg13* | 2.829678 | -0.31745 | 5.93E-06 | 0.000776 |
|  | *Ndufb3* | 3.525947 | -0.25151 | 7.66E-06 | 0.000986 |
|  | *Paip2* | 12.31065 | -0.24359 | 7.82E-06 | 0.000986 |
|  | *Bmpr1a* | 2.065966 | -0.25226 | 7.87E-06 | 0.000986 |
|  | *Ppm1g* | 2.786198 | -0.28038 | 9.84E-06 | 0.001177 |
|  | *Rps24* | 7.024712 | -0.25439 | 9.93E-06 | 0.001177 |
|  | *Sdhd* | 2.793973 | -0.31346 | 1.22E-05 | 0.001407 |
|  | *Bcl7b* | 2.868987 | -0.31146 | 1.39E-05 | 0.001539 |
|  | *Iffo1* | 2.637723 | -0.30216 | 1.5E-05 | 0.00165 |
|  | *Atrn* | 2.559996 | -0.28197 | 1.72E-05 | 0.001862 |
|  | *Lad1* | 1.46443 | -0.24662 | 1.75E-05 | 0.001873 |
|  | *Pabpc1* | 13.70041 | -0.3129 | 1.78E-05 | 0.001885 |
|  | *Ufl1* | 7.594142 | -0.31276 | 1.84E-05 | 0.001921 |
|  | *Rpl23a* | 8.073745 | -0.25631 | 1.88E-05 | 0.001942 |
|  | *Eif4a3* | 6.301812 | -0.30801 | 2.03E-05 | 0.002075 |
|  | *Rpl22l1* | 4.777776 | -0.30667 | 2.16E-05 | 0.002179 |
|  | *Stard4* | 2.32882 | -0.288 | 3.03E-05 | 0.002951 |
|  | *Csk* | 1.532568 | -0.22749 | 3.03E-05 | 0.002951 |
|  | *Kcmf1* | 4.249813 | -0.30652 | 3.22E-05 | 0.003108 |
|  | *Acadm* | 1.942246 | -0.26596 | 3.71E-05 | 0.003502 |
|  | *Mysm1* | 3.692042 | -0.30101 | 3.86E-05 | 0.0036 |
|  | *Aes* | 7.876177 | -0.2929 | 3.98E-05 | 0.003674 |
|  | *Mid1ip1* | 2.128018 | -0.26663 | 4.09E-05 | 0.003699 |
|  | *Zfp106* | 8.085221 | -0.28906 | 4.29E-05 | 0.003837 |
|  | *Nipal3* | 10.57332 | -0.24032 | 4.56E-05 | 0.00404 |
|  | *Jak1* | 3.504193 | -0.30065 | 5.07E-05 | 0.004442 |
|  | *Snx4* | 2.420301 | -0.26574 | 5.95E-05 | 0.005067 |
|  | *Meis2* | 2.87168 | -0.2724 | 6.5E-05 | 0.00548 |
|  | *Rpl7a* | 3.425001 | -0.25139 | 7.39E-05 | 0.006067 |
|  | *Desi1* | 9.133219 | -0.26945 | 7.4E-05 | 0.006067 |
|  | *Lmnb2* | 2.154541 | -0.26956 | 7.53E-05 | 0.006117 |
|  | *Rpl9* | 5.75942 | -0.26429 | 8.87E-05 | 0.00714 |
|  | *Rps5* | 10.63005 | -0.26004 | 9.12E-05 | 0.007269 |
|  | *Sidt2* | 1.669668 | -0.23465 | 0.000101 | 0.00795 |
|  | *Ahcyl1* | 2.770119 | -0.27018 | 0.000102 | 0.008009 |
|  | *Ppp1r13b* | 4.212516 | -0.28295 | 0.000113 | 0.008754 |
|  | *Rps28* | 4.071481 | -0.20629 | 0.000138 | 0.010403 |
|  | *Rpl23* | 21.96928 | -0.25163 | 0.000145 | 0.010885 |
|  | *Hist1h4h* | 1.587344 | -0.24418 | 0.000147 | 0.010929 |
|  | *Tjp2* | 1.695079 | -0.2139 | 0.000175 | 0.012441 |
|  | *Tmed5* | 1.711216 | -0.20447 | 0.000176 | 0.012441 |

**Table S1 Sex-biased expression genes in β cell of healthy 8-week-old mice (continued)**

| **Type** | **Gene symbol** | **Basemean** | **log2FoldChange** | **P value** | **P adjust** |
| --- | --- | --- | --- | --- | --- |
| Upregulated genes in female | *Llgl2* | 2.148208 | -0.2348 | 0.000183 | 0.012809 |
|  | *Rpl39* | 4.091345 | -0.25805 | 0.000189 | 0.01307 |
|  | *Sap18* | 2.064641 | -0.23722 | 0.000197 | 0.013508 |
|  | *Pax6* | 5.140614 | -0.24955 | 0.000203 | 0.013829 |
|  | *Ap1s2* | 4.256881 | -0.27075 | 0.000212 | 0.014205 |
|  | *Rpl32* | 11.86322 | -0.2159 | 0.000223 | 0.014783 |
|  | *Rps21* | 9.880253 | -0.27471 | 0.000238 | 0.015454 |
|  | *Nsun2* | 2.535707 | -0.24663 | 0.000252 | 0.015911 |
|  | *Kcnb1* | 3.564496 | -0.2948 | 0.000263 | 0.016481 |
|  | *Sult1d1* | 1.635052 | -0.25117 | 0.000274 | 0.017028 |
|  | *Fus* | 2.585693 | -0.22577 | 0.00029 | 0.017704 |
|  | *Gadd45g* | 4.242324 | -0.30826 | 0.000293 | 0.017704 |
|  | *Rplp1* | 12.41948 | -0.23648 | 0.000303 | 0.018192 |
|  | *Pde4b* | 2.037175 | -0.24745 | 0.000349 | 0.020812 |
|  | *Wdr7* | 2.217493 | -0.23754 | 0.000354 | 0.02088 |
|  | *Spop* | 3.068441 | -0.24135 | 0.000355 | 0.02088 |
|  | *Gramd1a* | 3.006773 | -0.25217 | 0.000398 | 0.022989 |
|  | *Spint2* | 9.363057 | -0.21522 | 0.000475 | 0.026405 |
|  | *Rpl27a* | 3.400188 | -0.20049 | 0.000486 | 0.026813 |
|  | *Gnaz* | 8.543707 | -0.27837 | 0.000503 | 0.027557 |
|  | *Rps10* | 2.882652 | -0.22126 | 0.000508 | 0.027696 |
|  | *Ppm1l* | 2.284931 | -0.23065 | 0.000538 | 0.029117 |
|  | *Akr1c19* | 2.278034 | -0.27804 | 0.000553 | 0.02969 |
|  | *Prlr* | 15.47032 | -0.25195 | 0.000573 | 0.030176 |
|  | *Gcg* | 4.506283 | -0.36332 | 0.000575 | 0.030176 |
|  | *Arpc1b* | 2.11007 | -0.24822 | 0.00058 | 0.030285 |
|  | *Cbx6* | 3.198832 | -0.26546 | 0.000592 | 0.03053 |
|  | *Aff4* | 4.563376 | -0.25729 | 0.000632 | 0.031874 |
|  | *Parva* | 3.827355 | -0.24687 | 0.000646 | 0.032355 |
|  | *Rpl21* | 14.69107 | -0.22594 | 0.000741 | 0.036901 |
|  | *Lias* | 3.449385 | -0.27395 | 0.000776 | 0.038471 |
|  | *Rps13* | 7.414783 | -0.23306 | 0.000798 | 0.039299 |
|  | *Mars* | 3.183157 | -0.24807 | 0.000854 | 0.041482 |
|  | *Nectin3* | 2.326381 | -0.23638 | 0.000859 | 0.041482 |
|  | *Ank* | 2.012045 | -0.21347 | 0.000861 | 0.041482 |
|  | *Exoc3* | 4.777531 | -0.27706 | 0.000902 | 0.04254 |
|  | *Rps8* | 10.26809 | -0.22797 | 0.000931 | 0.043558 |
|  | *Snrpg* | 2.010335 | -0.20097 | 0.000961 | 0.043833 |
|  | *Mdm1* | 2.984605 | -0.23854 | 0.000984 | 0.044478 |
|  | *Sfxn1* | 4.105434 | -0.23403 | 0.000992 | 0.044624 |
|  | *Rab3b* | 2.19273 | -0.23106 | 0.00104 | 0.046531 |
