## Supplemental table 1 to 9 for "Mouse Single Islet β Cell Transcriptomics Reveal Sexually Dimorphic Transcriptomes and Type 2 Diabetes Genes": Table S2.docx

**Table S2 Sex-biased expression genes in β cell of healthy 9-mounth-old mice**

| **Type** | **Gene symbol** | **Basemean** | **log2FoldChange** | **P value** | **P adjust** |
| --- | --- | --- | --- | --- | --- |
| Upregulated genes in male | *Necab2* | 3.244283 | 1.425571 | 5.68E-56 | 2.55E-52 |
|  | *Eif2s3y* | 1.47123 | 0.913968 | 2.32E-51 | 6.96E-48 |
|  | *Ins2* | 33498.4 | 0.334646 | 6.34E-49 | 1.42E-45 |
|  | *Scg2* | 161.5059 | 0.473698 | 4.03E-48 | 7.24E-45 |
|  | *Ddx3y* | 1.306758 | 0.645863 | 1.2E-41 | 1.79E-38 |
|  | *Malat1* | 1307.045 | 0.428951 | 3.47E-37 | 3.46E-34 |
|  | *Gc* | 1.910597 | 0.898893 | 4.79E-31 | 3.59E-28 |
|  | *Uty* | 1.204316 | 0.448413 | 6.86E-28 | 4.4E-25 |
|  | *Rnase4* | 21.35818 | 0.728147 | 4.59E-27 | 2.75E-24 |
|  | *Syt13* | 24.30754 | 0.573866 | 8.99E-23 | 4.76E-20 |
|  | *Naa20* | 1.743021 | 0.565373 | 1.97E-22 | 9.82E-20 |
|  | *Tssc4* | 7.340657 | 0.67588 | 2.2E-20 | 9.87E-18 |
|  | *Sfrp5* | 3.002722 | 1.072565 | 3.84E-20 | 1.64E-17 |
|  | *Cpe* | 44.66806 | 0.468883 | 5.48E-20 | 2.24E-17 |
|  | *Ubc* | 21.09991 | 0.473789 | 5.35E-17 | 1.85E-14 |
|  | *Paip2* | 8.012545 | 0.555365 | 6.26E-17 | 2.08E-14 |
|  | *Ndufb7* | 13.9671 | 0.523597 | 2.03E-16 | 6.52E-14 |
|  | *Tspan33* | 5.763164 | 0.670725 | 1.15E-14 | 3.14E-12 |
|  | *Il1r1* | 10.52837 | 0.606491 | 1.59E-14 | 4.22E-12 |
|  | *Fxyd6* | 3.544412 | 0.64288 | 2.5E-14 | 6.43E-12 |
|  | *Ccnd1* | 2.130443 | 0.523817 | 7.94E-14 | 1.88E-11 |
|  | *Cd47* | 3.353938 | 0.563217 | 1.16E-13 | 2.68E-11 |
|  | *Nkx2-2* | 2.384583 | 0.50452 | 2.98E-13 | 6.7E-11 |
|  | *Manf* | 22.46666 | 0.465525 | 6.78E-13 | 1.42E-10 |
|  | *Acly* | 11.45246 | 0.390063 | 9.5E-13 | 1.9E-10 |
|  | *Serp1* | 17.42789 | 0.422386 | 1.07E-12 | 2.09E-10 |
|  | *Rnf5* | 1.882625 | 0.473384 | 1.34E-12 | 2.56E-10 |
|  | *Spp1* | 1.338963 | 0.526562 | 1.47E-12 | 2.74E-10 |
|  | *Cox4i1* | 74.03853 | 0.332282 | 2.65E-12 | 4.86E-10 |
|  | *Nisch* | 24.87775 | 0.310775 | 3.91E-12 | 6.75E-10 |
|  | *Bsg* | 25.12124 | 0.314884 | 4.28E-12 | 7.12E-10 |
|  | *Tmem160* | 6.157395 | 0.567986 | 6.38E-12 | 1.01E-09 |
|  | *Ccnd2* | 59.42288 | 0.35551 | 2.02E-11 | 2.97E-09 |
|  | *Kmt2d* | 4.84398 | 0.50721 | 5.02E-11 | 7.17E-09 |
|  | *Tmod2* | 4.271286 | 0.517074 | 5.47E-11 | 7.68E-09 |
|  | *Scg3* | 36.79523 | 0.282196 | 6.03E-11 | 8.22E-09 |
|  | *Ucn3* | 57.46857 | 0.344322 | 6.58E-11 | 8.82E-09 |
|  | *Mrps28* | 1.207453 | 0.26108 | 6.95E-11 | 9.19E-09 |
|  | *Ttr* | 45.60207 | 0.410865 | 7.57E-11 | 9.86E-09 |
|  | *Hap1* | 3.856234 | 0.491932 | 4.49E-10 | 5.24E-08 |
|  | *Etv1* | 1.347278 | 0.359371 | 5.33E-10 | 6.14E-08 |

**Table S2 Sex-biased expression genes in β cell of healthy 9-mounth-old mice (continued)**

| **Type** | **Gene symbol** | **Basemean** | **log2FoldChange** | **P value** | **P adjust** |
| --- | --- | --- | --- | --- | --- |
| Upregulated genes in male | *Psap* | 8.560915 | 0.34477 | 6.9E-10 | 7.75E-08 |
|  | *Pcp4* | 1.577358 | 0.45092 | 9.28E-10 | 1.03E-07 |
|  | *Srsf7* | 3.826995 | 0.472858 | 9.51E-10 | 1.04E-07 |
|  | *Dapl1* | 2.128923 | 0.5577 | 9.98E-10 | 1.08E-07 |
|  | *Alcam* | 2.403334 | 0.439856 | 1.04E-09 | 1.1E-07 |
|  | *Etfb* | 6.76234 | 0.405274 | 1.04E-09 | 1.1E-07 |
|  | *Hspa8* | 20.8116 | 0.291075 | 1.17E-09 | 1.22E-07 |
|  | *Bambi* | 1.783861 | 0.419422 | 2.06E-09 | 2.1E-07 |
|  | *Pam* | 7.315907 | 0.4546 | 2.36E-09 | 2.33E-07 |
|  | *Insrr* | 5.145543 | 0.481558 | 2.9E-09 | 2.84E-07 |
|  | *Atn1* | 1.288258 | 0.311447 | 4.31E-09 | 4.12E-07 |
|  | *Chic1* | 17.01744 | 0.313832 | 6.57E-09 | 5.91E-07 |
|  | *A330076H08Rik* | 7.556491 | 0.442033 | 7.01E-09 | 6.24E-07 |
|  | *Osgin1* | 1.469992 | 0.336142 | 8.28E-09 | 7.23E-07 |
|  | *Os9* | 10.72439 | 0.427105 | 8.55E-09 | 7.39E-07 |
|  | *Btg2* | 4.594392 | 0.517712 | 9.14E-09 | 7.82E-07 |
|  | *Slc16a10* | 2.944213 | 0.425876 | 1.33E-08 | 1.13E-06 |
|  | *mt-Rnr1* | 144.8379 | 0.20381 | 4.31E-08 | 3.34E-06 |
|  | *Tram1* | 5.25477 | 0.379956 | 6.47E-08 | 4.85E-06 |
|  | *Ang* | 2.598466 | 0.445539 | 1.37E-07 | 9.99E-06 |
|  | *Rbm10* | 4.824942 | 0.406658 | 1.58E-07 | 1.14E-05 |
|  | *Rab34* | 2.565022 | 0.402087 | 2.02E-07 | 1.43E-05 |
|  | *Aplp1* | 10.25162 | 0.424585 | 3.26E-07 | 2.24E-05 |
|  | *Vwa5b2* | 1.8543 | 0.336032 | 4.68E-07 | 3.11E-05 |
|  | *Jam2* | 1.31834 | 0.273371 | 5.69E-07 | 3.73E-05 |
|  | *Ankrd10* | 2.633435 | 0.403961 | 1.11E-06 | 7.05E-05 |
|  | *Fam234a* | 3.275815 | 0.42929 | 1.26E-06 | 7.76E-05 |
|  | *Sephs2* | 2.448041 | 0.413829 | 1.32E-06 | 8.06E-05 |
|  | *Idh3b* | 9.32981 | 0.358929 | 1.4E-06 | 8.5E-05 |
|  | *Canx* | 13.22119 | 0.295095 | 1.47E-06 | 8.88E-05 |
|  | *Hspa1a* | 1.393889 | 0.309718 | 1.66E-06 | 9.89E-05 |
|  | *Pdia3* | 35.21426 | 0.233435 | 1.96E-06 | 0.000114 |
|  | *Fam189b* | 2.26536 | 0.387181 | 2.34E-06 | 0.000134 |
|  | *Ccdc107* | 1.538693 | 0.345069 | 2.59E-06 | 0.000147 |
|  | *Klc4* | 2.162059 | 0.34737 | 2.68E-06 | 0.000151 |
|  | *Serinc1* | 4.676694 | 0.325711 | 3.09E-06 | 0.000173 |
|  | *Ftl1* | 138.1902 | 0.203605 | 3.21E-06 | 0.000179 |
|  | *Cog2* | 3.634217 | 0.400981 | 4.26E-06 | 0.000231 |
|  | *Dnm2* | 6.627017 | 0.370015 | 5E-06 | 0.000267 |
|  | *Etv5* | 1.270122 | 0.226964 | 5.62E-06 | 0.000294 |
|  | *Ppp1r15a* | 1.552304 | 0.305944 | 6.56E-06 | 0.000337 |

**Table S2 Sex-biased expression genes in β cell of healthy 9-mounth-old mice (continued)**

| **Type** | **Gene symbol** | **Basemean** | **log2FoldChange** | **P value** | **P adjust** |
| --- | --- | --- | --- | --- | --- |
| Upregulated genes in male | *Pdx1* | 3.731679 | 0.420911 | 6.72E-06 | 0.000341 |
|  | *Cdk12* | 3.800464 | 0.400797 | 7.04E-06 | 0.000355 |
|  | *Mtss1l* | 1.466982 | 0.275982 | 7.62E-06 | 0.000383 |
|  | *Atp2a3* | 5.836227 | 0.343511 | 9.67E-06 | 0.000475 |
|  | *Slc6a17* | 1.418561 | 0.265054 | 1.02E-05 | 0.0005 |
|  | *Whrn* | 1.254436 | 0.210467 | 1.19E-05 | 0.000567 |
|  | *Spc25* | 12.3631 | 0.409147 | 1.3E-05 | 0.000617 |
|  | *Psma7* | 17.06471 | 0.253853 | 1.31E-05 | 0.000617 |
|  | *Cyth1* | 1.540156 | 0.292005 | 1.44E-05 | 0.000671 |
|  | *Slc39a7* | 8.0343 | 0.358176 | 1.47E-05 | 0.000679 |
|  | *H1f0* | 4.251612 | 0.388111 | 1.63E-05 | 0.000731 |
|  | *Rgs11* | 3.16548 | 0.352962 | 1.91E-05 | 0.000846 |
|  | *Ero1lb* | 44.86386 | 0.250485 | 1.97E-05 | 0.000866 |
|  | *Gabarapl1* | 12.29855 | 0.247214 | 2.31E-05 | 0.001002 |
|  | *Bag2* | 1.872184 | 0.347863 | 2.93E-05 | 0.001235 |
|  | *Spcs2* | 26.10876 | 0.244557 | 2.98E-05 | 0.00124 |
|  | *Pura* | 12.67934 | 0.283311 | 2.99E-05 | 0.00124 |
|  | *Adam22* | 6.362012 | 0.304238 | 3.02E-05 | 0.001246 |
|  | *Hid1* | 3.789477 | 0.328773 | 4.9E-05 | 0.001939 |
|  | *Bhlha15* | 1.721893 | 0.271521 | 5.15E-05 | 0.002029 |
|  | *Bmi1* | 1.379476 | 0.220006 | 5.41E-05 | 0.002107 |
|  | *Prrc2b* | 3.844779 | 0.37311 | 5.86E-05 | 0.002249 |
|  | *Pycr2* | 3.613898 | 0.30614 | 5.9E-05 | 0.002258 |
|  | *Ddost* | 28.78208 | 0.220195 | 6.06E-05 | 0.002308 |
|  | *Gnb2* | 3.847843 | 0.340278 | 6.65E-05 | 0.002524 |
|  | *Ripk4* | 1.364795 | 0.21199 | 6.73E-05 | 0.002542 |
|  | *Ctsa* | 2.317267 | 0.268546 | 7.63E-05 | 0.002833 |
|  | *Sez6l* | 3.242609 | 0.321657 | 8.2E-05 | 0.002994 |
|  | *Atf6* | 1.959441 | 0.280474 | 9.69E-05 | 0.003498 |
|  | *Pappa2* | 1.210377 | 0.210981 | 0.0001 | 0.003609 |
|  | *Pitpnc1* | 1.679984 | 0.281864 | 0.000119 | 0.00424 |
|  | *Nbeal1* | 2.07809 | 0.295525 | 0.000131 | 0.004611 |
|  | *Rsrp1* | 12.97639 | 0.243108 | 0.000146 | 0.005033 |
|  | *Cyld* | 1.706259 | 0.263239 | 0.000146 | 0.005033 |
|  | *Hsd17b12* | 3.048443 | 0.347631 | 0.000179 | 0.006034 |
|  | *Akr1c14* | 1.375663 | 0.20383 | 0.000181 | 0.006063 |
|  | *Ninj1* | 6.45138 | 0.327811 | 0.000192 | 0.00642 |
|  | *Selenop* | 8.496306 | 0.251915 | 0.000201 | 0.006651 |
|  | *Kif12* | 5.398855 | 0.286337 | 0.000216 | 0.006973 |
|  | *Araf* | 3.628543 | 0.338112 | 0.00022 | 0.00707 |
|  | *Hspa13* | 3.820149 | 0.36428 | 0.000222 | 0.00707 |

**Table S2 Sex-biased expression genes in β cell of healthy 9-mounth-old mice (continued)**

| **Type** | **Gene symbol** | **Basemean** | **log2FoldChange** | **P value** | **P adjust** |
| --- | --- | --- | --- | --- | --- |
|  | *Zfp106* | 7.072469 | 0.302547 | 0.000222 | 0.00707 |
| Upregulated genes in male | *Eif4a1* | 8.733011 | 0.264057 | 0.000233 | 0.007364 |
|  | *Rasgrf2* | 1.276145 | 0.209372 | 0.000244 | 0.007618 |
|  | *Dnajb1* | 1.906366 | 0.28915 | 0.000258 | 0.008001 |
|  | *Fbxo2* | 1.740883 | 0.265713 | 0.00026 | 0.008029 |
|  | *Egln2* | 2.373935 | 0.289691 | 0.000274 | 0.008447 |
|  | *Tmed2* | 2.96583 | 0.213275 | 0.000281 | 0.0086 |
|  | *Glrx5* | 7.332819 | 0.324309 | 0.000292 | 0.008866 |
|  | *Tmem65* | 1.833737 | 0.259577 | 0.000297 | 0.008982 |
|  | *Selenom* | 8.857555 | 0.291788 | 0.000319 | 0.009538 |
|  | *Napa* | 11.87379 | 0.277834 | 0.000365 | 0.010682 |
|  | *Abcc8* | 13.26307 | 0.225435 | 0.000397 | 0.011448 |
|  | *Slc4a7* | 3.910181 | 0.307246 | 0.000428 | 0.012228 |
|  | *Depp1* | 1.399767 | 0.240455 | 0.000509 | 0.014297 |
|  | *Rexo2* | 3.733613 | 0.284272 | 0.000511 | 0.014297 |
|  | *Vapa* | 3.966374 | 0.333959 | 0.000521 | 0.014542 |
|  | *Ckb* | 3.998093 | 0.388764 | 0.000564 | 0.015655 |
|  | *Eif4enif1* | 1.324901 | 0.208457 | 0.000616 | 0.016691 |
|  | *Selenot* | 4.263196 | 0.329023 | 0.00062 | 0.016691 |
|  | *Ppp2r3a* | 1.993621 | 0.268894 | 0.00064 | 0.017179 |
|  | *Vcp* | 10.21441 | 0.256897 | 0.000646 | 0.017283 |
|  | *Hnrnpa2b1* | 7.322598 | 0.261052 | 0.000651 | 0.017355 |
|  | *Selenof* | 10.46911 | 0.236021 | 0.000673 | 0.017809 |
|  | *Gsn* | 2.458363 | 0.302143 | 0.000718 | 0.018924 |
|  | *Sgcz* | 1.239187 | 0.204459 | 0.00073 | 0.019176 |
|  | *Ppp1cb* | 3.512301 | 0.287477 | 0.000742 | 0.01933 |
|  | *Cfap20* | 2.653915 | 0.291026 | 0.000751 | 0.019508 |
|  | *Gnao1* | 3.122919 | 0.291806 | 0.000761 | 0.019677 |
|  | *Mafk* | 2.064114 | 0.268566 | 0.000762 | 0.019677 |
|  | *Cst3* | 10.59333 | 0.251411 | 0.000818 | 0.020897 |
|  | *Slc9a3r2* | 1.50467 | 0.244186 | 0.00083 | 0.021022 |
|  | *Selenos* | 13.07888 | 0.264022 | 0.000851 | 0.021356 |
|  | *Ctsd* | 9.87045 | 0.278306 | 0.000859 | 0.021356 |
|  | *Hsp90aa1* | 10.6502 | 0.302299 | 0.000859 | 0.021356 |
|  | *Map4* | 1.389535 | 0.20558 | 0.00086 | 0.021356 |
|  | *Tmem214* | 2.649402 | 0.260827 | 0.000865 | 0.021425 |
|  | *Hnrnpk* | 7.108109 | 0.224462 | 0.000908 | 0.022308 |
|  | *Ddx50* | 3.501408 | 0.264896 | 0.000968 | 0.023503 |
|  | *Nap1l5* | 1.83321 | 0.275756 | 0.001035 | 0.025 |
|  | *Mbnl2* | 4.205779 | 0.258553 | 0.0011 | 0.026428 |
|  | *Syt4* | 4.916439 | 0.297732 | 0.001142 | 0.027294 |

**Table S2 Sex-biased expression genes in β cell of healthy 9-mounth-old mice (continued)**

| **Type** | **Gene symbol** | **Basemean** | **log2FoldChange** | **P value** | **P adjust** |
| --- | --- | --- | --- | --- | --- |
|  | *Nudt3* | 3.972544 | 0.245255 | 0.001157 | 0.027594 |
|  | *Nucb1* | 4.512597 | 0.298824 | 0.001278 | 0.029677 |
| Upregulated genes in male | *Cacybp* | 2.747054 | 0.26361 | 0.001281 | 0.029677 |
|  | *Mmaa* | 3.626497 | 0.302372 | 0.001335 | 0.030686 |
|  | *Chchd10* | 4.527841 | 0.280849 | 0.001368 | 0.031259 |
|  | *Gnmt* | 1.558879 | 0.229586 | 0.00137 | 0.031259 |
|  | *Il6ra* | 7.121038 | 0.325284 | 0.001384 | 0.031376 |
|  | *Gfpt1* | 3.488706 | 0.256477 | 0.001389 | 0.031376 |
|  | *Reep6* | 4.260698 | 0.30746 | 0.001425 | 0.031855 |
|  | *Atp2a2* | 13.86523 | 0.247882 | 0.001446 | 0.032245 |
|  | *Ptbp3* | 2.861037 | 0.289429 | 0.001464 | 0.032489 |
|  | *Tspan5* | 3.169664 | 0.292237 | 0.001469 | 0.032512 |
|  | *Ssbp4* | 1.746062 | 0.252859 | 0.001478 | 0.032645 |
|  | *Rrbp1* | 17.17366 | 0.242406 | 0.001485 | 0.032723 |
|  | *Ap1s1* | 3.564492 | 0.281963 | 0.001501 | 0.032905 |
|  | *Surf4* | 10.87142 | 0.211225 | 0.001512 | 0.033024 |
|  | *Tgoln1* | 6.537247 | 0.263223 | 0.001514 | 0.033024 |
|  | *Ncstn* | 2.352542 | 0.239857 | 0.001524 | 0.033172 |
|  | *Vat1* | 3.355562 | 0.270152 | 0.001568 | 0.033961 |
|  | *Fos* | 3.203143 | 0.353513 | 0.001599 | 0.03455 |
|  | *Epm2aip1* | 1.637066 | 0.22772 | 0.001683 | 0.036102 |
|  | *Atp6v0b* | 10.07541 | 0.236416 | 0.001767 | 0.03764 |
|  | *Tuba4a* | 1.769151 | 0.221161 | 0.001876 | 0.039491 |
|  | *Mlec* | 4.408185 | 0.302941 | 0.001926 | 0.040353 |
|  | *Lrrc58* | 2.872242 | 0.257817 | 0.002001 | 0.041354 |
|  | *Sec16a* | 2.359447 | 0.267035 | 0.002018 | 0.041598 |
|  | *Cltb* | 11.03106 | 0.205728 | 0.00208 | 0.042384 |
|  | *Tmem181b-ps* | 4.560206 | 0.241755 | 0.002096 | 0.042574 |
|  | *Vps37a* | 2.874821 | 0.262093 | 0.002133 | 0.04318 |
|  | *Pnrc1* | 1.747508 | 0.245352 | 0.002206 | 0.04434 |
|  | *Scnn1b* | 2.686974 | 0.239219 | 0.002259 | 0.045024 |
|  | *Edem2* | 12.30276 | 0.255713 | 0.002345 | 0.046231 |
|  | *Fbxo44* | 1.766939 | 0.238752 | 0.002352 | 0.04625 |
|  | *Capzb* | 5.795493 | 0.301606 | 0.00244 | 0.047677 |
|  | *Gstt1* | 2.089619 | 0.284504 | 0.002502 | 0.048697 |
|  | *Ankrd54* | 2.548785 | 0.278351 | 0.002503 | 0.048697 |
| Upregulated genes in female | *Xist* | 2.069689 | -1.80482 | 9.9E-140 | 8.9E-136 |
|  | *Iapp* | 1987.582 | -0.43604 | 3.59E-41 | 4.61E-38 |
|  | *Jup* | 3.867988 | -0.95771 | 1.01E-38 | 1.14E-35 |
|  | *Gcg* | 2.146939 | -1.0171 | 4.98E-37 | 4.48E-34 |
|  | *Enpp2* | 3.502088 | -0.96799 | 3.6E-35 | 2.95E-32 |

**Table S2 Sex-biased expression genes in β cell of healthy 9-mounth-old mice (continued)**

| **Type** | **Gene symbol** | **Basemean** | **log2FoldChange** | **P value** | **P adjust** |
| --- | --- | --- | --- | --- | --- |
| Upregulated genes in female | *Fmo1* | 1.612667 | -0.82033 | 3.91E-28 | 2.71E-25 |
|  | *Cish* | 1.575381 | -0.62942 | 1.46E-23 | 8.2E-21 |
|  | *Matn2* | 1.551672 | -0.60906 | 1.07E-20 | 5.05E-18 |
|  | *Pabpc1* | 8.482537 | -0.81695 | 6.63E-20 | 2.59E-17 |
|  | *Sytl4* | 6.68981 | -0.76035 | 7.33E-19 | 2.74E-16 |
|  | *Prss53* | 22.759 | -0.51851 | 6.45E-18 | 2.32E-15 |
|  | *Trpm5* | 3.307806 | -0.73123 | 5.56E-16 | 1.72E-13 |
|  | *Rplp1* | 10.25181 | -0.613 | 9.17E-16 | 2.75E-13 |
|  | *Rps27a* | 10.73957 | -0.53886 | 3.77E-15 | 1.09E-12 |
|  | *Rps29* | 5.828271 | -0.50234 | 7.45E-15 | 2.09E-12 |
|  | *Rps24* | 5.732397 | -0.50764 | 4.21E-14 | 1.05E-11 |
|  | *Rpl10* | 17.07405 | -0.41654 | 5.73E-14 | 1.39E-11 |
|  | *Itga11* | 1.122069 | -0.24285 | 3.27E-13 | 7.17E-11 |
|  | *Sh3pxd2a* | 4.245573 | -0.66556 | 9.45E-13 | 1.9E-10 |
|  | *Nf1* | 1.49074 | -0.47589 | 2.96E-12 | 5.32E-10 |
|  | *Ssr4* | 14.64574 | -0.46042 | 3.29E-12 | 5.79E-10 |
|  | *Shisal2b* | 1.813605 | -0.57532 | 4.21E-12 | 7.12E-10 |
|  | *Rpl15* | 3.296179 | -0.43194 | 5.98E-12 | 9.78E-10 |
|  | *Rps26* | 4.267271 | -0.60963 | 6.1E-12 | 9.79E-10 |
|  | *Cdh12* | 1.159434 | -0.22964 | 8.18E-12 | 1.27E-09 |
|  | *Rps25* | 3.150816 | -0.47906 | 9.63E-12 | 1.47E-09 |
|  | *Gnb1* | 2.975338 | -0.57788 | 1.07E-11 | 1.6E-09 |
|  | *Rpl27a* | 2.760003 | -0.41489 | 3.74E-11 | 5.42E-09 |
|  | *Grem2* | 1.125982 | -0.23099 | 5.73E-11 | 7.92E-09 |
|  | *Degs1* | 8.081232 | -0.49139 | 1.34E-10 | 1.72E-08 |
|  | *Epb41l4a* | 1.216154 | -0.28225 | 1.39E-10 | 1.76E-08 |
|  | *Chac1* | 1.580361 | -0.4545 | 1.87E-10 | 2.33E-08 |
|  | *Arglu1* | 1.700893 | -0.40318 | 2.35E-10 | 2.89E-08 |
|  | *Rps17* | 3.716417 | -0.46 | 3.19E-10 | 3.87E-08 |
|  | *Isg20* | 7.780975 | -0.62254 | 3.96E-10 | 4.75E-08 |
|  | *Socs2* | 1.63977 | -0.35592 | 4.27E-10 | 5.05E-08 |
|  | *Ppia* | 3.438343 | -0.36101 | 5.45E-10 | 6.2E-08 |
|  | *Zbtb20* | 3.438276 | -0.57422 | 1.53E-09 | 1.58E-07 |
|  | *Rpl21* | 13.94408 | -0.45606 | 2.16E-09 | 2.18E-07 |
|  | *Rpl6* | 9.255973 | -0.42906 | 2.21E-09 | 2.21E-07 |
|  | *Cox7b* | 4.2605 | -0.49751 | 3.87E-09 | 3.74E-07 |
|  | *Meis2* | 3.020561 | -0.4683 | 4.65E-09 | 4.4E-07 |
|  | *Rpl41* | 46.17481 | -0.20668 | 4.88E-09 | 4.57E-07 |
|  | *mt-Nd1* | 79.35959 | -0.24306 | 5.18E-09 | 4.78E-07 |
|  | *Rps21* | 9.220538 | -0.51573 | 5.21E-09 | 4.78E-07 |
|  | *Rpl23* | 15.93708 | -0.47224 | 6.35E-09 | 5.77E-07 |

**Table S2 Sex-biased expression genes in β cell of healthy 9-mounth-old mice (continued)**

| **Type** | **Gene symbol** | **Basemean** | **log2FoldChange** | **P value** | **P adjust** |
| --- | --- | --- | --- | --- | --- |
| Upregulated genes in female | *Prlr* | 17.03999 | -0.42491 | 7.63E-09 | 6.73E-07 |
|  | *Rpl37a* | 15.5674 | -0.32547 | 1.37E-08 | 1.15E-06 |
|  | *Myo1b* | 1.152261 | -0.21063 | 1.39E-08 | 1.15E-06 |
|  | *Prkcb* | 3.525651 | -0.45444 | 1.39E-08 | 1.15E-06 |
|  | *Atf4* | 4.146101 | -0.47884 | 1.46E-08 | 1.19E-06 |
|  | *Tpt1* | 8.554347 | -0.35012 | 1.55E-08 | 1.26E-06 |
|  | *Hsp90ab1* | 23.22874 | -0.3761 | 1.77E-08 | 1.42E-06 |
|  | *Dap* | 11.5771 | -0.41844 | 1.85E-08 | 1.47E-06 |
|  | *Usmg5* | 7.460221 | -0.35743 | 1.9E-08 | 1.5E-06 |
|  | *Rps12* | 4.47772 | -0.57047 | 1.92E-08 | 1.5E-06 |
|  | *Apobec1* | 1.397371 | -0.33257 | 4.63E-08 | 3.56E-06 |
|  | *Abr* | 1.755854 | -0.36378 | 5.66E-08 | 4.31E-06 |
|  | *Cebpd* | 1.233676 | -0.25001 | 5.76E-08 | 4.35E-06 |
|  | *Eif4ebp1* | 1.848493 | -0.43227 | 1.03E-07 | 7.67E-06 |
|  | *4732471J01Rik* | 1.961031 | -0.43262 | 1.16E-07 | 8.57E-06 |
|  | *AW822252* | 1.197701 | -0.23048 | 1.56E-07 | 1.13E-05 |
|  | *Mid1ip1* | 1.982559 | -0.35841 | 1.85E-07 | 1.32E-05 |
|  | *Tceal9* | 27.24593 | -0.28577 | 2.24E-07 | 1.57E-05 |
|  | *Ndufb3* | 2.746535 | -0.34015 | 2.85E-07 | 1.98E-05 |
|  | *Rpl12* | 3.038489 | -0.40192 | 3.02E-07 | 2.09E-05 |
|  | *Rps4x* | 3.776447 | -0.39026 | 4.1E-07 | 2.75E-05 |
|  | *Fmn2* | 2.022932 | -0.40012 | 6.71E-07 | 4.37E-05 |
|  | *Kidins220* | 3.481529 | -0.43117 | 7.09E-07 | 4.58E-05 |
|  | *Cox7a2l* | 5.758991 | -0.35371 | 8.58E-07 | 5.51E-05 |
|  | *Rps13* | 6.217807 | -0.39273 | 1.07E-06 | 6.8E-05 |
|  | *Mt2* | 5.421691 | -0.55743 | 1.12E-06 | 7.06E-05 |
|  | *Rpl38* | 7.355306 | -0.26208 | 1.17E-06 | 7.33E-05 |
|  | *Wdr7* | 1.907427 | -0.33774 | 1.18E-06 | 7.33E-05 |
|  | *Rpl19* | 7.491725 | -0.32338 | 1.66E-06 | 9.89E-05 |
|  | *Vps50* | 1.436317 | -0.25079 | 1.88E-06 | 0.000111 |
|  | *Rps14* | 22.50213 | -0.2616 | 1.91E-06 | 0.000112 |
|  | *Tnfrsf9* | 1.24331 | -0.2277 | 2.07E-06 | 0.00012 |
|  | *2310022B05Rik* | 1.307114 | -0.21172 | 2.21E-06 | 0.000127 |
|  | *Gapdh* | 5.044335 | -0.29905 | 3.53E-06 | 0.000196 |
|  | *Mir682* | 4.59293 | -0.26468 | 3.87E-06 | 0.000213 |
|  | *Rps10* | 2.41851 | -0.31947 | 3.96E-06 | 0.000217 |
|  | *Gpx3* | 1.442772 | -0.3403 | 4.07E-06 | 0.000222 |
|  | *Cox6a2* | 5.091221 | -0.45774 | 4.68E-06 | 0.000252 |
|  | *Rpl22l1* | 4.423108 | -0.37401 | 5.15E-06 | 0.000274 |
|  | *Sec61b* | 29.62933 | -0.24705 | 5.37E-06 | 0.000284 |
|  | *Ovol2* | 1.346758 | -0.2852 | 5.92E-06 | 0.000307 |

**Table S2 Sex-biased expression genes in β cell of healthy 9-mounth-old mice (continued)**

| **Type** | **Gene symbol** | **Basemean** | **log2FoldChange** | **P value** | **P adjust** |
| --- | --- | --- | --- | --- | --- |
| Upregulated genes in female | *D17Wsu92e* | 2.22547 | -0.36155 | 6.72E-06 | 0.000341 |
|  | *Pebp1* | 9.419318 | -0.26383 | 8.6E-06 | 0.00043 |
|  | *Akr1c19* | 2.296222 | -0.40465 | 9.08E-06 | 0.000451 |
|  | *Rps3a1* | 7.906193 | -0.34995 | 1.04E-05 | 0.000501 |
|  | *Tmed3* | 16.88895 | -0.27585 | 1.04E-05 | 0.000501 |
|  | *Aars* | 3.568821 | -0.36516 | 1.05E-05 | 0.000504 |
|  | *Banp* | 2.031939 | -0.32792 | 1.11E-05 | 0.000528 |
|  | *Wnt4* | 2.791629 | -0.39362 | 1.4E-05 | 0.000656 |
|  | *Phactr1* | 6.862169 | -0.3396 | 1.49E-05 | 0.000689 |
|  | *Rpsa* | 7.227187 | -0.39938 | 1.5E-05 | 0.000689 |
|  | *9530091C08Rik* | 2.08403 | -0.34048 | 1.56E-05 | 0.000713 |
|  | *Hist1h2bc* | 5.788111 | -0.39783 | 1.6E-05 | 0.000721 |
|  | *Rps6* | 2.429999 | -0.23509 | 1.79E-05 | 0.000802 |
|  | *Slc25a3* | 11.5787 | -0.34282 | 1.87E-05 | 0.000833 |
|  | *Eif5b* | 5.510171 | -0.39224 | 2.16E-05 | 0.000948 |
|  | *Rps7* | 6.762321 | -0.32873 | 2.29E-05 | 0.001 |
|  | *Arpc5* | 6.227688 | -0.34703 | 2.33E-05 | 0.001005 |
|  | *Col6a3* | 1.37735 | -0.26247 | 2.34E-05 | 0.001006 |
|  | *Naxd* | 3.051484 | -0.39937 | 2.39E-05 | 0.001025 |
|  | *Rpl8* | 16.03025 | -0.32 | 2.42E-05 | 0.001031 |
|  | *Wipi1* | 1.921781 | -0.34462 | 2.55E-05 | 0.00108 |
|  | *Rgs2* | 3.48951 | -0.45226 | 2.97E-05 | 0.00124 |
|  | *Rpl17* | 6.501736 | -0.29885 | 2.99E-05 | 0.00124 |
|  | *Rplp2* | 7.877646 | -0.33094 | 3.2E-05 | 0.001312 |
|  | *Eif3m* | 3.175848 | -0.31967 | 3.4E-05 | 0.001389 |
|  | *Rammet* | 1.999215 | -0.27033 | 3.59E-05 | 0.001461 |
|  | *Tmem106c* | 2.44372 | -0.32673 | 3.63E-05 | 0.001469 |
|  | *Rpl7a* | 2.601503 | -0.30612 | 3.85E-05 | 0.00155 |
|  | *Slc25a4* | 9.755105 | -0.30413 | 4.57E-05 | 0.001834 |
|  | *Rps3* | 6.103378 | -0.31982 | 4.67E-05 | 0.001864 |
|  | *Rps11* | 5.149106 | -0.29823 | 5.45E-05 | 0.002112 |
|  | *Rpl35a* | 5.608116 | -0.29447 | 5.63E-05 | 0.002173 |
|  | *S100a10* | 1.520238 | -0.26263 | 7.05E-05 | 0.002653 |
|  | *Rps28* | 3.271979 | -0.2415 | 7.14E-05 | 0.002675 |
|  | *Rpl32* | 8.738895 | -0.27035 | 7.32E-05 | 0.002729 |
|  | *Rps5* | 8.318153 | -0.30051 | 7.84E-05 | 0.002877 |
|  | *Eef1b2* | 2.128215 | -0.29418 | 8.23E-05 | 0.002994 |
|  | *Ndufa4* | 10.94865 | -0.28862 | 8.35E-05 | 0.003026 |
|  | *Rps15a* | 7.047382 | -0.3973 | 0.000114 | 0.004095 |
|  | *Rpl29* | 2.760162 | -0.26402 | 0.000122 | 0.004335 |
|  | *Ptch1* | 1.400291 | -0.21164 | 0.000127 | 0.004512 |

**Table S2 Sex-biased expression genes in β cell of healthy 9-mounth-old mice (continued)**

| **Type** | **Gene symbol** | **Basemean** | **log2FoldChange** | **P value** | **P adjust** |
| --- | --- | --- | --- | --- | --- |
| Upregulated genes in female | *Uqcrb* | 6.054907 | -0.30446 | 0.000135 | 0.004734 |
|  | *Mthfd1* | 3.210825 | -0.30239 | 0.00014 | 0.004864 |
|  | *Rps20* | 3.911586 | -0.33862 | 0.000157 | 0.005365 |
|  | *Synpr* | 1.833749 | -0.27103 | 0.000162 | 0.005523 |
|  | *2410021H03Rik* | 1.449082 | -0.26632 | 0.000179 | 0.006037 |
|  | *Cyb5a* | 2.96058 | -0.35917 | 0.000196 | 0.006511 |
|  | *Tmigd3* | 1.632957 | -0.261 | 0.000202 | 0.006677 |
|  | *Rpl13* | 13.2429 | -0.2292 | 0.00021 | 0.006894 |
|  | *Elovl6* | 1.599309 | -0.24708 | 0.000214 | 0.006956 |
|  | *Bcl7b* | 2.580466 | -0.32063 | 0.000224 | 0.00712 |
|  | *H3f3a* | 2.058363 | -0.21216 | 0.000235 | 0.007398 |
|  | *Ppig* | 3.110041 | -0.32369 | 0.000236 | 0.007398 |
|  | *Nap1l1* | 3.624707 | -0.30092 | 0.000258 | 0.008001 |
|  | *Csf2ra* | 1.433294 | -0.20896 | 0.000281 | 0.0086 |
|  | *Golga2* | 2.763231 | -0.31705 | 0.000306 | 0.009241 |
|  | *Eif3c* | 3.758235 | -0.30711 | 0.000312 | 0.009357 |
|  | *Rpl13a* | 23.68893 | -0.22938 | 0.000352 | 0.010373 |
|  | *Cfl1* | 8.360829 | -0.24643 | 0.000352 | 0.010373 |
|  | *Rhobtb1* | 2.310173 | -0.28787 | 0.000358 | 0.010525 |
|  | *Mt1* | 13.69757 | -0.38514 | 0.00038 | 0.011053 |
|  | *Ap1s2* | 4.012729 | -0.29681 | 0.000416 | 0.011952 |
|  | *Cox7c* | 8.851752 | -0.27219 | 0.000429 | 0.012228 |
|  | *Phf3* | 1.86287 | -0.25565 | 0.000465 | 0.013216 |
|  | *Mettl7a1* | 1.671109 | -0.24408 | 0.000475 | 0.013457 |
|  | *Rpl10a* | 3.262074 | -0.25915 | 0.000483 | 0.013642 |
|  | *Brk1* | 3.598364 | -0.31335 | 0.000569 | 0.015748 |
|  | *Gnl2* | 1.90547 | -0.24702 | 0.000614 | 0.016691 |
|  | *Fau* | 9.34107 | -0.29535 | 0.00062 | 0.016691 |
|  | *Fkbp9* | 4.105892 | -0.25991 | 0.000674 | 0.017809 |
|  | *Agt* | 1.599174 | -0.24815 | 0.000732 | 0.019176 |
|  | *Nfkbia* | 2.0273 | -0.28688 | 0.000734 | 0.01919 |
|  | *Ncoa6* | 1.782586 | -0.23423 | 0.000777 | 0.019948 |
|  | *Comt* | 2.40408 | -0.28536 | 0.000783 | 0.020056 |
|  | *Atf5* | 4.577901 | -0.359 | 0.000823 | 0.020907 |
|  | *H2afz* | 3.353136 | -0.24555 | 0.000851 | 0.021356 |
|  | *Lamtor3* | 2.036331 | -0.25261 | 0.000859 | 0.021356 |
|  | *1300002E11Rik* | 1.534325 | -0.2164 | 0.000871 | 0.021509 |
|  | *Ociad2* | 6.395835 | -0.27107 | 0.000886 | 0.021815 |
|  | *Anapc13* | 6.453737 | -0.2888 | 0.000937 | 0.022938 |
|  | *Rpgr* | 1.548502 | -0.23185 | 0.000956 | 0.02336 |
|  | *Coa4* | 1.345729 | -0.20116 | 0.00096 | 0.02338 |

**Table S2 Sex-biased expression genes in β cell of healthy 9-mounth-old mice (continued)**

| **Type** | **Gene symbol** | **Basemean** | **log2FoldChange** | **P value** | **P adjust** |
| --- | --- | --- | --- | --- | --- |
| Upregulated genes in female | *Maob* | 3.753138 | -0.30839 | 0.001065 | 0.025658 |
|  | *Rpl36al* | 5.441838 | -0.22242 | 0.001176 | 0.027815 |
|  | *Hint1* | 6.454253 | -0.31508 | 0.001194 | 0.028175 |
|  | *Adck5* | 1.518916 | -0.21103 | 0.001202 | 0.02828 |
|  | *Iars* | 1.782343 | -0.22941 | 0.001286 | 0.029715 |
|  | *Imp3* | 2.289987 | -0.27189 | 0.001386 | 0.031376 |
|  | *Sem1* | 2.641182 | -0.24834 | 0.001394 | 0.03141 |
|  | *Gng5* | 2.862015 | -0.22991 | 0.0014 | 0.031464 |
|  | *Zc2hc1c* | 1.420241 | -0.21131 | 0.001463 | 0.032489 |
|  | *Tomm6* | 3.316552 | -0.22093 | 0.0015 | 0.032905 |
|  | *Bmyc* | 2.568796 | -0.28794 | 0.001546 | 0.033554 |
|  | *Psmb1* | 3.209201 | -0.24571 | 0.001707 | 0.036525 |
|  | *Rabep1* | 2.539841 | -0.26664 | 0.00174 | 0.037151 |
|  | *Trib1* | 1.778807 | -0.23117 | 0.001787 | 0.037972 |
|  | *Chmp2a* | 4.712285 | -0.28331 | 0.001902 | 0.039936 |
|  | *Dhrs4* | 2.375122 | -0.29695 | 0.001964 | 0.04069 |
|  | *Rpl7* | 5.812447 | -0.28705 | 0.002042 | 0.041895 |
|  | *Mien1* | 2.763814 | -0.24687 | 0.002073 | 0.042352 |
|  | *Mafg* | 1.58132 | -0.2141 | 0.0022 | 0.04434 |
|  | *Scgn* | 8.944709 | -0.2829 | 0.002215 | 0.04434 |
|  | *Rpl11* | 9.60773 | -0.24237 | 0.002254 | 0.045024 |
|  | *Zfp516* | 2.81269 | -0.27376 | 0.002304 | 0.045714 |
|  | *Gnptg* | 3.268431 | -0.22534 | 0.002312 | 0.045769 |
|  | *Sltm* | 2.718556 | -0.28499 | 0.002374 | 0.046594 |
|  | *Zc3h3* | 2.336061 | -0.24014 | 0.002388 | 0.04677 |
