## Supplemental table 1 to 9 for "Mouse Single Islet β Cell Transcriptomics Reveal Sexually Dimorphic Transcriptomes and Type 2 Diabetes Genes": Table S3.docx

**Table S3 Sex-biased expression genes in β cell of 9-mounth-old T2D mice**

| **Type** | **Gene symbol** | **Basemean** | **log2FoldChange** | **P value** | **P adjust** |
| --- | --- | --- | --- | --- | --- |
| Upregulated genes in male | *Eif2s3y* | 1.565417 | 1.094034 | 1.90931E-81 | 8.76468E-78 |
|  | *Scg2* | 169.1795 | 0.457586 | 1.62514E-48 | 4.97348E-45 |
|  | *Ddx3y* | 1.276198 | 0.562089 | 9.94143E-44 | 1.82544E-40 |
|  | *Necab2* | 4.042517 | 1.133534 | 8.11421E-38 | 1.24161E-34 |
|  | *Uty* | 1.219289 | 0.463558 | 4.86389E-37 | 6.37935E-34 |
|  | *Naa20* | 1.770286 | 0.439966 | 1.36111E-15 | 8.92597E-13 |
|  | *Rnf5* | 1.982972 | 0.523402 | 2.64295E-15 | 1.61766E-12 |
|  | *Nkx2-2* | 2.410468 | 0.479397 | 1.23916E-13 | 6.6922E-11 |
|  | *Sdf2l1* | 3.862331 | 0.489877 | 1.13486E-12 | 5.78843E-10 |
|  | *Gc* | 2.332014 | 0.577055 | 1.38944E-12 | 6.7139E-10 |
|  | *Tmed2* | 2.888774 | 0.356069 | 3.73759E-12 | 1.71574E-09 |
|  | *Tssc4* | 6.950201 | 0.47455 | 4.75374E-11 | 1.98382E-08 |
|  | *Spp1* | 1.449726 | 0.494258 | 2.80492E-10 | 1.11965E-07 |
|  | *P4hb* | 14.77533 | 0.321963 | 5.17602E-10 | 1.90084E-07 |
|  | *Bambi* | 1.91103 | 0.409253 | 1.54678E-09 | 5.25964E-07 |
|  | *Cd47* | 3.328244 | 0.423413 | 9.47619E-09 | 2.80648E-06 |
|  | *Syp* | 5.164722 | 0.370449 | 1.47407E-08 | 4.10104E-06 |
|  | *Il1r1* | 12.50519 | 0.398575 | 1.88755E-08 | 5.09694E-06 |
|  | *Mrps28* | 1.231113 | 0.213988 | 2.94895E-08 | 7.73551E-06 |
|  | *Rasgrf2* | 1.315226 | 0.280902 | 7.51943E-08 | 1.86583E-05 |
|  | *Paip2* | 8.160833 | 0.330627 | 1.26207E-07 | 2.97103E-05 |
|  | *Atn1* | 1.282771 | 0.241061 | 2.57791E-07 | 5.6352E-05 |
|  | *Hyou1* | 3.048203 | 0.346366 | 2.8753E-07 | 6.1391E-05 |
|  | *Pappa2* | 1.325729 | 0.292922 | 7.20148E-07 | 0.000143732 |
|  | *Meg3* | 23.47778 | 0.284943 | 1.00136E-06 | 0.000195029 |
|  | *Rnase4* | 24.25724 | 0.339201 | 1.20467E-06 | 0.000225715 |
|  | *Ccnd1* | 2.574984 | 0.346763 | 1.4868E-06 | 0.000254122 |
|  | *Kmt2d* | 4.413322 | 0.354876 | 1.60556E-06 | 0.000268011 |
|  | *Acly* | 11.33837 | 0.247448 | 1.96652E-06 | 0.000316748 |
|  | *Tspan33* | 5.578722 | 0.414638 | 2.61063E-06 | 0.000413245 |
|  | *Pdia3* | 30.58145 | 0.227263 | 3.78259E-06 | 0.000563626 |
|  | *Tmbim6* | 5.873785 | 0.287399 | 3.80621E-06 | 0.000563626 |
|  | *Kctd13* | 1.484975 | 0.249898 | 5.1361E-06 | 0.000748485 |
|  | *Nnt* | 1.49006 | 0.24099 | 5.76146E-06 | 0.0008265 |
|  | *Tram1* | 5.606432 | 0.30353 | 5.88621E-06 | 0.000831404 |
|  | *Hap1* | 4.261206 | 0.358741 | 6.82804E-06 | 0.000943314 |
|  | *Slc16a10* | 3.129277 | 0.325063 | 6.88401E-06 | 0.000943314 |
|  | *Dtx3* | 2.307832 | 0.308141 | 7.24885E-06 | 0.000978702 |
|  | *Derl3* | 1.37897 | 0.230422 | 1.21173E-05 | 0.001483325 |
|  | *Selenof* | 9.825785 | 0.276931 | 1.28437E-05 | 0.001551555 |

**Table S3 Sex-biased expression genes in β cell of 9-mounth-old T2D mice (continued)**

| **Type** | **Gene symbol** | **Basemean** | **log2FoldChange** | **P value** | **P adjust** |
| --- | --- | --- | --- | --- | --- |
| Upregulated genes in male | *Dapl1* | 3.094251 | 0.451124 | 1.44059E-05 | 0.001717673 |
|  | *Pdia4* | 7.161075 | 0.326121 | 2.06757E-05 | 0.002402825 |
|  | *Tent5a* | 1.646579 | 0.285574 | 2.34165E-05 | 0.002654159 |
|  | *Itm2c* | 3.896665 | 0.275278 | 3.72649E-05 | 0.0041723 |
|  | *Copb2* | 6.644442 | 0.273293 | 3.83487E-05 | 0.004241916 |
|  | *Manf* | 25.23628 | 0.249507 | 3.95026E-05 | 0.004317539 |
|  | *Syt13* | 23.49679 | 0.225319 | 4.45043E-05 | 0.004751093 |
|  | *Clic4* | 1.596611 | 0.229493 | 5.16439E-05 | 0.005449921 |
|  | *Cst3* | 11.12087 | 0.266013 | 7.00149E-05 | 0.007063807 |
|  | *Pcp4* | 1.916138 | 0.327611 | 8.84604E-05 | 0.008732845 |
|  | *Etfb* | 7.315348 | 0.257408 | 0.000106059 | 0.010218322 |
|  | *Gadd45gip1* | 6.820377 | 0.275969 | 0.00010796 | 0.010218322 |
|  | *Sdhd* | 2.0884 | 0.275772 | 0.000112594 | 0.010548203 |
|  | *Akap8l* | 2.601281 | 0.315681 | 0.000142846 | 0.012984821 |
|  | *Rab3gap2* | 1.472903 | 0.208792 | 0.000161926 | 0.014158524 |
|  | *Unc50* | 1.75063 | 0.212947 | 0.000178106 | 0.015282197 |
|  | *Ptprn* | 8.325304 | 0.312999 | 0.000199691 | 0.016975564 |
|  | *Vegfa* | 1.752899 | 0.227135 | 0.000204052 | 0.017030942 |
|  | *Kif12* | 5.181117 | 0.278108 | 0.000216395 | 0.017898388 |
|  | *Sdc4* | 1.809251 | 0.228347 | 0.00023887 | 0.019407624 |
|  | *Spcs3* | 6.360416 | 0.249628 | 0.00024513 | 0.019719678 |
|  | *Cyb561* | 3.192056 | 0.284948 | 0.000378272 | 0.028007393 |
|  | *Rab34* | 2.469559 | 0.252766 | 0.000400677 | 0.029195395 |
|  | *Fxyd6* | 4.160209 | 0.296018 | 0.000424174 | 0.030188711 |
|  | *Ldlr* | 2.613563 | 0.254162 | 0.000435792 | 0.030639246 |
|  | *Spint2* | 8.66128 | 0.240165 | 0.00043949 | 0.030639246 |
|  | *Dio1* | 3.979043 | 0.300937 | 0.000440516 | 0.030639246 |
|  | *Chic1* | 16.12917 | 0.206386 | 0.000445765 | 0.030771189 |
|  | *Nbas* | 4.752153 | 0.241714 | 0.000474394 | 0.031927683 |
|  | *Sdhc* | 3.191769 | 0.273008 | 0.000485334 | 0.032288801 |
|  | *Rsrp1* | 12.64166 | 0.211453 | 0.000532602 | 0.034679582 |
|  | *1700086L19Rik* | 1.540502 | 0.228366 | 0.000538874 | 0.034840864 |
|  | *Dnajc3* | 19.45073 | 0.205506 | 0.000547084 | 0.034964147 |
|  | *Selenom* | 9.347638 | 0.249914 | 0.000556358 | 0.034964147 |
|  | *H1f0* | 3.587263 | 0.291541 | 0.000557831 | 0.034964147 |
|  | *2900055J20Rik* | 2.526588 | 0.235727 | 0.000559822 | 0.034964147 |
|  | *Sec11c* | 16.56363 | 0.232702 | 0.000576915 | 0.035548018 |
|  | *Abcc8* | 13.96821 | 0.20816 | 0.000604972 | 0.037028323 |
|  | *Hid1* | 4.301314 | 0.268494 | 0.000616242 | 0.0374683 |
|  | *Csad* | 2.03415 | 0.255615 | 0.00068127 | 0.040585916 |
|  | *Rpn1* | 13.91703 | 0.23034 | 0.000712901 | 0.041688833 |

**Table S3 Sex-biased expression genes in β cell of 9-mounth-old T2D mice (continued)**

| **Type** | **Gene symbol** | **Basemean** | **log2FoldChange** | **P value** | **P adjust** |
| --- | --- | --- | --- | --- | --- |
| Upregulated genes in female | *Xist* | 2.153581 | -1.93706 | 2.4578E-160 | 2.2565E-156 |
|  | *Cish* | 1.716479 | -0.91719 | 1.66144E-46 | 3.81341E-43 |
|  | *Enpp2* | 3.210527 | -0.92367 | 2.43364E-36 | 2.7929E-33 |
|  | *Socs2* | 1.616638 | -0.49001 | 4.53281E-21 | 4.62397E-18 |
|  | *G6pc2* | 102.8299 | -0.37441 | 1.09384E-19 | 1.00425E-16 |
|  | *Prlr* | 17.2077 | -0.59562 | 1.6099E-19 | 1.34368E-16 |
|  | *Jup* | 3.764578 | -0.66213 | 7.84175E-19 | 5.99959E-16 |
|  | *Fxyd2* | 1.214431 | -0.44606 | 4.46506E-16 | 3.15336E-13 |
|  | *Gcg* | 2.79143 | -0.67637 | 3.56763E-15 | 2.04715E-12 |
|  | *Cebpd* | 1.255522 | -0.28866 | 8.25282E-10 | 2.9142E-07 |
|  | *Prss53* | 21.08724 | -0.33773 | 2.59813E-09 | 8.51908E-07 |
|  | *Sytl4* | 6.728527 | -0.48697 | 3.9628E-09 | 1.25457E-06 |
|  | *Chga* | 652.1322 | -0.22814 | 8.70067E-09 | 2.6627E-06 |
|  | *Rpl23* | 18.66599 | -0.44043 | 1.39493E-08 | 4.00215E-06 |
|  | *Spc25* | 12.58776 | -0.43854 | 4.87796E-08 | 1.24401E-05 |
|  | *Cd9* | 1.187818 | -0.22565 | 9.16698E-08 | 2.21479E-05 |
|  | *Rpl10* | 17.59557 | -0.27475 | 2.08964E-07 | 4.79624E-05 |
|  | *Sphkap* | 8.868342 | -0.43345 | 2.29666E-07 | 5.14285E-05 |
|  | *Gpx3* | 1.732163 | -0.46512 | 6.61329E-07 | 0.000134926 |
|  | *Galnt9* | 1.266092 | -0.22996 | 1.23661E-06 | 0.000227067 |
|  | *Chst12* | 1.480606 | -0.23741 | 1.3295E-06 | 0.000239336 |
|  | *Hsp90ab1* | 25.31112 | -0.28855 | 1.48284E-06 | 0.000254122 |
|  | *P2ry1* | 1.934401 | -0.31061 | 1.84217E-06 | 0.000302018 |
|  | *Rps14* | 23.28303 | -0.21766 | 3.21328E-06 | 0.000491685 |
|  | *Fmo1* | 1.289827 | -0.24252 | 7.68102E-06 | 0.001022021 |
|  | *Hsbp1* | 5.470287 | -0.34831 | 8.2681E-06 | 0.001063468 |
|  | *Rplp1* | 10.92367 | -0.323 | 8.27804E-06 | 0.001063468 |
|  | *Hnrnpm* | 4.080875 | -0.36314 | 8.39212E-06 | 0.001063468 |
|  | *Ap1s2* | 4.147844 | -0.3482 | 8.45585E-06 | 0.001063468 |
|  | *Akr1c19* | 2.210838 | -0.3333 | 5.90996E-05 | 0.006165831 |
|  | *Fau* | 9.932548 | -0.29834 | 6.35077E-05 | 0.006551282 |
|  | *Cox7a2l* | 6.08677 | -0.27174 | 7.74904E-05 | 0.007733041 |
|  | *Vps35* | 5.180932 | -0.29223 | 0.000101821 | 0.009944876 |
|  | *Rpl21* | 14.69273 | -0.27514 | 0.000147787 | 0.013302299 |
|  | *Peg3* | 3.095583 | -0.30852 | 0.000157523 | 0.014040916 |
|  | *Gcsh* | 1.46553 | -0.20959 | 0.000161255 | 0.014158524 |
|  | *Rlf* | 1.758805 | -0.24176 | 0.000202097 | 0.017022485 |
|  | *Stip1* | 2.542513 | -0.26871 | 0.000258127 | 0.020429826 |
|  | *Gadd45g* | 3.342915 | -0.30432 | 0.000324557 | 0.025176915 |
|  | *Rpl27a* | 2.966972 | -0.22878 | 0.000326332 | 0.025176915 |

**Table S3 Sex-biased expression genes in β cell of 9-mounth-old T2D mice (continued)**

| **Type** | **Gene symbol** | **Basemean** | **log2FoldChange** | **P value** | **P adjust** |
| --- | --- | --- | --- | --- | --- |
| Upregulated genes in female | *Mt1* | 11.11573 | -0.35406 | 0.000349314 | 0.026287329 |
|  | *Imp3* | 2.583336 | -0.30017 | 0.000395114 | 0.029020356 |
|  | *Hist1h2bc* | 5.597822 | -0.30283 | 0.000404354 | 0.029231304 |
|  | *Rpl13a* | 25.5201 | -0.20481 | 0.000419024 | 0.030055177 |
|  | *Rgs2* | 6.067359 | -0.39791 | 0.000455068 | 0.031178943 |
|  | *Echdc2* | 2.415706 | -0.29484 | 0.000469772 | 0.031927683 |
|  | *Eif3m* | 3.160661 | -0.25902 | 0.000490587 | 0.032403449 |
|  | *Slc2a2* | 11.02216 | -0.26252 | 0.000553756 | 0.034964147 |
|  | *Zbtb20* | 3.904341 | -0.32015 | 0.0006852 | 0.040585916 |
|  | *Uba52* | 7.592675 | -0.20725 | 0.000702126 | 0.04132193 |
|  | *Psma6* | 3.12481 | -0.26667 | 0.000722263 | 0.041968939 |
|  | *2210016F16Rik* | 2.860618 | -0.27711 | 0.000788464 | 0.044684471 |
