## Supplemental table 1 to 9 for "Mouse Single Islet β Cell Transcriptomics Reveal Sexually Dimorphic Transcriptomes and Type 2 Diabetes Genes": Table S4.docx

**Table S4 Sex biased genes involved in GSEA**

| **Type** | **GSEA** | **Overlapped genes** |
| --- | --- | --- |
| Healthy (8W), male-up | Longevity regulating pathway - multiple species | *Ins1,* *Hspa8* \| |
| Healthy (8W), female-up | Ribosome | *Rps8, Rps13, Rpl21, Rps10, Rpl27a, Rplp1, Rps21, Rpl32, Rpl39, Rpl23, Rps28, Rps5, Rpl9, Rpl7a, Rpl22l1, Rpl23a, Rps24, Rps3a1, Rps12, Rpl5* |
| Healthy (9M), male-up | Protein processing in endoplasmic reticulum | *Rnf5, Hspa8, Os9, Tram1, Canx, Hspa1a, Pdia3, Ppp1r15a, Ero1lb, Bag2, Ddost, Atf6, Dnajb1, Fbxo2, Vcp, Hsp90aa1, Rrbp1, Edem2* |
|  | Longevity regulating pathway - multiple species | *Ins2, Hspa8, Hspa1a* |
| Healthy (9M), female-up | Oxidative phosphorylation | *Cox7c, Uqcrb, Ndufa4, Cox6a2, Cox7a2l, Ndufb3, mt-Nd1, Cox7b* |
|  | Ribosome | *Rpl11, Rpl7, Rpl36al, Fau, Rpl10a, Rpl13a, Rpl13, Rps20, Rpl29, Rps15a, Rps5, Rpl32, Rps28, Rpl35a, Rps11, Rps3, Rpl7a, Rplp2, Rpl17, Rpl8, Rps7, Rps6, Rpsa, Rps3a1, Rpl22l1, Rps10, Rps14, Rpl19, Rpl38, Rps13, Rps4x, Rpl12, Rps12, Rpl37a, Rpl23, Rps21, Rpl41, Rpl6, Rpl21, Rps17, Rpl27a, Rps25, Rps26, Rpl15, Rpl10, Rps24, Rps29, Rps27a, Rplp1* |
| T2D (9M), male-up | Notch signaling pathway | *Dtx3* |
| T2D (9M), male-up | Ribosome | *Uba52, Rpl13a, Rpl27a, Rpl21, Fau, Rplp1, Rps14, Rpl10, Rpl23* |
|  | Carbohydrate digestion and absorption | *Slc2a2, Fxyd2, G6pc2* |
|  | JAK-STAT signaling pathway | *Prlr, Socs2, Cish* |

*Note*: 8W, 8-week-old; 9M, 9-month-old; T2D, type 2 diabetes; male-up, male upregulation; female-up, female upregulation.
