## Supplemental table 1 to 9 for "Mouse Single Islet β Cell Transcriptomics Reveal Sexually Dimorphic Transcriptomes and Type 2 Diabetes Genes": Table S5 .docx

**Table S5 Sex-dependent T2D altered genes**

| **Group** | **Gene symbol** | **Basemean** | **log2FoldChange** | **Ci.lo** | **Ci.hi** | **P value** | **P adjust** |
| --- | --- | --- | --- | --- | --- | --- | --- |
| Female-dependent T2D upregulated genes | *Fxyd2* | 1.2175402 | 0.47390801 | 0.35667882 | 0.59113721 | 7.43E-14 | 3.32E-10 |
|  | *Cpe* | 42.7906513 | 0.3727489 | 0.26913045 | 0.47636736 | 3.33E-11 | 4.00E-08 |
|  | *Serp1* | 17.1086461 | 0.40687078 | 0.28719847 | 0.52654309 | 4.14E-10 | 3.36E-07 |
|  | *Scg3* | 36.3074537 | 0.26298399 | 0.17759765 | 0.34837033 | 1.82E-08 | 1.11E-05 |
|  | *Cd9* | 1.16845523 | 0.25612233 | 0.17249696 | 0.33974769 | 2.25E-08 | 1.26E-05 |
|  | *Igfbp7* | 1.253473 | 0.24993948 | 0.16431618 | 0.33556278 | 1.10E-07 | 5.15E-05 |
|  | *Ttr* | 44.7918192 | 0.34848207 | 0.22288788 | 0.47407626 | 4.98E-07 | 0.00020227 |
|  | *Rnase4* | 18.9575602 | 0.37715139 | 0.23692381 | 0.51737897 | 1.18E-06 | 0.00042317 |
|  | *Stip1* | 2.44457685 | 0.34321544 | 0.20725686 | 0.47917402 | 5.89E-06 | 0.00154785 |
|  | *Ero1lb* | 45.0570084 | 0.27707924 | 0.16454645 | 0.38961203 | 1.04E-05 | 0.00252074 |
|  | *Pnrc1* | 1.79107932 | 0.33538148 | 0.19337411 | 0.47738886 | 2.60E-05 | 0.00514197 |
|  | *Tmem160* | 5.71248153 | 0.38975171 | 0.22358266 | 0.55592075 | 2.99E-05 | 0.0056936 |
|  | *Etv1* | 1.27579596 | 0.22790115 | 0.13022707 | 0.32557523 | 3.33E-05 | 0.00584304 |
|  | *Cmas* | 1.50018853 | 0.27775853 | 0.15641567 | 0.39910138 | 4.89E-05 | 0.00824462 |
|  | *Aplp1* | 9.90696619 | 0.33972116 | 0.18499668 | 0.49444564 | 0.00010707 | 0.01428551 |
|  | *Pcsk1n* | 62.3732783 | 0.21404292 | 0.11623925 | 0.31184659 | 0.00011278 | 0.01477615 |
|  | *Akap13* | 3.94333349 | 0.34797428 | 0.18894028 | 0.50700828 | 0.00011406 | 0.01477615 |
|  | *Manf* | 20.6751814 | 0.27021553 | 0.14383686 | 0.3965942 | 0.0001707 | 0.01883807 |
|  | *Vcp* | 10.2487559 | 0.29426638 | 0.15582381 | 0.43270895 | 0.00018905 | 0.01988152 |
|  | *Mbnl2* | 4.20717368 | 0.28883023 | 0.15243643 | 0.42522403 | 0.00020153 | 0.02070678 |
|  | *Senp5* | 1.44106724 | 0.21588352 | 0.11340618 | 0.31836087 | 0.00021996 | 0.0218354 |
|  | *Depp1* | 1.3859847 | 0.23469273 | 0.11784015 | 0.3515453 | 0.00047037 | 0.04204626 |
| Female-dependent T2D downregulated genes | *Shisal2b* | 1.92446287 | -0.413057743 | -0.5765914 | -0.2495241 | 5.81E-06 | 0.00154785 |
|  | *Chac1* | 1.63103931 | -0.34532728 | -0.485971 | -0.2046835 | 1.12E-05 | 0.00256342 |
|  | *Rpl37a* | 15.8322986 | -0.235898521 | -0.3411162 | -0.1306808 | 7.26E-05 | 0.01100577 |
|  | *Fkbp9* | 3.99751392 | -0.291070998 | -0.4256536 | -0.1564884 | 0.00014019 | 0.01765014 |
|  | *Mid1ip1* | 2.02274785 | -0.286221766 | -0.4186968 | -0.1537467 | 0.00014276 | 0.01772413 |
|  | *Scn9a* | 1.45012408 | -0.224455067 | -0.329701 | -0.1192091 | 0.00017878 | 0.01925452 |
|  | *Rps24* | 6.19606055 | -0.270680485 | -0.3992564 | -0.1421046 | 0.00022229 | 0.0218354 |
|  | *Rps18* | 10.2233094 | -0.231894684 | -0.3463551 | -0.1174343 | 0.0004111 | 0.03711925 |
|  | *Syp* | 4.8590781 | -0.268784244 | -0.4040368 | -0.1335316 | 0.00055134 | 0.0478489 |
| Male-dependent T2D upregulated genes | *Iapp* | 1895.74737 | 0.30829159 | 0.26271489 | 0.3538683 | 2.56E-36 | 2.35E-32 |
|  | *Trpm5* | 2.9783659 | 0.39723356 | 0.24742511 | 0.54704201 | 1.65E-06 | 0.00033652 |
|  | *Prkcb* | 3.37771596 | 0.3187035 | 0.19289546 | 0.44451154 | 5.20E-06 | 0.00097502 |
|  | *Gmds* | 2.24645786 | 0.3156415 | 0.18935518 | 0.44192782 | 7.16E-06 | 0.00129017 |
|  | *Nf1* | 1.34918722 | 0.22415671 | 0.13180642 | 0.31650699 | 1.40E-05 | 0.00221008 |
|  | *Arglu1* | 1.6128599 | 0.24156022 | 0.14107557 | 0.34204488 | 1.72E-05 | 0.00259588 |
|  | *Gnb1* | 2.7056564 | 0.31762751 | 0.18361082 | 0.4516442 | 2.34E-05 | 0.00335303 |
|  | *Tmed3* | 16.7818662 | 0.21867459 | 0.12267994 | 0.31466924 | 5.21E-05 | 0.0062995 |
|  | *Rps13* | 6.03795469 | 0.28277857 | 0.15674638 | 0.40881075 | 7.00E-05 | 0.00834347 |

**Table S5 Sex-dependent T2D altered genes (continued)**

*Note:* Ci.lo, low bound of confidence interval; Ci.hi, high bound of confidence interval.

| **Group** | **Gene symbol** | **Basemean** | **log2FoldChange** | **Ci.lo** | **Ci.hi** | **P value** | **P adjust** |
| --- | --- | --- | --- | --- | --- | --- | --- |
| Male-dependent T2D upregulated genes | *9530091C08Rik* | 2.04870495 | 0.2791375 | 0.15327062 | 0.40500438 | 8.71E-05 | 0.00975206 |
|  | *Sh3pxd2a* | 3.90795254 | 0.32469702 | 0.1777295 | 0.47166454 | 9.34E-05 | 0.00997257 |
|  | *Rpl8* | 15.7823211 | 0.24476238 | 0.12725129 | 0.36227348 | 0.00025962 | 0.02091323 |
|  | *Rps27a* | 9.81673774 | 0.22990867 | 0.1175698 | 0.34224753 | 0.00034479 | 0.02619648 |
|  | *Rpl36al* | 5.52900364 | 0.21809403 | 0.11152322 | 0.32466485 | 0.00034518 | 0.02619648 |
|  | *Tmem63b* | 1.65109857 | 0.21514814 | 0.10775452 | 0.32254176 | 0.0004804 | 0.03267755 |
|  | *Rpl12* | 2.90298883 | 0.2529606 | 0.12554574 | 0.38037547 | 0.0005503 | 0.03635518 |
|  | *Fmn2* | 1.91944885 | 0.24688958 | 0.12061448 | 0.37316468 | 0.00068905 | 0.04275366 |
|  | *Rpl7a* | 2.58285608 | 0.22401976 | 0.10878567 | 0.33925385 | 0.00074794 | 0.04578909 |
| Male-dependent T2D downregulated genes | *Cox4i1* | 75.0408244 | -0.2902051 | -0.3556996 | -0.2247107 | 1.50E-16 | 2.75E-13 |
|  | *Malat1* | 1398.581 | -0.2149183 | -0.2645802 | -0.1652565 | 6.31E-16 | 9.66E-13 |
|  | *Ndufb7* | 15.1387322 | -0.3059034 | -0.402897 | -0.2089098 | 7.22E-09 | 3.32E-06 |
|  | *Ucn3* | 59.2002052 | -0.2626952 | -0.3500545 | -0.1753358 | 3.84E-08 | 1.47E-05 |
|  | *Ubc* | 22.8922403 | -0.2442571 | -0.3335771 | -0.1549371 | 7.11E-07 | 0.00017648 |
|  | *Pura* | 13.0228932 | -0.2643529 | -0.3738149 | -0.1548908 | 1.56E-05 | 0.00238303 |
|  | *Ivd* | 2.45318031 | -0.3225567 | -0.459062 | -0.1860515 | 2.49E-05 | 0.0034635 |
|  | *Gnmt* | 1.55539103 | -0.2567447 | -0.3687635 | -0.1447259 | 4.64E-05 | 0.00567511 |
|  | *Selenow* | 6.00876743 | -0.2776662 | -0.4071795 | -0.1481528 | 0.0001597 | 0.01481343 |
|  | *Zfyve27* | 1.54320634 | -0.2091412 | -0.3069145 | -0.1113679 | 0.0001662 | 0.01509519 |
|  | *H3f3b* | 8.3079373 | -0.2487028 | -0.3653323 | -0.1320733 | 0.00017516 | 0.01546644 |
|  | *Cct7* | 5.03374023 | -0.2595018 | -0.3847644 | -0.1342392 | 0.00028382 | 0.02266362 |
|  | *Cdk12* | 3.99928737 | -0.2947627 | -0.4396488 | -0.1498765 | 0.00037889 | 0.02783452 |
