## Supplemental table 1 to 9 for "Mouse Single Islet β Cell Transcriptomics Reveal Sexually Dimorphic Transcriptomes and Type 2 Diabetes Genes": Table S6 .docx

**Table S6 Sex-independent T2D altered genes**

| **Type** | **Gene symbol** | **Basemean** | **log2FoldChange** | **P value** | **P adjust** |
| --- | --- | --- | --- | --- | --- |
| sex-independent T2D upregulation | *Rgs2* | 6.9816331058545 | 0.780858694999606 | 2.96521125197364e-27 | 2.69774919704562e-23 |
|  | *Dapl1* | 2.58940829593874 | 0.52406411491305 | 2.16964208638028e-15 | 3.94788074037756e-12 |
|  | *Gcg* | 3.7609257963806 | 0.409930156303977 | 1.16616301258261e-12 | 1.32621888605958e-09 |
|  | *Fh1* | 2.95205923722246 | 0.356861143315339 | 4.54957273852882e-12 | 4.13920127751352e-09 |
|  | *Dnajb9* | 7.01075565808392 | 0.323539459159422 | 9.40046103137375e-12 | 7.12711620528653e-09 |
|  | *Pde1c* | 1.49882514607402 | 0.22121811430468 | 6.7577900125076e-11 | 4.39159810955672e-08 |
|  | *Rap1gap2* | 4.49275082648007 | 0.339939649934038 | 5.76485212523587e-09 | 2.28037498414765e-06 |
|  | *Gng4* | 3.35099319579076 | 0.295576860571475 | 9.17373245822594e-09 | 3.46369092840442e-06 |
|  | *Tmem39a* | 2.26075326627326 | 0.253299236112497 | 1.14936577859308e-08 | 4.02189609755378e-06 |
|  | *Pcbp1* | 4.71120033084496 | 0.344864794710894 | 1.27805815240305e-08 | 4.30658261872701e-06 |
|  | *Degs1* | 11.0152345181038 | 0.286595228927192 | 1.42203509986473e-08 | 4.62059833520332e-06 |
|  | *Pabpc1* | 12.9422354279682 | 0.354197685550903 | 2.62274937836708e-08 | 8.22819787737368e-06 |
|  | *Wnt4* | 3.74989960661229 | 0.333264603111104 | 4.67629428178664e-08 | 1.37241694760306e-05 |
|  | *Creld2* | 8.80748803935068 | 0.341375728420232 | 5.19101464608054e-08 | 1.43114700757699e-05 |
|  | *Hsp90aa1* | 13.1606195286397 | 0.29802341487722 | 7.33917765879772e-08 | 1.96387759822769e-05 |
|  | *Hars* | 2.19472402959625 | 0.270854858963729 | 8.50383357329002e-08 | 2.21051079570836e-05 |
|  | *Bhlha15* | 1.95874494176609 | 0.248460989728681 | 1.24140908414234e-07 | 3.05252428311541e-05 |
|  | *Dpagt1* | 1.62756620600451 | 0.212566469647577 | 1.66551632418312e-07 | 3.88535064549179e-05 |
|  | *Ubr4* | 4.97351243231987 | 0.308925013796021 | 2.08150419545053e-07 | 4.73438129255224e-05 |
|  | *H2afy* | 3.78589351196745 | 0.307656433073726 | 3.00733531400057e-07 | 6.51446111589933e-05 |
|  | *Ccnd1* | 2.20764632216373 | 0.25362544746402 | 3.42535543696531e-07 | 7.24741482918846e-05 |
|  | *Srm* | 1.73185463092603 | 0.204228820414266 | 4.7723400130422e-07 | 9.64861098636844e-05 |
|  | *Pcp4* | 1.67125703386228 | 0.271585105836723 | 6.04292788552213e-07 | 0.000118558935308954 |
|  | *Hsph1* | 2.23247231604952 | 0.243804642546483 | 9.74330105774749e-07 | 0.000180907251068136 |
|  | *Necab2* | 2.58377670540169 | 0.298136415537267 | 2.08495723951844e-06 | 0.000338731088663193 |
|  | *Mgat2* | 2.80478275578484 | 0.274239513185263 | 2.4944426793689e-06 | 0.000398148061349093 |
|  | *Dcaf12l1* | 2.92363214308799 | 0.271992623659289 | 2.66736373146612e-06 | 0.000418408193601358 |
|  | *Srsf2* | 3.09766150600146 | 0.280199856651777 | 2.83318850880939e-06 | 0.000429605817552464 |
|  | *Bag1* | 1.65799526711651 | 0.21530758390474 | 4.51687399299642e-06 | 0.000624250506592323 |
|  | *Gpd2* | 3.4334156247849 | 0.24345497774764 | 6.14159146242679e-06 | 0.000786988720072661 |
|  | *Pde3b* | 2.02725987889932 | 0.200242069351501 | 6.96310549583447e-06 | 0.000874836510006432 |
|  | *Gc* | 1.79567619201189 | 0.256335606933973 | 7.2888012233103e-06 | 0.000896128561211853 |
|  | *Gpx3* | 1.99283002430055 | 0.261710847501695 | 1.10702091199166e-05 | 0.00129124054580771 |
|  | *Ubxn4* | 11.4302354003128 | 0.21000820007059 | 1.12602338402751e-05 | 0.00129677984150409 |
|  | *Eif3b* | 2.42537969365035 | 0.248706160166453 | 1.31273799516202e-05 | 0.00147448028147951 |
|  | *Syt4* | 5.47787911684577 | 0.257427627281641 | 1.56160661329127e-05 | 0.00171174662261734 |
|  | *Il1r1* | 10.4698688869013 | 0.23056354559978 | 1.67677548250524e-05 | 0.00179474156939208 |
|  | *Ang* | 2.79498304293482 | 0.247319375078127 | 2.31625657965539e-05 | 0.00236778678221402 |
|  | *Mrps12* | 5.711173213115 | 0.260890590157112 | 2.39213821451175e-05 | 0.00237021875262718 |
|  | *Kcnq1ot1* | 7.31328115471196 | 0.254896734628176 | 2.44756021278304e-05 | 0.00239439815224732 |
|  | *Pdyn* | 1.85111447145866 | 0.271728817181959 | 2.53621393523578e-05 | 0.0024547313173165 |
|  | *Itpr2* | 1.83419787599566 | 0.221274422524036 | 3.08752060193049e-05 | 0.00289590334395501 |

**Table S6 Sex-independent T2D altered genes (continued)**

| **Type** | **Gene symbol** | **Basemean** | **log2FoldChange** | **P value** | **P adjust** |
| --- | --- | --- | --- | --- | --- |
| sex-independent T2D upregulation | *Map7d2* | 1.91849502719905 | 0.213523791124913 | 3.45468025427161e-05 | 0.00311194860924388 |
|  | *Erp29* | 3.75068042320457 | 0.231476187579025 | 7.67526436348496e-05 | 0.00626074763771177 |
|  | *Hdlbp* | 7.46817083535434 | 0.20383281148631 | 7.87065728240058e-05 | 0.00633692388984783 |
|  | *Atp1a1* | 5.68984092180349 | 0.223232626181102 | 8.48148698829828e-05 | 0.00670996248865546 |
|  | *Tagln2* | 3.83665246078073 | 0.235305873188828 | 0.000104807043959894 | 0.00808080072836538 |
|  | *Chchd10* | 5.0050760291632 | 0.223992689153653 | 0.000111761899148488 | 0.00847341465377456 |
|  | *Rpl23* | 21.8354873006807 | 0.215808293088759 | 0.000182396600664777 | 0.0124770246078808 |
|  | *Rpsa* | 9.17026572440106 | 0.227069811507736 | 0.000187798497968625 | 0.0126562276631004 |
|  | *Luc7l2* | 3.47727786142507 | 0.250265471660691 | 0.000195078342563782 | 0.0129953266326328 |
|  | *Fxyd6* | 3.53488748657835 | 0.214029400814538 | 0.000388030498033761 | 0.0229240355266958 |
|  | *C1qa* | 2.48995937572166 | 0.233977667190872 | 0.000466435851819305 | 0.0263579713034288 |
|  | *Isg20* | 10.1047865244135 | 0.241624392741444 | 0.000571832533802979 | 0.030967454717497 |
|  | *Purb* | 3.64656270868576 | 0.205791663607072 | 0.000583398294380333 | 0.0310395186097793 |
|  | *Itpa* | 2.71055731737933 | 0.210361605010901 | 0.000844969709431158 | 0.0413308301957241 |
|  | *Cdk11b* | 4.50267730581717 | 0.229556024998353 | 0.000944139571581391 | 0.0447384469908724 |
|  | *Psmd7* | 3.84140105408243 | 0.225219699291095 | 0.00102259891513982 | 0.046740342022844 |
| sex-independent T2D downregulation | *mt-Rnr1* | 116.945950273556 | -0.248751993767574 | 9.87165043447254e-24 | 4.49061378264156e-20 |
|  | *Matn2* | 1.40120374575314 | -0.318969848606198 | 7.01921098222637e-18 | 2.12869271720985e-14 |
|  | *Slc30a8* | 54.6025881556561 | -0.242440157528732 | 1.92581941406665e-15 | 3.94788074037756e-12 |
|  | *Cox6a2* | 4.16395332079213 | -0.467249321714078 | 5.28100252505983e-13 | 6.86379442471347e-10 |
|  | *mt-Cytb* | 78.209798821151 | -0.212301248952143 | 1.90692891664342e-12 | 1.92769325373575e-09 |
|  | *Cirbp* | 5.09974970149149 | -0.35611569864689 | 3.4325163250258e-11 | 2.40223334808344e-08 |
|  | *Fmo1* | 1.55434892653653 | -0.290740509656892 | 8.63622457859223e-11 | 5.2381580810688e-08 |
|  | *Gnai2* | 20.5064120526281 | -0.248437405139414 | 1.57394269772773e-10 | 8.94983166495431e-08 |
|  | *Pdia3* | 28.15485155815 | -0.211873662520481 | 1.79836377910796e-10 | 9.62441980136716e-08 |
|  | *Fos* | 2.16374593753948 | -0.429554708309344 | 2.04255223593814e-10 | 1.03239668014251e-07 |
|  | *Slc2a2* | 11.7762184921097 | -0.317094553330533 | 4.25204277401457e-10 | 2.03605711357814e-07 |
|  | *Btg2* | 3.19818186977009 | -0.364822377995686 | 6.38313533231332e-10 | 2.90368826266933e-07 |
|  | *Dusp1* | 1.50645450572233 | -0.287645590754734 | 4.83680239333754e-08 | 1.37516338045578e-05 |
|  | *Rps19* | 11.7068749962683 | -0.227913836950578 | 2.31913037869906e-07 | 5.14620687448879e-05 |
|  | *Fkbp1b* | 5.85402063046416 | -0.329495284609806 | 6.12471967412712e-07 | 0.000118558935308954 |
|  | *Creg1* | 8.93504595617999 | -0.266559226110427 | 7.11784406689397e-07 | 0.000134912802751253 |
|  | *Nipal3* | 7.37903208080561 | -0.22739954496069 | 1.02276344941417e-06 | 0.000186102037255402 |
|  | *Ndufs2* | 6.99730878886207 | -0.315880668294561 | 2.02806354708443e-06 | 0.000335478584570439 |
|  | *Gmpr* | 12.3519809733779 | -0.217911615586837 | 3.14753006607287e-06 | 0.000469446369526737 |
|  | *Tmem215* | 5.87889875324772 | -0.255268771348545 | 4.10595398817425e-06 | 0.000592951894990624 |
|  | *Calm2* | 5.73461633841649 | -0.236913292225279 | 4.27914639646855e-06 | 0.000608307404922982 |
|  | *Ncoa1* | 5.57716741371828 | -0.305769071221933 | 5.07959485009938e-06 | 0.000689763491734391 |
|  | *Tmem181b-ps* | 3.61097434761685 | -0.227967152982077 | 5.16232863475472e-06 | 0.000690689204691153 |
|  | *Swi5* | 9.61750183794156 | -0.231452239656315 | 5.3439988626409e-06 | 0.00070463335727981 |
|  | *Ier2* | 1.35205039325976 | -0.22411848172211 | 7.01946199499555e-06 | 0.000874836510006432 |
|  | *Mt2* | 4.87909705253229 | -0.343791294429716 | 8.13208484832939e-06 | 0.000973496157238168 |

**Table S6 Sex-independent T2D altered genes (continued)**

| **Type** | **Gene symbol** | **Basemean** | **log2FoldChange** | **P value** | **P adjust** |
| --- | --- | --- | --- | --- | --- |
| sex-independent T2D downregulation | *Sphkap* | 9.46311143780293 | -0.266647877707843 | 1.58798341754086e-05 | 0.00171993727771271 |
|  | *Mt1* | 12.6362251496949 | -0.300537149376481 | 1.8951049956351e-05 | 0.00200484479654513 |
|  | *H1f0* | 3.17484580898279 | -0.255071263207213 | 2.04459252705616e-05 | 0.0021381267599031 |
|  | *Sfrp5* | 2.01279093462017 | -0.335367468978035 | 3.00577304516936e-05 | 0.00284859616301571 |
|  | *Cars* | 1.80079228803627 | -0.210014681051699 | 3.42194210765659e-05 | 0.00311194860924388 |
|  | *Pcx* | 2.28445767698342 | -0.213214552229174 | 8.25082356732544e-05 | 0.00658473621188832 |
|  | *Fam210b* | 2.59624159871681 | -0.216900587631748 | 0.000113250902750606 | 0.00851534473739683 |
|  | *Cd81* | 2.09865966535751 | -0.273119525135886 | 0.000151390146518032 | 0.010760527757977 |
|  | *A330076H08Rik* | 6.04542433713011 | -0.204936618065806 | 0.000181128975621244 | 0.0124770246078808 |
|  | *Taok2* | 2.59232443630769 | -0.223455995998973 | 0.000218232150623013 | 0.0142840007652386 |
|  | *Gjd2* | 2.91216427472146 | -0.217763944913421 | 0.000311077594804451 | 0.018867893050206 |
|  | *Nagk* | 3.3131072396673 | -0.219244518265524 | 0.000519923045362772 | 0.0286682416164273 |
|  | *Jun* | 1.97169325974681 | -0.232371720106526 | 0.000689319364392244 | 0.0348412643180035 |
|  | *Mthfd2* | 2.60096619873337 | -0.208769709512769 | 0.00098994930373649 | 0.0461874808481774 |
