## Supplemental table 1 to 9 for "Mouse Single Islet β Cell Transcriptomics Reveal Sexually Dimorphic Transcriptomes and Type 2 Diabetes Genes": Table S7.docx

| **Gene symbol** | **Forward primer 5' to 3'** | **Reverse primer 5' to 3'** |
| --- | --- | --- |
| *Cox4i1* | ATTGGCAAGAGAGCCATTTCTAC | CACGCCGATCAGCGTAAGT |
| *Cpe* | CAGCAAGAGGACGGCATCTC | GTCCAACCGCCTCATTACCAT |
| *Ero1lb* | ACGCTAAGTAACGAAAGCAAAGA | CAGCAGGTCCACATACTGTGC |
| *Iapp* | CCACTTGAGAGCTACACCTGT | GAACCAAAAAGTTTGCCAGGC |
| *Malat1* | GGCGGAATTGCTGGTAGTTT | AGCATAGCAGTACACGCCTT |
| *Ndufb7* | CGGCGCTATCTCTGGGATG | TGGGCACAGTAGTCACGTTG |
| *Pura* | AGTACGGCGTGTTTATGCGAG | AACTTGGCCCACACCTTGTAG |
| *Rnase4* | CAGTACCACCAATATCCTGTGC | GCTGGTTCTTGCCCTGTATCTA |
| *Rpl37a* | GCTAAACGCACCAAGAAGG | CCACCGGCCACTGTTTTCAT |
| *Scg3* | CCGGCTCTTGGATACTGGTG | CCTTCGGGTTTGGGGAAAG |
| *Sdha* | GGAACACTCCAAAAACAGACCT | CCACCACTGGGTATTGAGTAGAA |
| *Serp1* | GCAACGTCGCTAAGACCTC | CATGCCCATCCTGATACTTTGAA |
| *Tmed3* | AGCAGGGCGTGAAGTTCTC | CTTCCACATAGCAGTCCACATC |
| *Ttr* | CACCAAATCGTACTGGAAGACA | GTCGTTGGCTGTGAAAACCAC |
| *Ucn3* | AAGCCTCTCCCACAAGTTCTA | GAGGTGCGTTTGGTTGTCATC |
| *Ins2* | AGCGTGGCTTCTTCTACACA | TGCCAAGGTCTGAAGGTCAC |

**Table S7 Primers for qPCR**
