## Supplemental table 1 to 9 for "Mouse Single Islet β Cell Transcriptomics Reveal Sexually Dimorphic Transcriptomes and Type 2 Diabetes Genes": Table S8.docx

**Table S8 Primers for scRNA-seq library construction**

| **Primer_ID** | **Sequence 5’ to 3’** | **Barcode/index** | | **Note** |
| --- | --- | --- | --- | --- |
| ISPCR | AAGCAGTGGTATCAACGCAGAGT | |  | cDNA-amplification |
| ISPCR-read2 | AAGCAGTGGTATCAACGCAGAGTCAGACGTGTGCTCTTCCGATCT | |  | cDNA-amplification |
| NEB index1 | CAAGCAGAAGACGGCATACGAGATCGTGATGTGACTGGAGTTCAGACGTGTGCTCTTCCGATC-s-T | | ATCACG | Sequencing library-amplification |
| NEB index10 | CAAGCAGAAGACGGCATACGAGATAAGCTAGTGACTGGAGTTCAGACGTGTGCTCTTCCGATC-s-T | | TAGCTT | Sequencing library-amplification |
| NEB index11 | CAAGCAGAAGACGGCATACGAGATGTAGCCGTGACTGGAGTTCAGACGTGTGCTCTTCCGATC-s-T | | GGCTAC | Sequencing library-amplification |
| NEB index12 | CAAGCAGAAGACGGCATACGAGATTACAAGGTGACTGGAGTTCAGACGTGTGCTCTTCCGATC-s-T | | CTTGTA | Sequencing library-amplification |
| NEB index2 | CAAGCAGAAGACGGCATACGAGATACATCGGTGACTGGAGTTCAGACGTGTGCTCTTCCGATC-s-T | | CGATGT | Sequencing library-amplification |
| NEB index3 | CAAGCAGAAGACGGCATACGAGATGCCTAAGTGACTGGAGTTCAGACGTGTGCTCTTCCGATC-s-T | | TTAGGC | Sequencing library-amplification |
| NEB index4 | CAAGCAGAAGACGGCATACGAGATTGGTCAGTGACTGGAGTTCAGACGTGTGCTCTTCCGATC-s-T | | TGACCA | Sequencing library-amplification |
| NEB index5 | CAAGCAGAAGACGGCATACGAGATCACTGTGTGACTGGAGTTCAGACGTGTGCTCTTCCGATC-s-T | | ACAGTG | Sequencing library-amplification |
| NEB index6 | CAAGCAGAAGACGGCATACGAGATATTGGCGTGACTGGAGTTCAGACGTGTGCTCTTCCGATC-s-T | | GCCAAT | Sequencing library-amplification |
| NEB index7 | CAAGCAGAAGACGGCATACGAGATGATCTGGTGACTGGAGTTCAGACGTGTGCTCTTCCGATC-s-T | | CAGATC | Sequencing library-amplification |
| NEB index8 | CAAGCAGAAGACGGCATACGAGATTCAAGTGTGACTGGAGTTCAGACGTGTGCTCTTCCGATC-s-T | | ACTTGA | Sequencing library-amplification |
| NEB index9 | CAAGCAGAAGACGGCATACGAGATCTGATCGTGACTGGAGTTCAGACGTGTGCTCTTCCGATC-s-T | | GATCAG | Sequencing library-amplification |
| Nextera S502 | AATGATACGGCGACCACCGAGATCTACAC-CTCTCTAT-TCGTCGGCAGCGTC | | CTCTCTAT | Sequencing library-amplification |
| Nextera S503 | AATGATACGGCGACCACCGAGATCTACAC-TATCCTCT-TCGTCGGCAGCGTC | | TATCCTCT | Sequencing library-amplification |
| Nextera S505 | AATGATACGGCGACCACCGAGATCTACAC-GTAAGGAG-TCGTCGGCAGCGTC | | GTAAGGAG | Sequencing library-amplification |
| Nextera S506 | AATGATACGGCGACCACCGAGATCTACAC-ACTGCATA-TCGTCGGCAGCGTC | | ACTGCATA | Sequencing library-amplification |
| Nextera S507 | AATGATACGGCGACCACCGAGATCTACAC-AAGGAGTA-TCGTCGGCAGCGTC | | AAGGAGTA | Sequencing library-amplification |
| Nextera S508 | AATGATACGGCGACCACCGAGATCTACAC-CTAAGCCT-TCGTCGGCAGCGTC | | CTAAGCCT | Sequencing library-amplification |
| solo-RT-1 | CAGACGTGTGCTCTTCCGATCTACGGAGAGTCGACTATNNNNNNNNNTTTTTTTTTTTTTTTTTTTTTTTTTTTTTTVN | | ACGGAGAGTCGACTAT | RT-Barcode-Primer |

**Table S8 Primers for scRNA-seq library construction (continued)**

| **Primer_ID** | **Sequence 5’ to 3’** | **Barcode/index** | **Note** |
| --- | --- | --- | --- |
| solo-RT-10 | CAGACGTGTGCTCTTCCGATCTCGAGAAGTCAGTACGTNNNNNNNNNTTTTTTTTTTTTTTTTTTTTTTTTTTTTTTVN | CGAGAAGTCAGTACGT | RT-Barcode-Primer |
| solo-RT-11 | CAGACGTGTGCTCTTCCGATCTGAATGAATCTGCGACGNNNNNNNNNTTTTTTTTTTTTTTTTTTTTTTTTTTTTTTVN | GAATGAATCTGCGACG | RT-Barcode-Primer |
| solo-RT-12 | CAGACGTGTGCTCTTCCGATCTCGCTTCATCTCTGAGANNNNNNNNNTTTTTTTTTTTTTTTTTTTTTTTTTTTTTTVN | CGCTTCATCTCTGAGA | RT-Barcode-Primer |
| solo-RT-13 | CAGACGTGTGCTCTTCCGATCTTCTCTAAGTCGCATCGNNNNNNNNNTTTTTTTTTTTTTTTTTTTTTTTTTTTTTTVN | TCTCTAAGTCGCATCG | RT-Barcode-Primer |
| solo-RT-14 | CAGACGTGTGCTCTTCCGATCTGATGCTAAGGCGATACNNNNNNNNNTTTTTTTTTTTTTTTTTTTTTTTTTTTTTTVN | GATGCTAAGGCGATAC | RT-Barcode-Primer |
| solo-RT-15 | CAGACGTGTGCTCTTCCGATCTTCGAGGCAGACTAAGTNNNNNNNNNTTTTTTTTTTTTTTTTTTTTTTTTTTTTTTVN | TCGAGGCAGACTAAGT | RT-Barcode-Primer |
| solo-RT-16 | CAGACGTGTGCTCTTCCGATCTACTGAGTAGGCTAGCANNNNNNNNNTTTTTTTTTTTTTTTTTTTTTTTTTTTTTTVN | ACTGAGTAGGCTAGCA | RT-Barcode-Primer |
| solo-RT-17 | CAGACGTGTGCTCTTCCGATCTTACGGATCAGCATGAGNNNNNNNNNTTTTTTTTTTTTTTTTTTTTTTTTTTTTTTVN | TACGGATCAGCATGAG | RT-Barcode-Primer |
| solo-RT-18 | CAGACGTGTGCTCTTCCGATCTACTGAGTAGCGATGACNNNNNNNNNTTTTTTTTTTTTTTTTTTTTTTTTTTTTTTVN | ACTGAGTAGCGATGAC | RT-Barcode-Primer |
| solo-RT-19 | CAGACGTGTGCTCTTCCGATCTCTAGCCTCAGTATGCTNNNNNNNNNTTTTTTTTTTTTTTTTTTTTTTTTTTTTTTVN | CTAGCCTCAGTATGCT | RT-Barcode-Primer |
| solo-RT-2 | CAGACGTGTGCTCTTCCGATCTATCGAGTAGTCCTCCTNNNNNNNNNTTTTTTTTTTTTTTTTTTTTTTTTTTTTTTVN | ATCGAGTAGTCCTCCT | RT-Barcode-Primer |
| solo-RT-20 | CAGACGTGTGCTCTTCCGATCTATCGAGTAGACGACGTNNNNNNNNNTTTTTTTTTTTTTTTTTTTTTTTTTTTTTTVN | ATCGAGTAGACGACGT | RT-Barcode-Primer |
| solo-RT-21 | CAGACGTGTGCTCTTCCGATCTGATCTAGTCAGCTCTCNNNNNNNNNTTTTTTTTTTTTTTTTTTTTTTTTTTTTTTVN | GATCTAGTCAGCTCTC | RT-Barcode-Primer |
| solo-RT-22 | CAGACGTGTGCTCTTCCGATCTGTAGTCATCATGCTCCNNNNNNNNNTTTTTTTTTTTTTTTTTTTTTTTTTTTTTTVN | GTAGTCATCATGCTCC | RT-Barcode-Primer |
| solo-RT-23 | CAGACGTGTGCTCTTCCGATCTAGTAGTCAGGATTCGGNNNNNNNNNTTTTTTTTTTTTTTTTTTTTTTTTTTTTTTVN | AGTAGTCAGGATTCGG | RT-Barcode-Primer |
| solo-RT-24 | CAGACGTGTGCTCTTCCGATCTAGGCCGTAGAGACTATNNNNNNNNNTTTTTTTTTTTTTTTTTTTTTTTTTTTTTTVN | AGGCCGTAGAGACTAT | RT-Barcode-Primer |
| solo-RT-25 | CAGACGTGTGCTCTTCCGATCTTCAGCTCGTCGCATATNNNNNNNNNTTTTTTTTTTTTTTTTTTTTTTTTTTTTTTVN | TCAGCTCGTCGCATAT | RT-Barcode-Primer |
| solo-RT-26 | CAGACGTGTGCTCTTCCGATCTCAGAATCAGCGACGTANNNNNNNNNTTTTTTTTTTTTTTTTTTTTTTTTTTTTTTVN | CAGAATCAGCGACGTA | RT-Barcode-Primer |
| solo-RT-27 | CAGACGTGTGCTCTTCCGATCTGAGCAGAGTATCAGTCNNNNNNNNNTTTTTTTTTTTTTTTTTTTTTTTTTTTTTTVN | GAGCAGAGTATCAGTC | RT-Barcode-Primer |
| solo-RT-28 | CAGACGTGTGCTCTTCCGATCTGCTCCTATCTGCGTAANNNNNNNNNTTTTTTTTTTTTTTTTTTTTTTTTTTTTTTVN | GCTCCTATCTGCGTAA | RT-Barcode-Primer |
| solo-RT-29 | CAGACGTGTGCTCTTCCGATCTTACTCGCGTAGAGGAANNNNNNNNNTTTTTTTTTTTTTTTTTTTTTTTTTTTTTTVN | TACTCGCGTAGAGGAA | RT-Barcode-Primer |

**Table S8 Primers for scRNA-seq library construction (continued)**

| **Primer_ID** | **Sequence 5’ to 3’** | **Barcode/index** | **Note** |
| --- | --- | --- | --- |
| solo-RT-3 | CAGACGTGTGCTCTTCCGATCTCTGATAGCAGTCTTCCNNNNNNNNNTTTTTTTTTTTTTTTTTTTTTTTTTTTTTTVN | CTGATAGCAGTCTTCC | RT-Barcode-Primer |
| solo-RT-30 | CAGACGTGTGCTCTTCCGATCTGGAATAAGTCCGTCAGNNNNNNNNNTTTTTTTTTTTTTTTTTTTTTTTTTTTTTTVN | GGAATAAGTCCGTCAG | RT-Barcode-Primer |
| solo-RT-31 | CAGACGTGTGCTCTTCCGATCTCGTAGCGAGTACGATANNNNNNNNNTTTTTTTTTTTTTTTTTTTTTTTTTTTTTTVN | CGTAGCGAGTACGATA | RT-Barcode-Primer |
| solo-RT-32 | CAGACGTGTGCTCTTCCGATCTCTGCCTAGTAGAAGGANNNNNNNNNTTTTTTTTTTTTTTTTTTTTTTTTTTTTTTVN | CTGCCTAGTAGAAGGA | RT-Barcode-Primer |
| solo-RT-33 | CAGACGTGTGCTCTTCCGATCTAAGCCGCGTATTCTCTNNNNNNNNNTTTTTTTTTTTTTTTTTTTTTTTTTTTTTTVN | AAGCCGCGTATTCTCT | RT-Barcode-Primer |
| solo-RT-34 | CAGACGTGTGCTCTTCCGATCTCAGCCGAAGTACGATANNNNNNNNNTTTTTTTTTTTTTTTTTTTTTTTTTTTTTTVN | CAGCCGAAGTACGATA | RT-Barcode-Primer |
| solo-RT-35 | CAGACGTGTGCTCTTCCGATCTAGCTCTCTCTAGAGTCNNNNNNNNNTTTTTTTTTTTTTTTTTTTTTTTTTTTTTTVN | AGCTCTCTCTAGAGTC | RT-Barcode-Primer |
| solo-RT-36 | CAGACGTGTGCTCTTCCGATCTTCAGGATCAGATCGGANNNNNNNNNTTTTTTTTTTTTTTTTTTTTTTTTTTTTTTVN | TCAGGATCAGATCGGA | RT-Barcode-Primer |
| solo-RT-37 | CAGACGTGTGCTCTTCCGATCTATGCGATTCCGCATCTNNNNNNNNNTTTTTTTTTTTTTTTTTTTTTTTTTTTTTTVN | ATGCGATTCCGCATCT | RT-Barcode-Primer |
| solo-RT-38 | CAGACGTGTGCTCTTCCGATCTGACTGCGAGAGACTTANNNNNNNNNTTTTTTTTTTTTTTTTTTTTTTTTTTTTTTVN | GACTGCGAGAGACTTA | RT-Barcode-Primer |
| solo-RT-39 | CAGACGTGTGCTCTTCCGATCTTAAGCGTCAGGATCGANNNNNNNNNTTTTTTTTTTTTTTTTTTTTTTTTTTTTTTVN | TAAGCGTCAGGATCGA | RT-Barcode-Primer |
| solo-RT-4 | CAGACGTGTGCTCTTCCGATCTCGGACTGAGCGTAATANNNNNNNNNTTTTTTTTTTTTTTTTTTTTTTTTTTTTTTVN | CGGACTGAGCGTAATA | RT-Barcode-Primer |
| solo-RT-40 | CAGACGTGTGCTCTTCCGATCTCAGTCCTAGTCATGCTNNNNNNNNNTTTTTTTTTTTTTTTTTTTTTTTTTTTTTTVN | CAGTCCTAGTCATGCT | RT-Barcode-Primer |
| solo-RT-41 | CAGACGTGTGCTCTTCCGATCTCGGAGTCGTAATAGCANNNNNNNNNTTTTTTTTTTTTTTTTTTTTTTTTTTTTTTVN | CGGAGTCGTAATAGCA | RT-Barcode-Primer |
| solo-RT-42 | CAGACGTGTGCTCTTCCGATCTCGCTATCGTATCTGCANNNNNNNNNTTTTTTTTTTTTTTTTTTTTTTTTTTTTTTVN | CGCTATCGTATCTGCA | RT-Barcode-Primer |
| solo-RT-43 | CAGACGTGTGCTCTTCCGATCTCAGCAGCTCAGATAAGNNNNNNNNNTTTTTTTTTTTTTTTTTTTTTTTTTTTTTTVN | CAGCAGCTCAGATAAG | RT-Barcode-Primer |
| solo-RT-44 | CAGACGTGTGCTCTTCCGATCTTAGCCGGCAGTATAAGNNNNNNNNNTTTTTTTTTTTTTTTTTTTTTTTTTTTTTTVN | TAGCCGGCAGTATAAG | RT-Barcode-Primer |
| solo-RT-45 | CAGACGTGTGCTCTTCCGATCTAAGGAGCAGGCTATCTNNNNNNNNNTTTTTTTTTTTTTTTTTTTTTTTTTTTTTTVN | AAGGAGCAGGCTATCT | RT-Barcode-Primer |
| solo-RT-46 | CAGACGTGTGCTCTTCCGATCTCTCCTAGTCTTCCTTCNNNNNNNNNTTTTTTTTTTTTTTTTTTTTTTTTTTTTTTVN | CTCCTAGTCTTCCTTC | RT-Barcode-Primer |
| solo-RT-47 | CAGACGTGTGCTCTTCCGATCTTCAGCTCTCGTATCAGNNNNNNNNNTTTTTTTTTTTTTTTTTTTTTTTTTTTTTTVN | TCAGCTCTCGTATCAG | RT-Barcode-Primer |
| solo-RT-48 | CAGACGTGTGCTCTTCCGATCTGAATAAGAGCAGCCTCNNNNNNNNNTTTTTTTTTTTTTTTTTTTTTTTTTTTTTTVN | GAATAAGAGCAGCCTC | RT-Barcode-Primer |

**Table S8 Primers for scRNA-seq library construction (continued)**

| **Primer_ID** | **Sequence 5’ to 3’** | **Barcode/index** | **Note** |
| --- | --- | --- | --- |
| solo-RT-49 | CAGACGTGTGCTCTTCCGATCTCTCTAATTCGGCGCATNNNNNNNNNTTTTTTTTTTTTTTTTTTTTTTTTTTTTTTVN | CTCTAATTCGGCGCAT | RT-Barcode-Primer |
| solo-RT-5 | CAGACGTGTGCTCTTCCGATCTCTCGTACTCTGCAGTANNNNNNNNNTTTTTTTTTTTTTTTTTTTTTTTTTTTTTTVN | CTCGTACTCTGCAGTA | RT-Barcode-Primer |
| solo-RT-50 | CAGACGTGTGCTCTTCCGATCTTCAGCTCTCGGAATCTNNNNNNNNNTTTTTTTTTTTTTTTTTTTTTTTTTTTTTTVN | TCAGCTCTCGGAATCT | RT-Barcode-Primer |
| solo-RT-51 | CAGACGTGTGCTCTTCCGATCTCTGATCCGTAGCCTATNNNNNNNNNTTTTTTTTTTTTTTTTTTTTTTTTTTTTTTVN | CTGATCCGTAGCCTAT | RT-Barcode-Primer |
| solo-RT-52 | CAGACGTGTGCTCTTCCGATCTAGAATAGCAGACGCTCNNNNNNNNNTTTTTTTTTTTTTTTTTTTTTTTTTTTTTTVN | AGAATAGCAGACGCTC | RT-Barcode-Primer |
| solo-RT-53 | CAGACGTGTGCTCTTCCGATCTAGCGTATTCGACGGAANNNNNNNNNTTTTTTTTTTTTTTTTTTTTTTTTTTTTTTVN | AGCGTATTCGACGGAA | RT-Barcode-Primer |
| solo-RT-54 | CAGACGTGTGCTCTTCCGATCTACGCAGCAGACTAGATNNNNNNNNNTTTTTTTTTTTTTTTTTTTTTTTTTTTTTTVN | ACGCAGCAGACTAGAT | RT-Barcode-Primer |
| solo-RT-55 | CAGACGTGTGCTCTTCCGATCTGACGGCTAGAGTAATCNNNNNNNNNTTTTTTTTTTTTTTTTTTTTTTTTTTTTTTVN | GACGGCTAGAGTAATC | RT-Barcode-Primer |
| solo-RT-56 | CAGACGTGTGCTCTTCCGATCTACTGATGAGGATTCGGNNNNNNNNNTTTTTTTTTTTTTTTTTTTTTTTTTTTTTTVN | ACTGATGAGGATTCGG | RT-Barcode-Primer |
| solo-RT-57 | CAGACGTGTGCTCTTCCGATCTAGAGCTTCAGTCCTTCNNNNNNNNNTTTTTTTTTTTTTTTTTTTTTTTTTTTTTTVN | AGAGCTTCAGTCCTTC | RT-Barcode-Primer |
| solo-RT-58 | CAGACGTGTGCTCTTCCGATCTAAGTCTGCATCTCGCTNNNNNNNNNTTTTTTTTTTTTTTTTTTTTTTTTTTTTTTVN | AAGTCTGCATCTCGCT | RT-Barcode-Primer |
| solo-RT-59 | CAGACGTGTGCTCTTCCGATCTCTAGAGTAGGCTAGCANNNNNNNNNTTTTTTTTTTTTTTTTTTTTTTTTTTTTTTVN | CTAGAGTAGGCTAGCA | RT-Barcode-Primer |
| solo-RT-6 | CAGACGTGTGCTCTTCCGATCTAGCTCTCGTCATATCGNNNNNNNNNTTTTTTTTTTTTTTTTTTTTTTTTTTTTTTVN | AGCTCTCGTCATATCG | RT-Barcode-Primer |
| solo-RT-60 | CAGACGTGTGCTCTTCCGATCTTAAGCGTAGCAGGCTANNNNNNNNNTTTTTTTTTTTTTTTTTTTTTTTTTTTTTTVN | TAAGCGTAGCAGGCTA | RT-Barcode-Primer |
| solo-RT-61 | CAGACGTGTGCTCTTCCGATCTAGAATAGTCGCGTAGCNNNNNNNNNTTTTTTTTTTTTTTTTTTTTTTTTTTTTTTVN | AGAATAGTCGCGTAGC | RT-Barcode-Primer |
| solo-RT-62 | CAGACGTGTGCTCTTCCGATCTAAGTCTGAGACTCGGANNNNNNNNNTTTTTTTTTTTTTTTTTTTTTTTTTTTTTTVN | AAGTCTGAGACTCGGA | RT-Barcode-Primer |
| solo-RT-63 | CAGACGTGTGCTCTTCCGATCTTCAGATGAGGCTCAGANNNNNNNNNTTTTTTTTTTTTTTTTTTTTTTTTTTTTTTVN | TCAGATGAGGCTCAGA | RT-Barcode-Primer |
| solo-RT-64 | CAGACGTGTGCTCTTCCGATCTGACGGCTCATAGGATANNNNNNNNNTTTTTTTTTTTTTTTTTTTTTTTTTTTTTTVN | GACGGCTCATAGGATA | RT-Barcode-Primer |
| solo-RT-65 | CAGACGTGTGCTCTTCCGATCTAGCTCTCTCTTCGAGANNNNNNNNNTTTTTTTTTTTTTTTTTTTTTTTTTTTTTTVN | AGCTCTCTCTTCGAGA | RT-Barcode-Primer |
| solo-RT-66 | CAGACGTGTGCTCTTCCGATCTCTGAAGTCAGGAATCGNNNNNNNNNTTTTTTTTTTTTTTTTTTTTTTTTTTTTTTVN | CTGAAGTCAGGAATCG | RT-Barcode-Primer |
| solo-RT-67 | CAGACGTGTGCTCTTCCGATCTTGACTAGTCTCTGCTGNNNNNNNNNTTTTTTTTTTTTTTTTTTTTTTTTTTTTTTVN | TGACTAGTCTCTGCTG | RT-Barcode-Primer |

**Table S8 Primers for scRNA-seq library construction (continued)**

| **Primer_ID** | **Sequence 5’ to 3’** | **Barcode/index** | **Note** |
| --- | --- | --- | --- |
| solo-RT-68 | CAGACGTGTGCTCTTCCGATCTTATCAGGCATGCCTTCNNNNNNNNNTTTTTTTTTTTTTTTTTTTTTTTTTTTTTTVN | TATCAGGCATGCCTTC | RT-Barcode-Primer |
| solo-RT-69 | CAGACGTGTGCTCTTCCGATCTGATCGATCAGGCAGTANNNNNNNNNTTTTTTTTTTTTTTTTTTTTTTTTTTTTTTVN | GATCGATCAGGCAGTA | RT-Barcode-Primer |
| solo-RT-7 | CAGACGTGTGCTCTTCCGATCTTCTTCGGAGGCTCATTNNNNNNNNNTTTTTTTTTTTTTTTTTTTTTTTTTTTTTTVN | TCTTCGGAGGCTCATT | RT-Barcode-Primer |
| solo-RT-70 | CAGACGTGTGCTCTTCCGATCTGATCGCGCATTATCTCNNNNNNNNNTTTTTTTTTTTTTTTTTTTTTTTTTTTTTTVN | GATCGCGCATTATCTC | RT-Barcode-Primer |
| solo-RT-71 | CAGACGTGTGCTCTTCCGATCTGTATCTTCAGCTCCGANNNNNNNNNTTTTTTTTTTTTTTTTTTTTTTTTTTTTTTVN | GTATCTTCAGCTCCGA | RT-Barcode-Primer |
| solo-RT-72 | CAGACGTGTGCTCTTCCGATCTCTTCTCTTCCTCCTAGNNNNNNNNNTTTTTTTTTTTTTTTTTTTTTTTTTTTTTTVN | CTTCTCTTCCTCCTAG | RT-Barcode-Primer |
| solo-RT-73 | CAGACGTGTGCTCTTCCGATCTAGAGCTTAGAGCCTAGNNNNNNNNNTTTTTTTTTTTTTTTTTTTTTTTTTTTTTTVN | AGAGCTTAGAGCCTAG | RT-Barcode-Primer |
| solo-RT-74 | CAGACGTGTGCTCTTCCGATCTAGTAGTCAGACTAGGCNNNNNNNNNTTTTTTTTTTTTTTTTTTTTTTTTTTTTTTVN | AGTAGTCAGACTAGGC | RT-Barcode-Primer |
| solo-RT-75 | CAGACGTGTGCTCTTCCGATCTTCTGAGAAGGCTCAGANNNNNNNNNTTTTTTTTTTTTTTTTTTTTTTTTTTTTTTVN | TCTGAGAAGGCTCAGA | RT-Barcode-Primer |
| solo-RT-76 | CAGACGTGTGCTCTTCCGATCTTATCAGGCAGGCTGAANNNNNNNNNTTTTTTTTTTTTTTTTTTTTTTTTTTTTTTVN | TATCAGGCAGGCTGAA | RT-Barcode-Primer |
| solo-RT-77 | CAGACGTGTGCTCTTCCGATCTGTAGTCATCGATGAGGNNNNNNNNNTTTTTTTTTTTTTTTTTTTTTTTTTTTTTTVN | GTAGTCATCGATGAGG | RT-Barcode-Primer |
| solo-RT-78 | CAGACGTGTGCTCTTCCGATCTAAGTCTGAGACTAGGCNNNNNNNNNTTTTTTTTTTTTTTTTTTTTTTTTTTTTTTVN | AAGTCTGAGACTAGGC | RT-Barcode-Primer |
| solo-RT-79 | CAGACGTGTGCTCTTCCGATCTTCGCGAGTCCTATTCANNNNNNNNNTTTTTTTTTTTTTTTTTTTTTTTTTTTTTTVN | TCGCGAGTCCTATTCA | RT-Barcode-Primer |
| solo-RT-8 | CAGACGTGTGCTCTTCCGATCTGGAATAAGTCTAGCGCNNNNNNNNNTTTTTTTTTTTTTTTTTTTTTTTTTTTTTTVN | GGAATAAGTCTAGCGC | RT-Barcode-Primer |
| solo-RT-80 | CAGACGTGTGCTCTTCCGATCTCGTAGGCAGCTGATAANNNNNNNNNTTTTTTTTTTTTTTTTTTTTTTTTTTTTTTVN | CGTAGGCAGCTGATAA | RT-Barcode-Primer |
| solo-RT-81 | CAGACGTGTGCTCTTCCGATCTTACGGATTCAGAGACGNNNNNNNNNTTTTTTTTTTTTTTTTTTTTTTTTTTTTTTVN | TACGGATTCAGAGACG | RT-Barcode-Primer |
| solo-RT-82 | CAGACGTGTGCTCTTCCGATCTAGTAGTCGTCTCTCTGNNNNNNNNNTTTTTTTTTTTTTTTTTTTTTTTTTTTTTTVN | AGTAGTCGTCTCTCTG | RT-Barcode-Primer |
| solo-RT-83 | CAGACGTGTGCTCTTCCGATCTGCTCCTATCATATCGGNNNNNNNNNTTTTTTTTTTTTTTTTTTTTTTTTTTTTTTVN | GCTCCTATCATATCGG | RT-Barcode-Primer |
| solo-RT-84 | CAGACGTGTGCTCTTCCGATCTAGCTCCTTCCGTAGTANNNNNNNNNTTTTTTTTTTTTTTTTTTTTTTTTTTTTTTVN | AGCTCCTTCCGTAGTA | RT-Barcode-Primer |
| solo-RT-85 | CAGACGTGTGCTCTTCCGATCTGTCGTAAGTCGTCTTCNNNNNNNNNTTTTTTTTTTTTTTTTTTTTTTTTTTTTTTVN | GTCGTAAGTCGTCTTC | RT-Barcode-Primer |
| solo-RT-86 | CAGACGTGTGCTCTTCCGATCTCAGAATCAGTCCTCCTNNNNNNNNNTTTTTTTTTTTTTTTTTTTTTTTTTTTTTTVN | CAGAATCAGTCCTCCT | RT-Barcode-Primer |

**Table S8 Primers for scRNA-seq library construction (continued)**

| **Primer_ID** | **Sequence 5’ to 3’** | **Barcode/index** | **Note** |
| --- | --- | --- | --- |
| solo-RT-87 | CAGACGTGTGCTCTTCCGATCTAGCAGCCTCAGATAAGNNNNNNNNNTTTTTTTTTTTTTTTTTTTTTTTTTTTTTTVN | AGCAGCCTCAGATAAG | RT-Barcode-Primer |
| solo-RT-88 | CAGACGTGTGCTCTTCCGATCTTGAGCATAGCTAGGCANNNNNNNNNTTTTTTTTTTTTTTTTTTTTTTTTTTTTTTVN | TGAGCATAGCTAGGCA | RT-Barcode-Primer |
| solo-RT-89 | CAGACGTGTGCTCTTCCGATCTGATCTAGAGGCTCAGANNNNNNNNNTTTTTTTTTTTTTTTTTTTTTTTTTTTTTTVN | GATCTAGAGGCTCAGA | RT-Barcode-Primer |
| solo-RT-9 | CAGACGTGTGCTCTTCCGATCTTAAGAGATCGCGTAGCNNNNNNNNNTTTTTTTTTTTTTTTTTTTTTTTTTTTTTTVN | TAAGAGATCGCGTAGC | RT-Barcode-Primer |
| solo-RT-90 | CAGACGTGTGCTCTTCCGATCTCTCATTATCGGCTACGNNNNNNNNNTTTTTTTTTTTTTTTTTTTTTTTTTTTTTTVN | CTCATTATCGGCTACG | RT-Barcode-Primer |
| solo-RT-91 | CAGACGTGTGCTCTTCCGATCTAGCATACGTCAGAAGCNNNNNNNNNTTTTTTTTTTTTTTTTTTTTTTTTTTTTTTVN | AGCATACGTCAGAAGC | RT-Barcode-Primer |
| solo-RT-92 | CAGACGTGTGCTCTTCCGATCTCTCGAGGAGCGTAATANNNNNNNNNTTTTTTTTTTTTTTTTTTTTTTTTTTTTTTVN | CTCGAGGAGCGTAATA | RT-Barcode-Primer |
| solo-RT-93 | CAGACGTGTGCTCTTCCGATCTCGAGAAGAGAGTAAGGNNNNNNNNNTTTTTTTTTTTTTTTTTTTTTTTTTTTTTTVN | CGAGAAGAGAGTAAGG | RT-Barcode-Primer |
| solo-RT-94 | CAGACGTGTGCTCTTCCGATCTTCGTAGATCATGCTCCNNNNNNNNNTTTTTTTTTTTTTTTTTTTTTTTTTTTTTTVN | TCGTAGATCATGCTCC | RT-Barcode-Primer |
| solo-RT-95 | CAGACGTGTGCTCTTCCGATCTTTCTTAGTCGGCTACGNNNNNNNNNTTTTTTTTTTTTTTTTTTTTTTTTTTTTTTVN | TTCTTAGTCGGCTACG | RT-Barcode-Primer |
| solo-RT-96 | CAGACGTGTGCTCTTCCGATCTTAGGCATTCTGCTGCTNNNNNNNNNTTTTTTTTTTTTTTTTTTTTTTTTTTTTTTVN | TAGGCATTCTGCTGCT | RT-Barcode-Primer |
