## Supplemental table 1 to 9 for "Mouse Single Islet β Cell Transcriptomics Reveal Sexually Dimorphic Transcriptomes and Type 2 Diabetes Genes": Table S9.docx

**Table S9 Differential expression genes detected by edgeR**

| **Type** | **Gene symbol** | **log2FoldChange** | **P value** | **P adjust** |
| --- | --- | --- | --- | --- |
| Female specific upregulated T2D genes | *Fxyd2* | 3.354357889 | 3.82E-26 | 3.41E-22 |
|  | *Hspa8* | 0.382179406 | 1.46E-14 | 5.48E-11 |
|  | *G6pc2* | 0.290553436 | 2.31E-10 | 3.44E-07 |
|  | *Rnase4* | 0.431257055 | 4.15E-10 | 5.29E-07 |
|  | *Ttr* | 0.348460395 | 5.53E-09 | 4.95E-06 |
|  | *Rgs2* | 0.728408297 | 7.74E-09 | 6.29E-06 |
|  | *Gc* | 1.232621273 | 9.25E-09 | 6.89E-06 |
|  | *Hsp90aa1* | 0.456117826 | 1.53E-08 | 1.05E-05 |
|  | *Cpe* | 0.30188975 | 2.16E-08 | 1.38E-05 |
|  | *Scg3* | 0.255857965 | 3.40E-08 | 2.02E-05 |
|  | *Serp1* | 0.345391136 | 5.04E-08 | 2.81E-05 |
|  | *Cd9* | 1.521235667 | 6.57E-08 | 3.46E-05 |
|  | *Igfbp7* | 1.237888119 | 7.96E-08 | 3.95E-05 |
|  | *Hsph1* | 0.8571005 | 1.54E-07 | 7.26E-05 |
|  | *Dapl1* | 1.049490845 | 4.32E-07 | 0.000175712 |
|  | *Bhlha15* | 0.927557189 | 4.77E-07 | 0.000185284 |
|  | *Ccnd1* | 0.848356909 | 6.08E-07 | 0.000226319 |
|  | *Pde1c* | 1.12080518 | 6.50E-07 | 0.000232329 |
|  | *Etv1* | 1.279438614 | 2.20E-06 | 0.000727551 |
|  | *Pcbp1* | 0.57313111 | 3.12E-06 | 0.000997484 |
|  | *Cd44* | 1.093472746 | 3.54E-06 | 0.001090774 |
|  | *Mgat2* | 0.705648964 | 3.77E-06 | 0.001123694 |
|  | *Pcsk1n* | 0.216300865 | 4.10E-06 | 0.001155841 |
|  | *Cflar* | 0.831824993 | 1.20E-05 | 0.002888416 |
|  | *Necab2* | 0.756155104 | 1.28E-05 | 0.003000325 |
|  | *Cmas* | 0.951884032 | 2.25E-05 | 0.004569106 |
|  | *Stip1* | 0.585582546 | 2.46E-05 | 0.004777938 |
|  | *Scel* | 0.973837391 | 2.57E-05 | 0.004884372 |
|  | *Lamp1* | 0.494011143 | 2.68E-05 | 0.004997391 |
|  | *Ero1lb* | 0.235928259 | 2.80E-05 | 0.005100734 |
|  | *Rap1gap2* | 0.502172489 | 3.21E-05 | 0.005615152 |
|  | *H2afy* | 0.555254199 | 3.83E-05 | 0.00622341 |
|  | *Gca* | 0.918439941 | 7.42E-05 | 0.01086258 |
|  | *Ybx1* | 0.897053495 | 7.53E-05 | 0.01086258 |
|  | *Fxyd6* | 0.547792736 | 9.19E-05 | 0.012641258 |
|  | *Spc25* | 0.326788672 | 0.000102642 | 0.013359347 |
|  | *Osgin1* | 0.870209579 | 0.000105445 | 0.013465283 |
|  | *Vcp* | 0.293355234 | 0.000109357 | 0.013645467 |
|  | *Edem2* | 0.278749028 | 0.000109909 | 0.013645467 |
|  | *Chchd10* | 0.39019538 | 0.000120401 | 0.014196074 |
|  | *Hspa13* | 0.469986476 | 0.000121751 | 0.014196074 |
|  | *Sec23b* | 0.299131471 | 0.000122284 | 0.014196074 |
|  | *Rela* | 0.868492257 | 0.000131244 | 0.014747189 |
|  | *Cpt1a* | 0.84471271 | 0.000160088 | 0.017451506 |
|  | *Jam2* | 1.002172611 | 0.000163326 | 0.01758997 |
|  | *Srsf3* | 0.495628645 | 0.000166006 | 0.017665821 |

**Table S9 Differential expression genes detected by edgeR (continued)**

| **Type** | **Gene symbol** | **log2FoldChange** | **P value** | **P adjust** |
| --- | --- | --- | --- | --- |
| Female specific upregulated T2D genes | *Prrc2b* | 0.484321886 | 0.000175915 | 0.018500026 |
|  | *Gng4* | 0.462097987 | 0.000198293 | 0.020142495 |
|  | *Actb* | 0.213556443 | 0.000207346 | 0.02082547 |
|  | *Mrps12* | 0.395460713 | 0.000228137 | 0.022416372 |
|  | *Rspry1* | 0.862110532 | 0.000228201 | 0.022416372 |
|  | *Pcp4* | 1.014793426 | 0.000239533 | 0.023273775 |
|  | *Oxtr* | 0.979634136 | 0.000282434 | 0.027147075 |
|  | *Sel1l3* | 0.840687127 | 0.000309702 | 0.029451324 |
|  | *Bfar* | 0.716278364 | 0.000327839 | 0.030067685 |
|  | *Rab3a* | 0.342816378 | 0.000329741 | 0.030067685 |
|  | *Tmem160* | 0.357925987 | 0.000335462 | 0.030067685 |
|  | *Cdk11b* | 0.473589059 | 0.000336365 | 0.030067685 |
|  | *Gde1* | 0.316374067 | 0.000371981 | 0.032282933 |
|  | *Top1* | 0.380497303 | 0.000392128 | 0.033055718 |
|  | *Thap1* | 0.875684545 | 0.00043066 | 0.035063603 |
|  | *Iffo1* | 0.593319762 | 0.000451168 | 0.036008812 |
|  | *Aplp1* | 0.275493888 | 0.000527403 | 0.040641858 |
|  | *Manf* | 0.235332899 | 0.000555826 | 0.042466036 |
|  | *Pnrc1* | 0.684799129 | 0.000620018 | 0.046574272 |
|  | *Fh1* | 0.481556376 | 0.000676217 | 0.049339806 |
|  | *Tmsb10* | 0.424013085 | 0.000697845 | 0.049904293 |
| Female specific downregulated T2D genes | *Matn2* | -1.550122309 | 1.84E-14 | 5.48E-11 |
|  | *mt-Cytb* | -0.326972603 | 1.54E-12 | 3.45E-09 |
|  | *Fmo1* | -1.466941836 | 5.11E-11 | 9.14E-08 |
|  | *Sdf2l1* | -0.651903764 | 5.42E-10 | 6.06E-07 |
|  | *2900055J20Rik* | -0.76908299 | 4.34E-09 | 4.31E-06 |
|  | *Shisal2b* | -1.116500876 | 2.52E-07 | 0.000112429 |
|  | *Cox6a2* | -0.586614346 | 3.29E-07 | 0.000139912 |
|  | *Rps19* | -0.320075598 | 7.99E-07 | 0.000274835 |
|  | *Nipal3* | -0.35691335 | 4.14E-06 | 0.001155841 |
|  | *Gnai2* | -0.247188283 | 4.83E-06 | 0.001309024 |
|  | *Igfbp5* | -1.26599221 | 8.75E-06 | 0.002299284 |
|  | *Creg1* | -0.355245 | 1.07E-05 | 0.002701584 |
|  | *Pdcd10* | -0.760877425 | 1.09E-05 | 0.002701584 |
|  | *Ssr4* | -0.270348681 | 1.36E-05 | 0.003110412 |
|  | *Mt1* | -0.397758633 | 1.58E-05 | 0.003443001 |
|  | *Cox6a1* | -0.304590828 | 1.90E-05 | 0.004050366 |
|  | *Mt2* | -0.554658612 | 1.97E-05 | 0.004090538 |
|  | *Dlgap1* | -0.95239107 | 2.40E-05 | 0.004772306 |
|  | *Pdia3* | -0.22437149 | 3.04E-05 | 0.005440471 |
|  | *Arpc5* | -0.355595018 | 3.27E-05 | 0.005615152 |
|  | *Ndufs2* | -0.395075398 | 3.59E-05 | 0.005936981 |
|  | *Stxbp5l* | -1.202333365 | 4.07E-05 | 0.00649581 |
|  | *Fkbp9* | -0.411156285 | 5.13E-05 | 0.008041106 |
|  | *Chac1* | -0.928112648 | 5.92E-05 | 0.009130811 |

**Table S9 Differential expression genes detected by edgeR (continued)**

| **Type** | **Gene symbol** | **log2FoldChange** | **P value** | **P adjust** |
| --- | --- | --- | --- | --- |
| Female specific downregulated T2D genes | *Sec14l4* | -0.564240865 | 6.61E-05 | 0.009842642 |
|  | *Uqcrb* | -0.318830115 | 7.74E-05 | 0.0109842 |
|  | *Atf5* | -0.475228845 | 9.05E-05 | 0.012641234 |
|  | *Sec11c* | -0.258233146 | 9.75E-05 | 0.013205309 |
|  | *Rpl37a* | -0.221216239 | 0.000103121 | 0.013359347 |
|  | *Mapk15* | -0.889728062 | 0.000122183 | 0.014196074 |
|  | *Scn9a* | -0.780728859 | 0.000127674 | 0.014631742 |
|  | *Hdac5* | -0.679227972 | 0.000145756 | 0.016085358 |
|  | *Tma7* | -0.281121442 | 0.000184917 | 0.019220637 |
|  | *Rogdi* | -0.507099189 | 0.000192213 | 0.019749383 |
|  | *Thsd4* | -1.027724399 | 0.000325292 | 0.030067685 |
|  | *Ddost* | -0.200958966 | 0.000358821 | 0.031538138 |
|  | *Ndufb8* | -0.275464093 | 0.000359871 | 0.031538138 |
|  | *4732471J01Rik* | -0.700298209 | 0.000389051 | 0.033055718 |
|  | *Jakmip1* | -0.569525829 | 0.00039284 | 0.033055718 |
|  | *Ncoa1* | -0.370809892 | 0.000395678 | 0.033055718 |
|  | *Trim24* | -1.057898723 | 0.00043148 | 0.035063603 |
|  | *Mthfd1* | -0.401899739 | 0.000449149 | 0.036008812 |
|  | *Jup* | -0.376714547 | 0.000498755 | 0.039401076 |
|  | *Tmem181b-ps* | -0.344080591 | 0.000502486 | 0.039401076 |
|  | *Pigp* | -0.453809893 | 0.000515002 | 0.040031338 |
|  | *Swi5* | -0.249465763 | 0.000566146 | 0.042887921 |
|  | *Slc1a4* | -1.004152602 | 0.00065337 | 0.048670621 |
|  | *Plut* | -0.675133377 | 0.000671489 | 0.049339806 |
|  | *Ccdc12* | -0.556428323 | 0.000678912 | 0.049339806 |
|  | *Ewsr1* | -0.365229102 | 0.000697198 | 0.049904293 |
| Male specific upregulated T2D genes | *Iapp* | 0.365696475 | 2.67E-44 | 2.45E-40 |
|  | *Pabpc1* | 0.952773421 | 5.18E-28 | 2.38E-24 |
|  | *Rgs2* | 0.925627497 | 2.68E-13 | 2.73E-10 |
|  | *Degs1* | 0.450824168 | 2.81E-11 | 2.58E-08 |
|  | *Creld2* | 0.482314582 | 8.91E-09 | 4.30E-06 |
|  | *Wnt4* | 0.676054526 | 1.77E-08 | 7.08E-06 |
|  | *P4hb* | 0.276242987 | 2.11E-08 | 7.76E-06 |
|  | *Ppia* | 0.445318142 | 3.86E-08 | 1.31E-05 |
|  | *Rps29* | 0.374728484 | 4.54E-08 | 1.49E-05 |
|  | *Rpl23* | 0.338246702 | 5.50E-08 | 1.74E-05 |
|  | *Pdyn* | 1.24676481 | 2.36E-07 | 5.71E-05 |
|  | *Fh1* | 0.575419592 | 2.47E-07 | 5.81E-05 |
|  | *Gcg* | 1.049962199 | 3.19E-07 | 7.33E-05 |
|  | *Tmem39a* | 0.600211872 | 1.01E-06 | 0.00020623 |
|  | *Rps27a* | 0.286279266 | 1.82E-06 | 0.000328252 |
|  | *Dvl1* | 0.834743448 | 2.17E-06 | 0.000376264 |
|  | *Ankrd46* | 0.767788654 | 4.21E-06 | 0.000677489 |
|  | *Rps16* | 0.346369534 | 4.52E-06 | 0.000705046 |
|  | *Srm* | 0.687894708 | 6.41E-06 | 0.000965154 |

**Table S9 Differential expression genes detected by edgeR (continued)**

| **Type** | **Gene symbol** | **log2FoldChange** | **P value** | **P adjust** |
| --- | --- | --- | --- | --- |
| Male specific upregulated T2D genes | *Nf1* | 0.98983317 | 8.84E-06 | 0.001268907 |
|  | *Dapl1* | 0.628467129 | 9.64E-06 | 0.00134103 |
|  | *Tpt1* | 0.25722494 | 1.32E-05 | 0.001806937 |
|  | *Gabbr2* | 0.889057535 | 1.38E-05 | 0.001862385 |
|  | *Cdh12* | 0.825587961 | 1.60E-05 | 0.002130196 |
|  | *Ubr4* | 0.386592421 | 1.91E-05 | 0.002480142 |
|  | *Copb2* | 0.289780347 | 2.00E-05 | 0.002483024 |
|  | *Pde1c* | 0.848353759 | 2.56E-05 | 0.003013108 |
|  | *Rpsa* | 0.311415137 | 2.78E-05 | 0.003227498 |
|  | *Arglu1* | 0.708440292 | 2.95E-05 | 0.00338888 |
|  | *Ank* | 0.614532645 | 3.42E-05 | 0.003784912 |
|  | *Tmed3* | 0.210845869 | 3.64E-05 | 0.00395844 |
|  | *Zxda* | 0.974754082 | 4.64E-05 | 0.004876044 |
|  | *Rps4x* | 0.370358124 | 6.44E-05 | 0.006427388 |
|  | *Isg20* | 0.348497049 | 7.62E-05 | 0.007426023 |
|  | *Slc33a1* | 0.475479732 | 7.68E-05 | 0.007426023 |
|  | *Prkcb* | 0.393200493 | 7.94E-05 | 0.007599825 |
|  | *Rps28* | 0.308170062 | 8.57E-05 | 0.008020687 |
|  | *AI597479* | 0.797233482 | 0.000111193 | 0.009723333 |
|  | *Gadd45b* | 0.918866727 | 0.000137765 | 0.011278177 |
|  | *Pde3b* | 0.573382297 | 0.000138782 | 0.011278177 |
|  | *Tmem63b* | 0.681064033 | 0.00016671 | 0.012973684 |
|  | *Dpagt1* | 0.703702225 | 0.000183533 | 0.013939865 |
|  | *Ahsa2* | 0.753680436 | 0.000192514 | 0.014417441 |
|  | *Usp35* | 0.745270337 | 0.000194959 | 0.014417441 |
|  | *Rps25* | 0.348480965 | 0.000197003 | 0.014417441 |
|  | *Atp5g3* | 0.447833052 | 0.000209852 | 0.015173763 |
|  | *Rps3* | 0.263783078 | 0.000239696 | 0.016931769 |
|  | *Rplp1* | 0.242877402 | 0.000246361 | 0.017269739 |
|  | *Rpl7a* | 0.399490277 | 0.000305764 | 0.020579067 |
|  | *Jup* | 0.371598965 | 0.000324627 | 0.021446408 |
|  | *Gnb1* | 0.453299293 | 0.000338031 | 0.022172393 |
|  | *Arel1* | 0.678981361 | 0.000347011 | 0.022374065 |
|  | *Rps3a1* | 0.244417525 | 0.000348415 | 0.022374065 |
|  | *Ube2j1* | 0.466465423 | 0.000394395 | 0.024977474 |
|  | *Hars* | 0.586670571 | 0.000414778 | 0.02591091 |
|  | *Gemin6* | 0.734957265 | 0.000479597 | 0.02955797 |
|  | *Rfx6* | 0.408587174 | 0.000515178 | 0.030645654 |
|  | *Mta1* | 0.590322244 | 0.000517268 | 0.030645654 |
|  | *Rimbp2* | 0.420366568 | 0.000521745 | 0.030712733 |
|  | *Actr3* | 0.395979792 | 0.00054289 | 0.031347343 |
|  | *Emb* | 0.210049844 | 0.00054618 | 0.031347343 |
|  | *Dnajb9* | 0.242965446 | 0.000584382 | 0.03333158 |
|  | *Rps17* | 0.300828735 | 0.000588034 | 0.03333284 |

**Table S9 Differential expression genes detected by edgeR (continued)**

| **Type** | **Gene symbol** | **log2FoldChange** | **P value** | **P adjust** |
| --- | --- | --- | --- | --- |
| Male specific upregulated T2D genes | *Ampd2* | 0.499544667 | 0.000599156 | 0.033569875 |
|  | *Cd44* | 0.661031713 | 0.000637629 | 0.035137389 |
|  | *Zbtb20* | 0.413065803 | 0.000654298 | 0.035552797 |
|  | *Gapdh* | 0.247568276 | 0.000668942 | 0.03613467 |
|  | *Gmds* | 0.473512287 | 0.000714333 | 0.03791745 |
|  | *Slc25a3* | 0.207921989 | 0.000739243 | 0.039014173 |
|  | *Ostc* | 0.248827918 | 0.000807376 | 0.041649353 |
|  | *Ndufb3* | 0.327775815 | 0.000897709 | 0.045405735 |
|  | *Rps26* | 0.32412455 | 0.000967947 | 0.047788493 |
| Male specific downregulated T2D genes | *mt-Rnr1* | -0.316918703 | 4.23E-24 | 1.29E-20 |
|  | *Slc30a8* | -0.320037583 | 4.61E-16 | 1.06E-12 |
|  | *Malat1* | -0.201242921 | 1.27E-14 | 2.32E-11 |
|  | *Cox4i1* | -0.277137859 | 5.13E-14 | 7.85E-11 |
|  | *Ubb* | -0.235718383 | 8.70E-14 | 1.14E-10 |
|  | *Spc25* | -0.509890575 | 1.97E-13 | 2.26E-10 |
|  | *Cirbp* | -0.517176138 | 1.12E-10 | 9.34E-08 |
|  | *Btg2* | -0.596322294 | 2.15E-10 | 1.52E-07 |
|  | *Ccnd2* | -0.257737663 | 3.83E-10 | 2.51E-07 |
|  | *Gmpr* | -0.333931597 | 1.24E-09 | 7.57E-07 |
|  | *Sphkap* | -0.452711745 | 3.95E-09 | 2.27E-06 |
|  | *Slc2a2* | -0.380040008 | 7.01E-09 | 3.62E-06 |
|  | *Ucn3* | -0.258239181 | 7.10E-09 | 3.62E-06 |
|  | *Sfrp5* | -0.913265258 | 1.40E-08 | 6.44E-06 |
|  | *Psma7* | -0.289392937 | 1.62E-08 | 6.80E-06 |
|  | *Selenow* | -0.448935329 | 1.63E-08 | 6.80E-06 |
|  | *Pdia3* | -0.233100918 | 1.89E-08 | 7.23E-06 |
|  | *Ier2* | -1.343824837 | 3.49E-08 | 1.23E-05 |
|  | *H3f3b* | -0.367751997 | 5.75E-08 | 1.76E-05 |
|  | *Ndufb7* | -0.29689112 | 6.67E-08 | 1.98E-05 |
|  | *A330076H08Rik* | -0.380069116 | 1.27E-07 | 3.66E-05 |
|  | *Gnai2* | -0.25482013 | 1.49E-07 | 4.11E-05 |
|  | *Syt13* | -0.245989332 | 1.52E-07 | 4.11E-05 |
|  | *Nisch* | -0.218681422 | 1.76E-07 | 4.48E-05 |
|  | *Podxl2* | -0.596907078 | 4.20E-07 | 9.42E-05 |
|  | *Gm1821* | -0.208077515 | 4.92E-07 | 0.0001075 |
|  | *Ubc* | -0.245988314 | 7.65E-07 | 0.000163297 |
|  | *Pcx* | -0.620875103 | 8.47E-07 | 0.000176735 |
|  | *Cct7* | -0.403983106 | 1.09E-06 | 0.000217668 |
|  | *Pfdn2* | -0.469236642 | 1.32E-06 | 0.000258618 |
|  | *Pyy* | -0.981583341 | 1.65E-06 | 0.000316136 |
|  | *Prnp* | -0.223205401 | 1.82E-06 | 0.000328252 |
|  | *Ncoa1* | -0.43647533 | 1.82E-06 | 0.000328252 |
|  | *Morf4l1* | -0.466316558 | 2.11E-06 | 0.000371981 |
|  | *Cnbp* | -0.271828497 | 3.08E-06 | 0.000524322 |
|  | *H1f0* | -0.460087774 | 3.26E-06 | 0.000544523 |

**Table S9 Differential expression genes detected by edgeR (continued)**

| **Type** | **Gene symbol** | **log2FoldChange** | **P value** | **P adjust** |
| --- | --- | --- | --- | --- |
| Male specific downregulated T2D genes | *Tmem215* | -0.37990192 | 4.03E-06 | 0.000660417 |
|  | *Pfn1* | -0.382893865 | 4.53E-06 | 0.000705046 |
|  | *Pdx1* | -0.484452804 | 5.75E-06 | 0.000880279 |
|  | *Ivd* | -0.634965849 | 7.69E-06 | 0.001139176 |
|  | *Tmem181b-ps* | -0.356367237 | 9.58E-06 | 0.00134103 |
|  | *Cd81* | -0.718895917 | 1.92E-05 | 0.002480142 |
|  | *Ndufs2* | -0.400512988 | 1.99E-05 | 0.002483024 |
|  | *Fkbp1b* | -0.403004953 | 2.02E-05 | 0.002483024 |
|  | *Ddost* | -0.201999397 | 2.03E-05 | 0.002483024 |
|  | *Eif5* | -0.28333995 | 2.32E-05 | 0.002798186 |
|  | *Zfyve27* | -0.715330788 | 2.51E-05 | 0.002989338 |
|  | *Tmem250-ps* | -0.328041252 | 3.08E-05 | 0.003489967 |
|  | *Cars* | -0.675343612 | 3.39E-05 | 0.003784912 |
|  | *Idh3b* | -0.258583915 | 4.17E-05 | 0.004453395 |
|  | *Csnk1a1* | -0.308601364 | 4.67E-05 | 0.004876044 |
|  | *Fam210b* | -0.465022645 | 5.27E-05 | 0.00537876 |
|  | *Exoc3* | -0.444626972 | 6.58E-05 | 0.006494939 |
|  | *Psap* | -0.206971279 | 8.17E-05 | 0.007732161 |
|  | *Ppp1r1a* | -0.218306297 | 8.65E-05 | 0.008020687 |
|  | *Lamtor1* | -0.353305116 | 9.51E-05 | 0.00860181 |
|  | *Creg1* | -0.268739167 | 9.54E-05 | 0.00860181 |
|  | *Galnt9* | -1.025370165 | 9.55E-05 | 0.00860181 |
|  | *Mt1* | -0.312637019 | 9.69E-05 | 0.008642168 |
|  | *Nipal3* | -0.247055023 | 0.000102563 | 0.009056131 |
|  | *Ap1s1* | -0.403095732 | 0.000113965 | 0.009723333 |
|  | *Swi5* | -0.247513393 | 0.000114355 | 0.009723333 |
|  | *Pura* | -0.218320425 | 0.000118926 | 0.010019284 |
|  | *Prune2* | -0.405986927 | 0.000131097 | 0.010944235 |
|  | *Pdhb* | -0.419837786 | 0.000137836 | 0.011278177 |
|  | *Calm3* | -0.246464218 | 0.000143847 | 0.011587268 |
|  | *Ctsb* | -0.246868369 | 0.000147347 | 0.01176597 |
|  | *Atp6ap1* | -0.254969333 | 0.000150221 | 0.011892038 |
|  | *Pde5a* | -0.460966787 | 0.000161757 | 0.012695827 |
|  | *Ddit4* | -0.658129542 | 0.000177971 | 0.01373367 |
|  | *Yipf4* | -0.440829707 | 0.000194353 | 0.014417441 |
|  | *Atp5c1* | -0.336889417 | 0.000197822 | 0.014417441 |
|  | *Gjd2* | -0.417649998 | 0.000216019 | 0.015497698 |
|  | *Dnm2* | -0.273251334 | 0.000227258 | 0.016177576 |
|  | *Dusp10* | -0.790566073 | 0.000277068 | 0.01927515 |
|  | *Gnmt* | -0.751665902 | 0.000285506 | 0.019712773 |
|  | *Taf9* | -0.508818076 | 0.000294969 | 0.020214148 |
|  | *Hint2* | -0.539806477 | 0.000305041 | 0.020579067 |
|  | *Zfr* | -0.342062308 | 0.000307016 | 0.020579067 |
|  | *Romo1* | -0.28323616 | 0.000320773 | 0.021345378 |
|  | *Calm2* | -0.261143752 | 0.000342208 | 0.022287223 |

**Table S9 Differential expression genes detected by edgeR (continued)**

| **Type** | **Gene symbol** | **log2FoldChange** | **P value** | **P adjust** |
| --- | --- | --- | --- | --- |
| Male specific downregulated T2D genes | *Selenos* | -0.225961591 | 0.000355969 | 0.022700421 |
|  | *Jph3* | -0.572952149 | 0.000406056 | 0.025539838 |
|  | *Cdk12* | -0.373452936 | 0.000489218 | 0.029949925 |
|  | *Reep6* | -0.340794417 | 0.000504236 | 0.030645654 |
|  | *Atp6v1e1* | -0.221014825 | 0.00050885 | 0.030645654 |
|  | *Ywhaz* | -0.28640224 | 0.000513837 | 0.030645654 |
|  | *Cntfr* | -0.826572262 | 0.00053758 | 0.031275528 |
|  | *Capn9* | -0.837581133 | 0.000538118 | 0.031275528 |
|  | *C030006K11Rik* | -0.720307073 | 0.000601811 | 0.033569875 |
|  | *Ate1* | -0.442402425 | 0.00064129 | 0.035137389 |
|  | *Ppy* | -0.94558633 | 0.000642827 | 0.035137389 |
|  | *Cox6a2* | -0.407179249 | 0.000673755 | 0.036181804 |
|  | *Slc35a2* | -0.690376294 | 0.000698826 | 0.037310002 |
|  | *Prmt1* | -0.482245383 | 0.000756701 | 0.039707359 |
|  | *Nudc* | -0.258910946 | 0.00076197 | 0.039756655 |
|  | *T2* | -0.574045544 | 0.000811852 | 0.041649353 |
|  | *Kmt2d* | -0.25786166 | 0.000821707 | 0.041920758 |
|  | *Ctsa* | -0.362236842 | 0.000899907 | 0.045405735 |
|  | *Irs2* | -0.408861391 | 0.000910117 | 0.045669945 |
|  | *Wnk1* | -0.278571769 | 0.000963519 | 0.047788493 |
|  | *Hyal1* | -0.79164195 | 0.000980376 | 0.048143293 |
|  | *Abhd17a* | -0.415758929 | 0.00098724 | 0.048222483 |
|  | *Mrpl37* | -0.630831569 | 0.000995147 | 0.048351508 |
|  | *Atraid* | -0.269598052 | 0.001034886 | 0.049755804 |
| Sex-biased genes in T2D | *Eif2s3y* | 3.793125123 | 1.71E-92 | 7.83E-89 |
|  | *Ddx3y* | 2.719582249 | 1.15E-49 | 3.53E-46 |
|  | *Scg2* | 0.474914416 | 4.92E-49 | 1.13E-45 |
|  | *Uty* | 2.446154025 | 1.37E-39 | 2.51E-36 |
|  | *Necab2* | 1.217450508 | 1.62E-24 | 1.49E-21 |
|  | *Kdm5d* | 1.502361189 | 1.79E-18 | 1.26E-15 |
|  | *Naa20* | 1.27274744 | 1.03E-15 | 6.31E-13 |
|  | *Spp1* | 2.238448221 | 1.46E-14 | 7.87E-12 |
|  | *Sdf2l1* | 0.713889377 | 2.34E-14 | 1.19E-11 |
|  | *Gc* | 1.078946342 | 5.60E-13 | 2.71E-10 |
|  | *Rnf5* | 1.142045169 | 8.71E-13 | 4.00E-10 |
|  | *Nkx2-2* | 0.871222667 | 1.53E-12 | 6.68E-10 |
|  | *P4hb* | 0.368090118 | 2.58E-12 | 1.03E-09 |
|  | *Paip2* | 0.434925835 | 2.71E-11 | 9.58E-09 |
|  | *Cd47* | 0.69523336 | 3.17E-10 | 1.04E-07 |
|  | *Mrps28* | 1.250160401 | 4.12E-10 | 1.30E-07 |
|  | *Tmed2* | 0.537830846 | 3.56E-09 | 1.05E-06 |
|  | *Bambi* | 0.903894003 | 3.96E-09 | 1.14E-06 |
|  | *Itm2c* | 0.521062983 | 5.88E-09 | 1.64E-06 |
|  | *Atn1* | 1.192022558 | 1.18E-08 | 3.18E-06 |
|  | *Rasgrf2* | 1.229157356 | 2.04E-08 | 5.21E-06 |

**Table S9 Differential expression genes detected by edgeR (continued)**

| **Type** | **Gene symbol** | **log2FoldChange** | **P value** | **P adjust** |
| --- | --- | --- | --- | --- |
| Sex-biased genes in T2D | *Tssc4* | 0.420883743 | 4.77E-08 | 1.18E-05 |
|  | *Tm9sf1* | 0.966060594 | 1.41E-07 | 3.32E-05 |
|  | *Hyou1* | 0.521393595 | 2.54E-07 | 5.55E-05 |
|  | *Pdia4* | 0.388382888 | 6.47E-07 | 0.000129172 |
|  | *Copb2* | 0.344222043 | 8.49E-07 | 0.000165924 |
|  | *Selenof* | 0.293979795 | 1.18E-06 | 0.000222021 |
|  | *Syp* | 0.370927806 | 1.96E-06 | 0.000338954 |
|  | *Tent5a* | 0.881513623 | 2.13E-06 | 0.000361856 |
|  | *Ccnd1* | 0.581796472 | 2.55E-06 | 0.000425394 |
|  | *Cst3* | 0.296491232 | 2.96E-06 | 0.000475985 |
|  | *Acly* | 0.255687562 | 3.13E-06 | 0.000487103 |
|  | *Senp3* | 1.054729578 | 3.33E-06 | 0.000509263 |
|  | *Pappa2* | 1.132846347 | 3.80E-06 | 0.000572596 |
|  | *Rnase4* | 0.285240817 | 4.12E-06 | 0.000610213 |
|  | *Kctd13* | 0.848182383 | 4.43E-06 | 0.000646095 |
|  | *Hap1* | 0.453368246 | 4.76E-06 | 0.000672267 |
|  | *Creld2* | 0.373856788 | 7.02E-06 | 0.000933496 |
|  | *Nnt* | 0.772596885 | 9.14E-06 | 0.001165456 |
|  | *Tmbim6* | 0.308209227 | 1.17E-05 | 0.001411826 |
|  | *Dnajc3* | 0.239664053 | 1.25E-05 | 0.001488018 |
|  | *Tspan33* | 0.420091732 | 1.34E-05 | 0.001552261 |
|  | *Spint2* | 0.278092259 | 1.64E-05 | 0.001853452 |
|  | *Tspan13* | 0.215000635 | 2.33E-05 | 0.002607666 |
|  | *Ldlr* | 0.515126791 | 2.56E-05 | 0.002797357 |
|  | *Irak1* | 0.228403934 | 2.96E-05 | 0.003201666 |
|  | *Spcs3* | 0.301288664 | 3.31E-05 | 0.003493573 |
|  | *Dennd6b* | 1.000514767 | 3.49E-05 | 0.003643752 |
|  | *Tram1* | 0.325481118 | 3.56E-05 | 0.003668206 |
|  | *Rab3gap2* | 0.734198022 | 4.02E-05 | 0.004058598 |
|  | *Sdhc* | 0.450677061 | 4.77E-05 | 0.004758522 |
|  | *St18* | 0.359511804 | 4.93E-05 | 0.004814435 |
|  | *Clic4* | 0.638431897 | 5.46E-05 | 0.005170424 |
|  | *Rab34* | 0.504966308 | 5.70E-05 | 0.00533585 |
|  | *Tmem218* | 0.421159427 | 6.22E-05 | 0.005707505 |
|  | *Nbas* | 0.321624102 | 6.56E-05 | 0.005958632 |
|  | *Fbxl16* | 0.531327234 | 8.27E-05 | 0.007162786 |
|  | *Kmt2d* | 0.363594491 | 8.84E-05 | 0.00758455 |
|  | *Derl3* | 0.859386144 | 9.64E-05 | 0.008120661 |
|  | *Unc50* | 0.570283325 | 0.000100112 | 0.008355714 |
|  | *Slc16a10* | 0.410197974 | 0.000103655 | 0.008496923 |
|  | *1700086L19Rik* | 0.754934549 | 0.000111825 | 0.009005801 |
|  | *Csad* | 0.575056091 | 0.000120857 | 0.009648557 |
|  | *Rab27a* | 0.425915325 | 0.000124683 | 0.00986826 |
|  | *Rai1* | 0.789959075 | 0.000133692 | 0.010477058 |
|  | *Vegfa* | 0.587629953 | 0.000135799 | 0.010477058 |

**Table S9 Differential expression genes detected by edgeR (continued)**

| **Type** | **Gene symbol** | **log2FoldChange** | **P value** | **P adjust** |
| --- | --- | --- | --- | --- |
| Sex-biased genes in T2D | *Dtx3* | 0.50299309 | 0.000150092 | 0.01138835 |
|  | *Sdc4* | 0.5475546 | 0.000152615 | 0.011416875 |
|  | *Il1r1* | 0.246163184 | 0.000160769 | 0.011808141 |
|  | *Tap2* | 0.81441129 | 0.000191041 | 0.013702697 |
|  | *6330403K07Rik* | 1.03350089 | 0.000202725 | 0.014207761 |
|  | *Gadd45gip1* | 0.277597701 | 0.000210666 | 0.014542315 |
|  | *Pdia6* | 0.201927459 | 0.000216008 | 0.01477683 |
|  | *Tmem179* | 0.92830023 | 0.00023625 | 0.015832182 |
|  | *Tmem39a* | 0.438652239 | 0.000252501 | 0.016677803 |
|  | *Slc6a19* | 0.787965226 | 0.000256276 | 0.016806241 |
|  | *Foxk1* | 0.699990753 | 0.000267549 | 0.017421066 |
|  | *Arhgap39* | 0.71290296 | 0.000270367 | 0.017480548 |
|  | *Sdhd* | 0.54233869 | 0.000287849 | 0.018480737 |
|  | *Akap8l* | 0.496482281 | 0.000299816 | 0.018983538 |
|  | *Smim26* | 0.678675925 | 0.000306288 | 0.019229624 |
|  | *Lrrc8d* | 0.741145094 | 0.000321917 | 0.01987471 |
|  | *Manf* | 0.203125989 | 0.000325481 | 0.019921579 |
|  | *Cdk16* | 0.593504901 | 0.000372925 | 0.022377957 |
|  | *Vopp1* | 0.861273645 | 0.000403366 | 0.023892305 |
|  | *Ube2d3* | 0.30811553 | 0.000440709 | 0.025608557 |
|  | *Kif12* | 0.299577141 | 0.000444054 | 0.025640646 |
|  | *Sepsecs* | 0.472513426 | 0.000461456 | 0.026478921 |
|  | *Zmym3* | 0.787601253 | 0.00052214 | 0.029230281 |
|  | *Car10* | 0.307287988 | 0.000563795 | 0.030628414 |
|  | *Fgf1* | 0.831210818 | 0.000586488 | 0.031673816 |
|  | *Slc35a5* | 0.640917825 | 0.000654527 | 0.035141584 |
|  | *Arf1* | 0.216914813 | 0.000678699 | 0.036227532 |
|  | *Cops7a* | 0.404905049 | 0.000703654 | 0.036705931 |
|  | *Tnks* | 0.773596231 | 0.000727518 | 0.037524414 |
|  | *Tomm7* | 0.344612079 | 0.000737795 | 0.037841887 |
|  | *Ift57* | 0.635766601 | 0.00077235 | 0.039192114 |
|  | *Brms1l* | 0.775102701 | 0.000773354 | 0.039192114 |
|  | *Ptprn* | 0.266088635 | 0.000794754 | 0.039324569 |
|  | *Serinc1* | 0.282146284 | 0.000800969 | 0.039324569 |
|  | *AI593442* | 1.072851699 | 0.000814244 | 0.039553324 |
|  | *Atxn2l* | 0.63641874 | 0.000827252 | 0.039857221 |
|  | *Sdcbp* | 0.382338174 | 0.000829183 | 0.039857221 |
|  | *Trabd* | 0.34951981 | 0.000852161 | 0.040281012 |
|  | *Atf6* | 0.476911618 | 0.000895714 | 0.041476787 |
|  | *Gm15417* | 0.660198756 | 0.000943905 | 0.043154376 |
|  | *Krt8* | 0.27095051 | 0.000972934 | 0.043910709 |
|  | *Map2k3* | 0.781670765 | 0.000975687 | 0.043910709 |
|  | *Mid2* | 0.704601536 | 0.001022278 | 0.044923298 |
|  | *Cryba2* | 0.694759474 | 0.001022652 | 0.044923298 |
|  | *Rtn4rl1* | 0.709148609 | 0.001028101 | 0.044947595 |

**Table S9 Differential expression genes detected by edgeR (continued)**

| **Type** | **Gene symbol** | **log2FoldChange** | **P value** | **P adjust** |
| --- | --- | --- | --- | --- |
| Sex-biased genes in T2D | *Parp16* | 0.692537875 | 0.00104496 | 0.045403457 |
|  | *Bpnt1* | 0.329439019 | 0.001081637 | 0.046247473 |
|  | *Pitpnc1* | 0.609303209 | 0.001128477 | 0.047525443 |
|  | *Lpar1* | 0.88120675 | 0.001135238 | 0.047591875 |
|  | *Xist* | -4.73990759 | 1.77E-174 | 1.62E-170 |
|  | *Cish* | -2.367956679 | 6.10E-39 | 9.33E-36 |
|  | *Fxyd2* | -3.442725695 | 6.74E-36 | 8.84E-33 |
|  | *Enpp2* | -1.185815282 | 1.66E-26 | 1.90E-23 |
|  | *Prlr* | -0.657221004 | 3.04E-25 | 3.11E-22 |
|  | *G6pc2* | -0.372473576 | 1.97E-20 | 1.64E-17 |
|  | *Jup* | -0.968962557 | 1.00E-19 | 7.68E-17 |
|  | *Spc25* | -0.590777964 | 7.86E-16 | 5.15E-13 |
|  | *Socs2* | -1.298602208 | 9.25E-15 | 5.31E-12 |
|  | *Prss53* | -0.386139168 | 2.10E-12 | 8.74E-10 |
|  | *Gcg* | -0.987816597 | 5.57E-12 | 2.13E-09 |
|  | *Sytl4* | -0.586036702 | 9.38E-12 | 3.44E-09 |
|  | *Chga* | -0.235463886 | 2.80E-10 | 9.53E-08 |
|  | *Rn45s* | -0.209749312 | 6.82E-10 | 2.09E-07 |
|  | *Cebpd* | -1.38486545 | 1.41E-08 | 3.69E-06 |
|  | *Mt1* | -0.451234461 | 1.35E-07 | 3.27E-05 |
|  | *Cd9* | -1.297423282 | 1.58E-07 | 3.63E-05 |
|  | *Rpl23* | -0.330836192 | 2.15E-07 | 4.81E-05 |
|  | *1700001K19Rik* | -1.152983966 | 3.40E-07 | 7.26E-05 |
|  | *Gpx3* | -1.361741919 | 5.35E-07 | 0.000109163 |
|  | *Chst12* | -0.824183544 | 1.08E-06 | 0.000206156 |
|  | *Rps14* | -0.223342238 | 1.39E-06 | 0.000255935 |
|  | *Rpl10* | -0.243381856 | 1.45E-06 | 0.000261131 |
|  | *Pyy* | -0.958040476 | 1.69E-06 | 0.000299199 |
|  | *Selenow* | -0.39148281 | 2.67E-06 | 0.000436995 |
|  | *Galnt9* | -1.143662669 | 3.09E-06 | 0.000487103 |
|  | *Uba52* | -0.284504013 | 4.56E-06 | 0.00065483 |
|  | *Podxl2* | -0.601244528 | 5.29E-06 | 0.000736096 |
|  | *Rpl21* | -0.287620022 | 5.57E-06 | 0.000763185 |
|  | *Hsp90ab1* | -0.240318553 | 7.43E-06 | 0.000974002 |
|  | *Sphkap* | -0.344809857 | 7.83E-06 | 0.001012322 |
|  | *Cox7a2l* | -0.320819585 | 1.03E-05 | 0.001298779 |
|  | *P2ry1* | -0.669066999 | 1.12E-05 | 0.001393501 |
|  | *Epb41l4a* | -1.103461115 | 1.14E-05 | 0.001396621 |
|  | *Ndufa5* | -0.392777256 | 1.30E-05 | 0.001534357 |
|  | *Gadd45g* | -0.504868169 | 2.49E-05 | 0.002753717 |
|  | *Igfbp7* | -0.909268161 | 3.22E-05 | 0.003441161 |
|  | *Slc2a2* | -0.287798416 | 3.75E-05 | 0.003822328 |
|  | *Cited2* | -0.983132355 | 4.83E-05 | 0.004765604 |
|  | *Ppy* | -0.923027915 | 4.99E-05 | 0.004819635 |
|  | *Fmo1* | -1.040790454 | 5.29E-05 | 0.005058148 |

**Table S9 Differential expression genes detected by edgeR (continued)**

| **Type** | **Gene symbol** | **log2FoldChange** | **P value** | **P adjust** |
| --- | --- | --- | --- | --- |
| Sex-biased genes in T2D | *Naca* | -0.287534529 | 6.85E-05 | 0.00616922 |
|  | *1110059G10Rik* | -0.906259778 | 7.20E-05 | 0.006417502 |
|  | *Romo1* | -0.314578645 | 7.73E-05 | 0.006823967 |
|  | *Hsbp1* | -0.349335112 | 7.93E-05 | 0.006933654 |
|  | *Ccdc80* | -0.53799316 | 9.16E-05 | 0.007788791 |
|  | *Mkl2* | -0.895367644 | 0.000102158 | 0.008449644 |
|  | *Hnrnpm* | -0.404043572 | 0.000106504 | 0.008653211 |
|  | *Naaladl1* | -0.731171954 | 0.000138409 | 0.010589477 |
|  | *Trpm5* | -0.449689882 | 0.000152955 | 0.011416875 |
|  | *Rpl36* | -0.231769941 | 0.000160741 | 0.011808141 |
|  | *Nf1* | -0.837846055 | 0.000190804 | 0.013702697 |
|  | *Hist1h2bc* | -0.337443047 | 0.00020212 | 0.014207761 |
|  | *Rpl28* | -0.30508463 | 0.000206241 | 0.014344719 |
|  | *Apobec1* | -0.866754477 | 0.000217283 | 0.01477683 |
|  | *Tmem238* | -0.852216277 | 0.000229834 | 0.015515454 |
|  | *Psma6* | -0.413498559 | 0.000290872 | 0.018545123 |
|  | *Peg3* | -0.40965155 | 0.000307892 | 0.019229624 |
|  | *Prkd1* | -0.769607788 | 0.00032255 | 0.01987471 |
|  | *2210016F16Rik* | -0.439720381 | 0.000366847 | 0.022158044 |
|  | *Cadm1* | -1.088950775 | 0.000383763 | 0.022878731 |
|  | *Eif2s2* | -0.285337167 | 0.000421456 | 0.024803745 |
|  | *Actg1* | -0.231111082 | 0.000425891 | 0.024905133 |
|  | *Hmgn2* | -0.356180322 | 0.000479602 | 0.02718043 |
|  | *Pcbd1* | -0.288059876 | 0.000505593 | 0.028477611 |
|  | *Ap1s2* | -0.338516979 | 0.000530533 | 0.029520162 |
|  | *Rps15a* | -0.269678722 | 0.000536886 | 0.029591655 |
|  | *Rplp1* | -0.218328023 | 0.000538265 | 0.029591655 |
|  | *Wbp2* | -0.29826074 | 0.000549196 | 0.030012886 |
|  | *Echdc2* | -0.493970346 | 0.000696786 | 0.036705931 |
|  | *Cry2* | -0.434157664 | 0.000701371 | 0.036705931 |
|  | *Rlf* | -0.566229773 | 0.000718539 | 0.037270646 |
|  | *Lgi3* | -0.79605568 | 0.000776927 | 0.039192114 |
|  | *Matn2* | -0.757628886 | 0.000789058 | 0.039324569 |
|  | *Metap2* | -0.255636445 | 0.00079814 | 0.039324569 |
|  | *Vps35* | -0.281817781 | 0.000809404 | 0.039527335 |
|  | *Ggact* | -0.523376026 | 0.000852479 | 0.040281012 |
|  | *Mt2* | -0.428240838 | 0.000856364 | 0.040281012 |
|  | *Nipal1* | -0.279896651 | 0.000859937 | 0.040281012 |
|  | *Car15* | -0.780744087 | 0.000899018 | 0.041476787 |
|  | *Ncoa7* | -0.49949626 | 0.00094478 | 0.043154376 |
|  | *1300002E11Rik* | -0.60357943 | 0.000960568 | 0.043658275 |
|  | *Stip1* | -0.406425653 | 0.001009219 | 0.044923298 |
|  | *Cdkn1a* | -0.673887585 | 0.001018622 | 0.044923298 |
|  | *Cmas* | -0.657641313 | 0.001048419 | 0.045403457 |
|  | *Coa3* | -0.208951174 | 0.001072783 | 0.046240468 |

**Table S9 Differential expression genes detected by edgeR (continued)**

| **Type** | **Gene symbol** | **log2FoldChange** | **P value** | **P adjust** |
| --- | --- | --- | --- | --- |
| Sex-biased genes in T2D | *Atp6v1e1* | -0.202465085 | 0.001084811 | 0.046247473 |
|  | *Top1* | -0.302786286 | 0.001092727 | 0.046247473 |
|  | *Akr1c19* | -0.53118042 | 0.001093095 | 0.046247473 |
|  | *Rps21* | -0.22872768 | 0.001164048 | 0.048577823 |
| Sex-biased genes in healthy | *Eif2s3y* | 3.634124448 | 5.66E-58 | 1.70E-54 |
|  | *Scg2* | 0.525378677 | 1.86E-53 | 4.19E-50 |
|  | *Ddx3y* | 3.018192761 | 4.56E-49 | 6.84E-46 |
|  | *Ins2* | 0.291783079 | 2.68E-36 | 3.45E-33 |
|  | *Necab2* | 1.815893888 | 4.18E-36 | 4.70E-33 |
|  | *Uty* | 2.541683218 | 2.97E-31 | 2.97E-28 |
|  | *Malat1* | 0.362744521 | 1.77E-30 | 1.59E-27 |
|  | *Gc* | 2.167691716 | 3.16E-30 | 2.58E-27 |
|  | *Naa20* | 1.888378499 | 3.01E-26 | 2.26E-23 |
|  | *Rnase4* | 0.687038861 | 4.78E-26 | 3.31E-23 |
|  | *Paip2* | 0.617950493 | 6.90E-18 | 3.25E-15 |
|  | *Sfrp5* | 1.452764879 | 5.58E-15 | 1.86E-12 |
|  | *Spp1* | 2.668404228 | 6.79E-15 | 2.18E-12 |
|  | *Tssc4* | 0.616685355 | 2.06E-14 | 6.39E-12 |
|  | *Cpe* | 0.373998457 | 6.06E-14 | 1.70E-11 |
|  | *Syt13* | 0.41967216 | 9.90E-14 | 2.70E-11 |
|  | *Cd47* | 0.862782933 | 1.14E-13 | 3.01E-11 |
|  | *Ccnd1* | 1.23359522 | 1.57E-13 | 4.03E-11 |
|  | *Ubc* | 0.405220022 | 6.69E-13 | 1.54E-10 |
|  | *Mrps28* | 1.619582885 | 7.53E-13 | 1.69E-10 |
|  | *Ucn3* | 0.338453932 | 3.91E-12 | 8.36E-10 |
|  | *Acly* | 0.426464936 | 4.46E-12 | 9.11E-10 |
|  | *Etv1* | 1.665630911 | 7.30E-12 | 1.43E-09 |
|  | *Atn1* | 1.695367277 | 7.85E-12 | 1.50E-09 |
|  | *Nkx2-2* | 0.908458518 | 8.33E-12 | 1.56E-09 |
|  | *Rnf5* | 1.192106805 | 9.51E-12 | 1.74E-09 |
|  | *Tspan33* | 0.640418739 | 1.17E-11 | 2.07E-09 |
|  | *Fxyd6* | 0.892159969 | 2.45E-11 | 4.24E-09 |
|  | *Manf* | 0.40503042 | 7.44E-11 | 1.24E-08 |
|  | *Ttr* | 0.365378508 | 9.91E-11 | 1.62E-08 |
|  | *Ccnd2* | 0.306927404 | 2.38E-10 | 3.69E-08 |
|  | *Hap1* | 0.657944447 | 3.30E-10 | 4.95E-08 |
|  | *Hspa8* | 0.31700855 | 3.37E-10 | 4.96E-08 |
|  | *Scg3* | 0.270611689 | 8.95E-10 | 1.24E-07 |
|  | *Srsf7* | 0.648509611 | 2.46E-09 | 3.16E-07 |
|  | *Pycr2* | 0.636370148 | 2.82E-09 | 3.57E-07 |
|  | *Etfb* | 0.444272525 | 3.05E-09 | 3.75E-07 |
|  | *Alcam* | 0.800811923 | 3.26E-09 | 3.96E-07 |
|  | *Tmod2* | 0.601374859 | 3.74E-09 | 4.46E-07 |
|  | *Bsg* | 0.283853966 | 3.77E-09 | 4.46E-07 |
|  | *Bcl2l2* | 1.336844641 | 5.95E-09 | 6.94E-07 |

**Table S9 Differential expression genes detected by edgeR (continued)**

| **Type** | **Gene symbol** | **log2FoldChange** | **P value** | **P adjust** |
| --- | --- | --- | --- | --- |
| Sexbiased genes in healthy | *A330076H08Rik* | 0.439715973 | 9.63E-09 | 1.06E-06 |
|  | *Bambi* | 1.060962915 | 1.04E-08 | 1.12E-06 |
|  | *Glrx5* | 0.4909659 | 1.51E-08 | 1.62E-06 |
|  | *Hspa1a* | 1.496541745 | 1.95E-08 | 2.04E-06 |
|  | *Chic1* | 0.311490746 | 2.60E-08 | 2.62E-06 |
|  | *Tram1* | 0.484769049 | 2.93E-08 | 2.92E-06 |
|  | *Slc16a10* | 0.663662832 | 3.45E-08 | 3.37E-06 |
|  | *Osgin1* | 1.147089375 | 3.65E-08 | 3.53E-06 |
|  | *Btg2* | 0.576837639 | 4.66E-08 | 4.33E-06 |
|  | *Ndufb7* | 0.343411211 | 4.67E-08 | 4.33E-06 |
|  | *Atp8b2* | 1.153190489 | 1.36E-07 | 1.23E-05 |
|  | *Serp1* | 0.306766007 | 1.44E-07 | 1.28E-05 |
|  | *Tmem160* | 0.458584561 | 1.98E-07 | 1.71E-05 |
|  | *Pam* | 0.40268769 | 3.06E-07 | 2.59E-05 |
|  | *Os9* | 0.362280776 | 3.47E-07 | 2.89E-05 |
|  | *Il1r1* | 0.395500349 | 4.22E-07 | 3.45E-05 |
|  | *Pcp4* | 1.322552098 | 5.81E-07 | 4.66E-05 |
|  | *Mtss1l* | 1.077740228 | 6.36E-07 | 5.01E-05 |
|  | *Insrr* | 0.481456673 | 6.46E-07 | 5.01E-05 |
|  | *Etv5* | 1.241593065 | 6.53E-07 | 5.01E-05 |
|  | *Rab34* | 0.689738307 | 6.63E-07 | 5.01E-05 |
|  | *Edem2* | 0.358752193 | 7.46E-07 | 5.59E-05 |
|  | *Cog2* | 0.609518347 | 8.97E-07 | 6.56E-05 |
|  | *Slc6a17* | 1.019811117 | 1.15E-06 | 8.33E-05 |
|  | *Slc39a7* | 0.389965495 | 1.20E-06 | 8.62E-05 |
|  | *Pdx1* | 0.599406442 | 1.84E-06 | 0.000127886 |
|  | *Itm2c* | 0.427654625 | 2.16E-06 | 0.000148178 |
|  | *Rbm10* | 0.449506166 | 2.37E-06 | 0.000159952 |
|  | *Cyld* | 0.889863031 | 2.48E-06 | 0.00016552 |
|  | *Cox4i1* | 0.202645219 | 2.66E-06 | 0.00017294 |
|  | *Kmt2d* | 0.438448819 | 2.73E-06 | 0.00017627 |
|  | *Fam234a* | 0.587869901 | 3.07E-06 | 0.000192669 |
|  | *Aplp1* | 0.350467345 | 4.80E-06 | 0.000291744 |
|  | *Selenop* | 0.314854134 | 5.44E-06 | 0.000319514 |
|  | *Jam2* | 1.230481666 | 5.51E-06 | 0.000321697 |
|  | *Cyth1* | 0.998238017 | 6.14E-06 | 0.000351748 |
|  | *Dapl1* | 0.895475059 | 6.21E-06 | 0.00035345 |
|  | *Dnajb1* | 0.810749009 | 6.48E-06 | 0.000365058 |
|  | *Vwa5b2* | 0.743828131 | 6.54E-06 | 0.000365058 |
|  | *Ddx50* | 0.496489605 | 8.71E-06 | 0.000480331 |
|  | *Fgf1* | 1.104827588 | 8.80E-06 | 0.000482106 |
|  | *Hnrnpa2b1* | 0.348005637 | 1.01E-05 | 0.000544953 |
|  | *Nbeal1* | 0.714281821 | 1.07E-05 | 0.000571269 |
|  | *Canx* | 0.272475503 | 1.11E-05 | 0.000591944 |
|  | *Cd44* | 1.007784364 | 1.17E-05 | 0.000621164 |

**Table S9 Differential expression genes detected by edgeR (continued)**

| **Type** | **Gene symbol** | **log2FoldChange** | **P value** | **P adjust** |
| --- | --- | --- | --- | --- |
| Sex-biased genes in healthy | *Akr1c14* | 0.904488617 | 1.22E-05 | 0.000638555 |
|  | *Prrc2b* | 0.539325792 | 1.23E-05 | 0.000638555 |
|  | *Whrn* | 1.03651079 | 1.24E-05 | 0.000638555 |
|  | *Fam189b* | 0.732095204 | 1.24E-05 | 0.000638555 |
|  | *Bag2* | 0.880498748 | 1.51E-05 | 0.000764731 |
|  | *Serinc1* | 0.370476666 | 1.73E-05 | 0.000869355 |
|  | *Cfp* | 1.174240218 | 1.84E-05 | 0.000920288 |
|  | *Npff* | 1.028225876 | 1.96E-05 | 0.00097369 |
|  | *Selenom* | 0.327963977 | 2.10E-05 | 0.001031907 |
|  | *Pappa2* | 1.368093773 | 2.12E-05 | 0.001036655 |
|  | *Scn3a* | 1.22252477 | 2.25E-05 | 0.001087457 |
|  | *Rgs16* | 0.898944575 | 2.60E-05 | 0.001236108 |
|  | *Ang* | 0.65712061 | 2.64E-05 | 0.001245895 |
|  | *Tap2* | 1.041791981 | 2.65E-05 | 0.001245895 |
|  | *Phf1* | 0.788876557 | 2.73E-05 | 0.001279448 |
|  | *Pitpnc1* | 0.853295103 | 2.78E-05 | 0.00129421 |
|  | *Pura* | 0.264238012 | 2.81E-05 | 0.001295133 |
|  | *Abcc8* | 0.260614207 | 2.95E-05 | 0.001350823 |
|  | *Egln2* | 0.628799695 | 3.17E-05 | 0.001448513 |
|  | *Ppp1r15a* | 0.875463387 | 3.39E-05 | 0.001540601 |
|  | *Abcb4* | 0.935601285 | 3.67E-05 | 0.001650889 |
|  | *Cdk12* | 0.495572247 | 4.00E-05 | 0.001787205 |
|  | *Sel1l3* | 1.070997749 | 4.05E-05 | 0.001800955 |
|  | *Vat1* | 0.454163857 | 5.15E-05 | 0.002235906 |
|  | *Tspan13* | 0.230035703 | 5.31E-05 | 0.002295972 |
|  | *Cacybp* | 0.534825529 | 5.42E-05 | 0.002332982 |
|  | *Gnao1* | 0.494560341 | 5.60E-05 | 0.002385461 |
|  | *Bmi1* | 0.833461306 | 5.93E-05 | 0.00251439 |
|  | *Bhlha15* | 0.7693416 | 6.10E-05 | 0.002572737 |
|  | *Rgs11* | 0.49961024 | 6.13E-05 | 0.002576285 |
|  | *H1f0* | 0.436431421 | 6.49E-05 | 0.002711284 |
|  | *Ankrd10* | 0.571938189 | 6.91E-05 | 0.002877338 |
|  | *Ripk4* | 0.885872988 | 7.23E-05 | 0.002982121 |
|  | *Psma7* | 0.226869859 | 7.27E-05 | 0.002983789 |
|  | *Ctsd* | 0.307424281 | 7.78E-05 | 0.003148457 |
|  | *Cflar* | 0.741237238 | 7.95E-05 | 0.003204728 |
|  | *Sephs2* | 0.613025774 | 8.14E-05 | 0.003265908 |
|  | *Ccdc107* | 0.951837019 | 8.47E-05 | 0.003383044 |
|  | *Pabpn1* | 0.532390932 | 8.54E-05 | 0.003394563 |
|  | *Tm9sf1* | 0.815076684 | 8.99E-05 | 0.003544489 |
|  | *Slc4a7* | 0.433683517 | 9.60E-05 | 0.00375176 |
|  | *Nucb1* | 0.416708116 | 9.96E-05 | 0.003874329 |
|  | *Napa* | 0.272398963 | 0.000100068 | 0.003876767 |
|  | *Chchd10* | 0.388195626 | 0.000103936 | 0.003975207 |
|  | *Map4* | 0.935676004 | 0.000104951 | 0.003997034 |

**Table S9 Differential expression genes detected by edgeR (continued)**

| **Type** | **Gene symbol** | **log2FoldChange** | **P value** | **P adjust** |
| --- | --- | --- | --- | --- |
| Sex-biased genes in healthy | *Araf* | 0.464242197 | 0.000110713 | 0.004196027 |
|  | *Atp2a2* | 0.266421772 | 0.000112203 | 0.004217001 |
|  | *Hsd17b12* | 0.546070878 | 0.000112604 | 0.004217001 |
|  | *Spcs2* | 0.20749603 | 0.000118714 | 0.004427395 |
|  | *Golgb1* | 0.208891355 | 0.00011938 | 0.004433823 |
|  | *Psap* | 0.233232949 | 0.000121367 | 0.004489085 |
|  | *Ptbp3* | 0.521427993 | 0.000137224 | 0.005040092 |
|  | *Fam19a1* | 0.849815674 | 0.000138599 | 0.00505221 |
|  | *Rasgrf2* | 1.0366918 | 0.00013884 | 0.00505221 |
|  | *Cst3* | 0.272079521 | 0.000142317 | 0.005157837 |
|  | *Ctsa* | 0.483996358 | 0.000149567 | 0.005377224 |
|  | *Atp6v0b* | 0.264285537 | 0.000156519 | 0.005563011 |
|  | *Fuom* | 0.655360419 | 0.000161702 | 0.005721949 |
|  | *Senp3* | 0.921288271 | 0.000167722 | 0.005911708 |
|  | *Actb* | 0.223581823 | 0.000171563 | 0.006000038 |
|  | *Tmem201* | 0.711048951 | 0.000178651 | 0.00619237 |
|  | *Irgm1* | 0.962561193 | 0.00017913 | 0.00619237 |
|  | *Cfap20* | 0.550782479 | 0.000190854 | 0.006547321 |
|  | *Selenot* | 0.434530647 | 0.000209469 | 0.007131462 |
|  | *Depp1* | 1.01374869 | 0.000217952 | 0.007364488 |
|  | *Mafk* | 0.63000828 | 0.000222437 | 0.007459932 |
|  | *Gnb2* | 0.407287232 | 0.000232505 | 0.007768603 |
|  | *Gria2* | 0.891460475 | 0.000240025 | 0.007960682 |
|  | *Smad2* | 0.992417849 | 0.000244728 | 0.008086808 |
|  | *Fam174b* | 0.248798951 | 0.000247261 | 0.008140608 |
|  | *Chchd4* | 0.436136162 | 0.000262087 | 0.008567041 |
|  | *Hspa13* | 0.446372267 | 0.000271743 | 0.008849377 |
|  | *Nudt3* | 0.351492061 | 0.000273106 | 0.008861663 |
|  | *Gabarapl1* | 0.205924748 | 0.000277369 | 0.008967607 |
|  | *Zfp106* | 0.299205476 | 0.00027907 | 0.00898762 |
|  | *Selenof* | 0.242660887 | 0.000304904 | 0.009642016 |
|  | *Kif12* | 0.311975979 | 0.00030543 | 0.009642016 |
|  | *Cltb* | 0.234538025 | 0.00030646 | 0.009642016 |
|  | *Trap1* | 0.841982139 | 0.000306811 | 0.009642016 |
|  | *Pamr1* | 0.653382667 | 0.000308286 | 0.009654603 |
|  | *Klc4* | 0.552710286 | 0.000310319 | 0.009662573 |
|  | *Scnn1g* | 0.809054597 | 0.00031659 | 0.009780625 |
|  | *Rfng* | 0.862152168 | 0.000316662 | 0.009780625 |
|  | *Sgcd* | 0.959048671 | 0.000320244 | 0.009857373 |
|  | *Glul* | 0.540171678 | 0.000330912 | 0.010150984 |
|  | *Zfp580* | 0.991815815 | 0.000353857 | 0.010762395 |
|  | *Slc48a1* | 0.400617152 | 0.00038709 | 0.011558702 |
|  | *Syngr1* | 0.798409303 | 0.000391237 | 0.011643841 |
|  | *Mbnl2* | 0.33525222 | 0.000410365 | 0.012092978 |
|  | *Hsp90aa1* | 0.288550297 | 0.000420394 | 0.012337001 |

**Table S9 Differential expression genes detected by edgeR (continued)**

| **Type** | **Gene symbol** | **log2FoldChange** | **P value** | **P adjust** |
| --- | --- | --- | --- | --- |
| Sex-biased genes in healthy | *Wwox* | 0.851860531 | 0.000426267 | 0.012439236 |
|  | *Vps37a* | 0.46643888 | 0.000429781 | 0.012442389 |
|  | *Tmem59* | 0.22779921 | 0.000430528 | 0.012442389 |
|  | *Dnm2* | 0.296270858 | 0.000442054 | 0.012734544 |
|  | *Tmed2* | 0.324730008 | 0.000448068 | 0.012866574 |
|  | *Arhgef1* | 0.744388801 | 0.000457308 | 0.01309009 |
|  | *Atp2a3* | 0.298948026 | 0.000476234 | 0.013551497 |
|  | *Ncstn* | 0.490270839 | 0.000476443 | 0.013551497 |
|  | *Gorasp1* | 0.753589551 | 0.000485112 | 0.013754539 |
|  | *Cct7* | 0.321437893 | 0.000494047 | 0.013920044 |
|  | *Edem3* | 0.76774925 | 0.0005027 | 0.014119584 |
|  | *Sgcz* | 1.234987736 | 0.000548152 | 0.015290865 |
|  | *Iffo1* | 0.619085816 | 0.00056597 | 0.015652108 |
|  | *Nap1l5* | 0.691020413 | 0.000568966 | 0.015686696 |
|  | *Eif4enif1* | 0.88306256 | 0.000576895 | 0.015856674 |
|  | *Asic1* | 0.79205142 | 0.000586586 | 0.016048994 |
|  | *Sec31a* | 0.248746822 | 0.000590316 | 0.016078049 |
|  | *Tmem191c* | 0.875959848 | 0.000596597 | 0.016200034 |
|  | *Ssbp4* | 0.714742762 | 0.000614683 | 0.016640883 |
|  | *Pi4ka* | 0.553816729 | 0.000620696 | 0.01671282 |
|  | *Igtp* | 0.784882138 | 0.000621059 | 0.01671282 |
|  | *Arhgap18* | 0.96383419 | 0.000636692 | 0.017082341 |
|  | *Reep6* | 0.387027259 | 0.000646616 | 0.017296988 |
|  | *Arl2* | 0.541213468 | 0.000649163 | 0.01731357 |
|  | *Atf6* | 0.56310726 | 0.000672586 | 0.017832444 |
|  | *Clcn3* | 0.417973358 | 0.000716559 | 0.018942454 |
|  | *Ube2d3* | 0.327278958 | 0.000720572 | 0.018992673 |
|  | *Thap1* | 0.89011444 | 0.000730019 | 0.019142007 |
|  | *Nxt2* | 0.768698707 | 0.000730497 | 0.019142007 |
|  | *Tob2* | 0.767437749 | 0.000743544 | 0.019427243 |
|  | *Hdlbp* | 0.275312584 | 0.000775927 | 0.020040328 |
|  | *Nemf* | 0.37119449 | 0.00078578 | 0.020236641 |
|  | *Pcyox1* | 0.393637962 | 0.000789976 | 0.020286573 |
|  | *Fbxo2* | 0.632065644 | 0.000803206 | 0.020567556 |
|  | *Eral1* | 0.770393149 | 0.000834005 | 0.021235228 |
|  | *Uqcr11* | 0.258137224 | 0.000837442 | 0.02126251 |
|  | *Nfyb* | 0.657471069 | 0.000856338 | 0.021678433 |
|  | *Tcim* | 0.849634411 | 0.000869918 | 0.021901463 |
|  | *Gnmt* | 0.726564449 | 0.000874468 | 0.021954508 |
|  | *Plekhb1* | 0.914489397 | 0.000910242 | 0.022647031 |
|  | *Fem1a* | 0.639110687 | 0.000913593 | 0.022647031 |
|  | *Rsph9* | 0.834207755 | 0.000914286 | 0.022647031 |
|  | *Selenos* | 0.231472211 | 0.000915102 | 0.022647031 |
|  | *Ap1s1* | 0.391323135 | 0.000917169 | 0.022647031 |
|  | *Hid1* | 0.35317319 | 0.000931057 | 0.022926966 |

**Table S9 Differential expression genes detected by edgeR (continued)**

| **Type** | **Gene symbol** | **log2FoldChange** | **P value** | **P adjust** |
| --- | --- | --- | --- | --- |
| Sex-biased genes in healthy | *Mrpl3* | 0.826903286 | 0.000954125 | 0.023303469 |
|  | *Selenon* | 0.910834014 | 0.000962342 | 0.023440457 |
|  | *Pgap1* | 0.666409076 | 0.000965287 | 0.023448638 |
|  | *Rheb* | 0.375819903 | 0.000993046 | 0.02399032 |
|  | *Gsn* | 0.498754673 | 0.000993945 | 0.02399032 |
|  | *Rap1b* | 0.573799988 | 0.000995593 | 0.02399032 |
|  | *Madd* | 0.409657739 | 0.001007588 | 0.024214452 |
|  | *Lima1* | 0.797997534 | 0.001014555 | 0.024223842 |
|  | *Flnb* | 0.408333714 | 0.00102335 | 0.024332981 |
|  | *Ptpn1* | 0.378361977 | 0.001050567 | 0.024914245 |
|  | *Lta4h* | 0.748909143 | 0.001083367 | 0.025557221 |
|  | *Eif2ak1* | 0.768206862 | 0.001087564 | 0.025589071 |
|  | *Tuba4a* | 0.59748024 | 0.001115512 | 0.02617812 |
|  | *Bpnt1* | 0.365295975 | 0.001199828 | 0.027865764 |
|  | *Mpp2* | 0.914105788 | 0.001227049 | 0.028424537 |
|  | *Rbm8a* | 0.46946504 | 0.001232013 | 0.028466149 |
|  | *Zfp956* | 0.899413954 | 0.001242925 | 0.028644638 |
|  | *Rab7* | 0.241822252 | 0.001312656 | 0.02996244 |
|  | *Rragb* | 1.143649322 | 0.001331273 | 0.030239651 |
|  | *Degs2* | 0.803854199 | 0.001332321 | 0.030239651 |
|  | *Rexo2* | 0.343386951 | 0.001347259 | 0.030501673 |
|  | *Epm2aip1* | 0.676715347 | 0.001410174 | 0.031686599 |
|  | *Bach1* | 0.844494958 | 0.001422567 | 0.031885373 |
|  | *Eif4a1* | 0.220022909 | 0.001445802 | 0.03224532 |
|  | *Gca* | 0.891752933 | 0.001465633 | 0.032606712 |
|  | *Nenf* | 0.388808617 | 0.001475593 | 0.032747239 |
|  | *Trabd* | 0.382205466 | 0.001495103 | 0.032911057 |
|  | *Foxk2* | 0.820846673 | 0.001497571 | 0.032911057 |
|  | *Rai1* | 0.705183512 | 0.001504888 | 0.032990085 |
|  | *Por* | 0.355581616 | 0.001603311 | 0.034808108 |
|  | *Gramd1a* | 0.42420781 | 0.001632788 | 0.035362652 |
|  | *Man2b1* | 0.561110495 | 0.001640639 | 0.035447272 |
|  | *Tmem65* | 0.583251641 | 0.001647326 | 0.0355064 |
|  | *Pes1* | 0.717040334 | 0.001727414 | 0.03685217 |
|  | *Hsph1* | 0.5337743 | 0.001731537 | 0.03685217 |
|  | *Pisd* | 0.457070889 | 0.001736691 | 0.03685217 |
|  | *Fam118a* | 0.759033168 | 0.001738465 | 0.03685217 |
|  | *Snx17* | 0.481769448 | 0.00174907 | 0.036989745 |
|  | *Senp5* | 0.648284911 | 0.001769171 | 0.037327012 |
|  | *Acadl* | 0.560030817 | 0.001796448 | 0.037813761 |
|  | *Zfr* | 0.327430762 | 0.00185902 | 0.038948412 |
|  | *Dnaja1* | 0.310310377 | 0.00186465 | 0.038953983 |
|  | *Idh3b* | 0.229802463 | 0.001867954 | 0.038953983 |
|  | *Aplp2* | 0.515629933 | 0.001897711 | 0.03938954 |
|  | *Smu1* | 0.577341404 | 0.001902538 | 0.03938954 |

**Table S9 Differential expression genes detected by edgeR (continued)**

| **Type** | **Gene symbol** | **log2FoldChange** | **P value** | **P adjust** |
| --- | --- | --- | --- | --- |
| Sex-biased genes in healthy | *Bclaf1* | 0.353331805 | 0.00190637 | 0.03938954 |
|  | *Nr3c1* | 0.475100354 | 0.001919088 | 0.039561377 |
|  | *Mospd2* | 0.849167557 | 0.001927078 | 0.039635197 |
|  | *Tmbim1* | 0.644514272 | 0.001943712 | 0.03979438 |
|  | *Map7* | 0.473439186 | 0.001947813 | 0.03979438 |
|  | *Rab18* | 0.385650007 | 0.0019481 | 0.03979438 |
|  | *Tpm4* | 0.616425297 | 0.001954157 | 0.039827589 |
|  | *Kctd13* | 0.670566202 | 0.001964478 | 0.039947343 |
|  | *Chd5* | 0.637384578 | 0.001998525 | 0.040407812 |
|  | *Helq* | 0.916218304 | 0.002018768 | 0.040683155 |
|  | *Irs2* | 0.424077477 | 0.002028807 | 0.040793993 |
|  | *Dhx15* | 0.564170938 | 0.002037798 | 0.040883321 |
|  | *Nsa2* | 0.73662471 | 0.002088456 | 0.041713434 |
|  | *Cyfip2* | 0.300620355 | 0.002141771 | 0.042683459 |
|  | *Ankrd54* | 0.494078405 | 0.002157803 | 0.042907808 |
|  | *Dennd2d* | 0.803072805 | 0.00216767 | 0.043008866 |
|  | *Tm4sf4* | 0.228140496 | 0.002174704 | 0.043053395 |
|  | *Lmna* | 0.355076417 | 0.002200379 | 0.043302186 |
|  | *Mtmr11* | 0.655927002 | 0.002285289 | 0.044749836 |
|  | *Ckb* | 0.415334636 | 0.002376456 | 0.046333163 |
|  | *Snrnp70* | 0.319766437 | 0.002423139 | 0.047039242 |
|  | *Bcas3* | 0.810106601 | 0.002434919 | 0.047166053 |
|  | *Rbm12b2* | 0.728866499 | 0.002452625 | 0.047320232 |
|  | *Spc25* | 0.240423942 | 0.002453408 | 0.047320232 |
|  | *Eea1* | 0.502408475 | 0.002488373 | 0.047822572 |
|  | *Atp2b1* | 0.290913334 | 0.002490094 | 0.047822572 |
|  | *Nr4a1* | 0.907284115 | 0.002505854 | 0.048022635 |
|  | *Fbxo44* | 0.599459253 | 0.002532872 | 0.048437127 |
|  | *Mgat2* | 0.510804926 | 0.002550824 | 0.048471052 |
|  | *Sobp* | 0.8132397 | 0.002583036 | 0.048979599 |
|  | *Zeb1* | 0.839953787 | 0.002622262 | 0.049307296 |
|  | *Xist* | -4.551614726 | 1.11E-143 | 9.98E-140 |
|  | *Iapp* | -0.552825803 | 3.70E-63 | 1.66E-59 |
|  | *Jup* | -1.706444531 | 8.91E-53 | 1.60E-49 |
|  | *Pabpc1* | -0.957612124 | 5.01E-24 | 3.22E-21 |
|  | *Enpp2* | -1.117048429 | 7.94E-24 | 4.76E-21 |
|  | *Gcg* | -1.776175085 | 3.81E-21 | 2.14E-18 |
|  | *Fmo1* | -2.166347895 | 3.04E-19 | 1.61E-16 |
|  | *Sytl4* | -0.798420153 | 4.98E-18 | 2.48E-15 |
|  | *Naaladl1* | -1.542084825 | 7.24E-18 | 3.25E-15 |
|  | *Rps27a* | -0.555924727 | 2.59E-17 | 1.11E-14 |
|  | *Prss53* | -0.480166937 | 1.42E-16 | 5.79E-14 |
|  | *Nf1* | -2.137579076 | 6.38E-16 | 2.49E-13 |
|  | *Itga11* | -1.81695849 | 6.94E-16 | 2.60E-13 |
|  | *Cish* | -1.78646843 | 1.54E-15 | 5.54E-13 |

**Table S9 Differential expression genes detected by edgeR (continued)**

| **Type** | **Gene symbol** | **log2FoldChange** | **P value** | **P adjust** |
| --- | --- | --- | --- | --- |
| Sex-biased genes in healthy | *Rplp1* | -0.588934211 | 4.23E-15 | 1.46E-12 |
|  | *Prlr* | -0.497275396 | 2.79E-14 | 8.35E-12 |
|  | *Matn2* | -1.572484966 | 5.33E-14 | 1.54E-11 |
|  | *Cdh12* | -1.534689509 | 1.63E-13 | 4.08E-11 |
|  | *Rpl23* | -0.491662241 | 5.25E-13 | 1.27E-10 |
|  | *Rpl10* | -0.391367527 | 5.35E-13 | 1.27E-10 |
|  | *Rps24* | -0.547574333 | 1.22E-12 | 2.68E-10 |
|  | *Rpl21* | -0.481304829 | 4.08E-12 | 8.54E-10 |
|  | *Rps26* | -0.771357944 | 6.40E-12 | 1.28E-09 |
|  | *Grem2* | -1.69970547 | 1.14E-11 | 2.05E-09 |
|  | *Gnb1* | -0.939398553 | 5.53E-11 | 9.37E-09 |
|  | *Hsp90ab1* | -0.386268863 | 1.78E-10 | 2.85E-08 |
|  | *Degs1* | -0.48221624 | 2.04E-10 | 3.22E-08 |
|  | *mt-Nd1* | -0.252228333 | 2.62E-10 | 3.99E-08 |
|  | *Mt2* | -0.756249441 | 4.48E-10 | 6.50E-08 |
|  | *Rpl41* | -0.226179267 | 6.55E-10 | 9.34E-08 |
|  | *Mt1* | -0.533708639 | 7.70E-10 | 1.08E-07 |
|  | *Ssr4* | -0.376918828 | 1.08E-09 | 1.46E-07 |
|  | *4732471J01Rik* | -1.139744475 | 1.09E-09 | 1.46E-07 |
|  | *Rps25* | -0.651406995 | 1.73E-09 | 2.28E-07 |
|  | *Shisal2b* | -1.410572343 | 2.14E-09 | 2.79E-07 |
|  | *Rps29* | -0.459508425 | 2.99E-09 | 3.73E-07 |
|  | *1700001K19Rik* | -1.335542123 | 6.71E-09 | 7.73E-07 |
|  | *Isg20* | -0.586632837 | 7.27E-09 | 8.28E-07 |
|  | *Lgi3* | -1.057717875 | 7.62E-09 | 8.56E-07 |
|  | *Rpsa* | -0.525102409 | 9.06E-09 | 1.01E-06 |
|  | *Rpl6* | -0.39473644 | 1.66E-08 | 1.75E-06 |
|  | *Rpl15* | -0.542018025 | 2.02E-08 | 2.09E-06 |
|  | *Trpm5* | -0.820369994 | 2.57E-08 | 2.62E-06 |
|  | *Slc25a3* | -0.387197279 | 3.10E-08 | 3.06E-06 |
|  | *Rps12* | -0.64586775 | 4.15E-08 | 3.96E-06 |
|  | *Zbtb20* | -0.748847426 | 4.52E-08 | 4.28E-06 |
|  | *Prkcb* | -0.622732548 | 8.59E-08 | 7.88E-06 |
|  | *Arglu1* | -0.981445338 | 1.34E-07 | 1.22E-05 |
|  | *Epb41l4a* | -1.405081949 | 1.84E-07 | 1.60E-05 |
|  | *Kidins220* | -0.631566452 | 2.46E-07 | 2.10E-05 |
|  | *Rps17* | -0.508843766 | 3.27E-07 | 2.75E-05 |
|  | *Tceal9* | -0.25703482 | 4.03E-07 | 3.32E-05 |
|  | *Sh3pxd2a* | -0.597968391 | 5.02E-07 | 4.07E-05 |
|  | *Myo1b* | -1.276316573 | 5.93E-07 | 4.72E-05 |
|  | *Usmg5* | -0.340302953 | 6.61E-07 | 5.01E-05 |
|  | *Hist1h2bc* | -0.490379403 | 6.63E-07 | 5.01E-05 |
|  | *Rps21* | -0.411092205 | 7.57E-07 | 5.63E-05 |
|  | *D16Ertd472e* | -1.079757432 | 8.10E-07 | 5.97E-05 |
|  | *Wnt4* | -0.719109162 | 1.37E-06 | 9.79E-05 |

**Table S9 Differential expression genes detected by edgeR (continued)**

| **Type** | **Gene symbol** | **log2FoldChange** | **P value** | **P adjust** |
| --- | --- | --- | --- | --- |
| Sex-biased genes in healthy | *Tpt1* | -0.3203774 | 1.40E-06 | 9.93E-05 |
|  | *Rpl19* | -0.340942612 | 1.74E-06 | 0.000121843 |
|  | *Tmed3* | -0.273896684 | 2.12E-06 | 0.000146571 |
|  | *AW822252* | -1.271392152 | 2.28E-06 | 0.000154926 |
|  | *Rpl37a* | -0.25916765 | 2.49E-06 | 0.00016552 |
|  | *Pdyn* | -1.43466046 | 2.50E-06 | 0.00016552 |
|  | *Chac1* | -1.135881132 | 2.58E-06 | 0.000169357 |
|  | *Gapdh* | -0.377570056 | 2.75E-06 | 0.00017627 |
|  | *Dap* | -0.323939188 | 2.78E-06 | 0.000176987 |
|  | *Meis2* | -0.610143205 | 2.92E-06 | 0.000185011 |
|  | *Rps14* | -0.245118154 | 3.87E-06 | 0.000241459 |
|  | *Eif4ebp1* | -0.96772861 | 4.08E-06 | 0.000252641 |
|  | *Mid1ip1* | -0.703020866 | 4.19E-06 | 0.000257722 |
|  | *Phactr1* | -0.378508478 | 4.78E-06 | 0.000291744 |
|  | *Cox7b* | -0.490106051 | 5.08E-06 | 0.000306144 |
|  | *Socs2* | -0.815963794 | 5.13E-06 | 0.000307477 |
|  | *Rps13* | -0.381168376 | 5.18E-06 | 0.000308372 |
|  | *Cebpd* | -1.231144331 | 5.38E-06 | 0.000318197 |
|  | *Rpl27a* | -0.503753208 | 5.74E-06 | 0.000333105 |
|  | *2310022B05Rik* | -0.885711225 | 5.87E-06 | 0.000338212 |
|  | *Rps3a1* | -0.343829452 | 6.51E-06 | 0.000365058 |
|  | *Mir682* | -0.3306284 | 7.94E-06 | 0.00044061 |
|  | *Eif3m* | -0.538418948 | 8.96E-06 | 0.000487976 |
|  | *Sec61b* | -0.227193297 | 9.18E-06 | 0.000496847 |
|  | *Pebp1* | -0.265519948 | 1.26E-05 | 0.000648704 |
|  | *Atf4* | -0.484273272 | 1.28E-05 | 0.000652719 |
|  | *Mthfd1* | -0.502043977 | 1.47E-05 | 0.000744235 |
|  | *Fmn2* | -0.796978223 | 2.03E-05 | 0.001000598 |
|  | *Ppia* | -0.378045282 | 2.21E-05 | 0.001072319 |
|  | *Apobec1* | -1.123746113 | 2.51E-05 | 0.001204773 |
|  | *Tnfrsf9* | -1.125569662 | 2.58E-05 | 0.001233822 |
|  | *Rpl12* | -0.520634196 | 2.79E-05 | 0.00129481 |
|  | *Rpl8* | -0.274460167 | 3.42E-05 | 0.001542812 |
|  | *Tmem106c* | -0.586812102 | 4.29E-05 | 0.001901643 |
|  | *Rps3* | -0.337843699 | 4.41E-05 | 0.001942027 |
|  | *Klkb1* | -0.936159225 | 4.70E-05 | 0.002061018 |
|  | *Tpt1* | -0.3203774 | 1.40E-06 | 9.93E-05 |
|  | *Abr* | -0.722564535 | 5.60E-05 | 0.002385461 |
|  | *Eif5b* | -0.407058246 | 7.00E-05 | 0.002897329 |
|  | *Rpl4* | -0.200502213 | 7.65E-05 | 0.003126333 |
|  | *Rps15a* | -0.359782583 | 7.73E-05 | 0.003145274 |
|  | *Rpl38* | -0.248697449 | 8.88E-05 | 0.003516522 |
|  | *Erh* | -0.518938688 | 9.05E-05 | 0.003553203 |
|  | *Arpc5* | -0.330396972 | 0.000100582 | 0.003879955 |
|  | *Scgn* | -0.318414326 | 0.000103843 | 0.003975207 |

**Table S9 Differential expression genes detected by edgeR (continued)**

| **Type** | **Gene symbol** | **log2FoldChange** | **P value** | **P adjust** |
| --- | --- | --- | --- | --- |
| Sex-biased genes in healthy | *Cox6a2* | -0.426666559 | 0.00011111 | 0.004196027 |
|  | *Ndufb3* | -0.441180157 | 0.000137386 | 0.005040092 |
|  | *Utrn* | -0.434146472 | 0.000146834 | 0.005300187 |
|  | *2900055J20Rik* | -0.43691275 | 0.000156591 | 0.005563011 |
|  | *Rpl11* | -0.263822836 | 0.000168879 | 0.005929253 |
|  | *Tma7* | -0.26881367 | 0.000178661 | 0.00619237 |
|  | *Cox7a2l* | -0.311054359 | 0.000182094 | 0.00627072 |
|  | *Gpx3* | -1.198381135 | 0.000200495 | 0.006851894 |
|  | *Ndufa4* | -0.251662394 | 0.000214289 | 0.007268035 |
|  | *Hsbp1* | -0.326505525 | 0.000237664 | 0.007911561 |
|  | *Tnk2* | -1.034504474 | 0.00026212 | 0.008567041 |
|  | *S100a10* | -0.895154211 | 0.000279988 | 0.00898762 |
|  | *Stxbp5l* | -1.136240393 | 0.000298065 | 0.009533845 |
|  | *Aars* | -0.420951774 | 0.000303492 | 0.009642016 |
|  | *Fau* | -0.278851464 | 0.00031069 | 0.009662573 |
|  | *Actg1* | -0.225732499 | 0.000354436 | 0.010762395 |
|  | *Mettl7a1* | -0.694290214 | 0.000357646 | 0.010823318 |
|  | *Rps11* | -0.306515518 | 0.000362956 | 0.010947141 |
|  | *Uqcrb* | -0.29557026 | 0.00037271 | 0.011203733 |
|  | *Rgs2* | -0.524176225 | 0.000376775 | 0.011288172 |
|  | *Trib1* | -0.631938653 | 0.000399587 | 0.011853105 |
|  | *Ptch1* | -0.816956101 | 0.000402065 | 0.011887357 |
|  | *Hint1* | -0.337073993 | 0.000421391 | 0.012337001 |
|  | *Gm11423* | -0.944253917 | 0.000429309 | 0.012442389 |
|  | *Ankrd12* | -0.370645695 | 0.000493328 | 0.013920044 |
|  | *Rps7* | -0.272884167 | 0.000532225 | 0.014902302 |
|  | *Rps5* | -0.247689051 | 0.000549505 | 0.015290865 |
|  | *Vps50* | -0.710090236 | 0.000560534 | 0.015549635 |
|  | *Psmb1* | -0.385710049 | 0.000587463 | 0.016048994 |
|  | *Igsf9b* | -0.742043938 | 0.000763466 | 0.01983247 |
|  | *Nebl* | -0.846017493 | 0.000770649 | 0.019961363 |
|  | *Col6a3* | -0.942867154 | 0.000828924 | 0.02116581 |
|  | *9530091C08Rik* | -0.585278585 | 0.000858647 | 0.021678433 |
|  | *Slc40a1* | -0.712513838 | 0.000889339 | 0.022265671 |
|  | *Rhobtb1* | -0.513826103 | 0.000951375 | 0.023299623 |
|  | *Trib3* | -1.039412808 | 0.001016065 | 0.024223842 |
|  | *Zc3h3* | -0.454782267 | 0.00108204 | 0.025557221 |
|  | *Gng5* | -0.348642579 | 0.001118613 | 0.026182535 |
|  | *Tmigd3* | -0.687450862 | 0.001164997 | 0.027197375 |
|  | *Igfbp5* | -0.971168316 | 0.001190345 | 0.027717154 |
|  | *Sgsh* | -0.827468908 | 0.001256899 | 0.028892606 |
|  | *Arel1* | -0.726081386 | 0.001306094 | 0.029946878 |
|  | *Tfam* | -0.736119545 | 0.00131344 | 0.02996244 |
|  | *Banp* | -0.561128849 | 0.001355161 | 0.030603474 |
|  | *Naxd* | -0.500306218 | 0.001364218 | 0.030730808 |

**Table S9 Differential expression genes detected by edgeR (continued)**

| **Type** | **Gene symbol** | **log2FoldChange** | **P value** | **P adjust** |
| --- | --- | --- | --- | --- |
| Sex-biased genes in healthy | *Rammet* | -0.486069116 | 0.001483454 | 0.032840592 |
|  | *Wipi1* | -0.619499189 | 0.001497622 | 0.032911057 |
|  | *Rplp2* | -0.237813973 | 0.001539318 | 0.033662759 |
|  | *Tomm6* | -0.307421203 | 0.001587149 | 0.034594168 |
|  | *Pak4* | -0.591360884 | 0.001589607 | 0.034594168 |
|  | *Akr1c19* | -0.57193034 | 0.001704394 | 0.036648548 |
|  | *Csf2ra* | -0.652029411 | 0.001723377 | 0.03685217 |
|  | *Slc25a4* | -0.219995332 | 0.001733055 | 0.03685217 |
|  | *Rps4x* | -0.313210477 | 0.002062278 | 0.041282314 |
|  | *Ncoa6* | -0.54367624 | 0.002189661 | 0.043254235 |
|  | *Ggact* | -0.57497961 | 0.002201724 | 0.043302186 |
|  | *Fdft1* | -0.560416488 | 0.002266594 | 0.044480669 |
|  | *Dbt* | -0.690293306 | 0.002303298 | 0.045004431 |
|  | *Nfkbia* | -0.561406212 | 0.002385393 | 0.046406743 |
|  | *Gabbr2* | -0.720910134 | 0.002540945 | 0.048441022 |
|  | *Gramd3* | -0.375564882 | 0.002543854 | 0.048441022 |
|  | *Rack1* | -0.330049976 | 0.002600869 | 0.049143507 |
|  | *Prkar2a* | -0.858399515 | 0.002602616 | 0.049143507 |
|  | *Mapre3* | -0.408512976 | 0.002612671 | 0.049229947 |

*Note:* The value of log2FoldChange > 0 represents upregulation in male, and the value of log2FoldChange < 0 represents upregulation in female.
